# Single-cell transcriptome profiles of *Drosophila fruitless*-expressing neurons from both sexes

**DOI:** 10.1101/2022.03.09.483618

**Authors:** Colleen M Palmateer, Catherina Artikis, Savannah G Brovero, Benjamin Friedman, Alexis Gresham, Michelle N Arbeitman

## Abstract

*Drosophila melanogaster* reproductive behaviors are orchestrated by *fruitless* neurons. We performed single-cell RNA-sequencing on *fru P1* pupal neurons. Uniform Manifold Approximation and Projection (UMAP) clustering generates an atlas containing 113 clusters. While the male and female neurons overlap in UMAP space, more than half the UMAP clusters have sex-differences in neuron number, and nearly all clusters display sex-differential expression. Based on an examination of enriched marker genes, we annotate clusters as circadian clock neurons, mushroom body Kenyon cell neurons, neurotransmitter- and/or neuropeptide-producing, and those that express *doublesex*. Enriched marker gene analyses also shows that genes that encode members of the immunoglobulin superfamily of cell adhesion molecules, transcription factors, neuropeptides, neuropeptide receptors, and Wnts have unique patterns of enriched expression across the clusters. *In vivo* spatial gene-expression links to the UMAP clusters are provided. A functional analysis of *fru P1* circadian neurons shows they have dimorphic roles in sleep/activity and period length. Given that most clusters are comprised of male and female neurons indicates that the sexes have *fru P1* neurons with common gene expression programs. Sex-specific expression is overlaid on this program, to build the potential for vastly different sex-specific behaviors.

## Introduction

A current goal of neuroscience research is to generate cell atlases to molecularly classify cellular components of the nervous system, with the ultimate goal of understanding how the cells work together to generate a behavior (Ngai, 2022). The advent of single-cell RNA-sequencing (scRNA-seq) approaches have allowed researchers to get closer to this goal. *Drosophila* is an excellent model for this approach, given there are defined and experimentally tractable sets of neurons that generate the potential for complex behaviors. Indeed, a *Drosophila* scRNA-seq atlas has been generated to understand neurons underlying adult circadian biology, with the characterization of the adult core circadian clock neurons (Ma et al., 2021). In addition, there are atlases that have been generated to understand the cellular components of the brain and ventral nerve cord during development and adult stages, using single-cell genome-wide approaches (Allen et al., 2020; Croset et al., 2018; Davie et al., 2018; Konstantinides et al., 2018; Kurmangaliyev et al., 2020; Li et al., 2017; Li et al., 2022; McLaughlin et al., 2021; Ozel et al., 2021; Simon and Konstantinides, 2021; Xie et al., 2021). Here, we add to these cell atlas resources with a scRNA-seq study to understand how the potential for sexually dimorphic adult reproductive behaviors are specified in the developing nervous system in males and females. We gain insight into how neurons that arise from a sex-shared developmental trajectory can underlie vastly different behaviors in males and females (Ren et al., 2016).

In *Drosophila*, the neuronal substrates that direct reproductive behaviors are specified by the sex-specific transcription factors encoded by *fruitless* (*fru*; *fru P1* transcript isoforms) and *doublesex* (*dsx*) (**Figure 1A**), produced as an outcome of alternative pre-mRNA splicing by the sex determination hierarchy (reviewed in Andrew et al., 2019; Cline and Meyer, 1996). The identification of master regulatory transcription factors has provided an unprecedented molecular inroad into a behavioral question, allowing for high-resolution genomic and genetic interrogation, microscopic visualization, and the physiological manipulation of neurons directing behavior. *dsx* and *fru P1*-expressing neurons (*fru P1* neurons hereafter) are present in both sexes and each set arises from sex-shared developmental lineages (Ito et al., 1996; Lee et al., 2002; Manoli et al., 2005; Ren *et al*., 2016; Robinett et al., 2010; Ryner et al., 1996; Sanders and Arbeitman, 2008; Stockinger et al., 2005). However, these neurons direct dramatically different innate behaviors in the sexes due to differences in morphology, connectivity, physiology, and number. Males display an intricate courtship display that includes chasing the female, tapping her with his leg, and singing a song by wing vibration. The female will either accept or reject the male’s courtship advances. Once the female has mated, she shows a broad range of post-mating changes including changes in her receptivity to subsequent courtship displays (reviewed in Anholt et al., 2020; Auer and Benton, 2016; Dauwalder, 2011; Greenspan and Ferveur, 2000; Manoli et al., 2006; Peng et al., 2021; Villella and Hall, 2008; Yamamoto et al., 2014).

**Figure 1.**
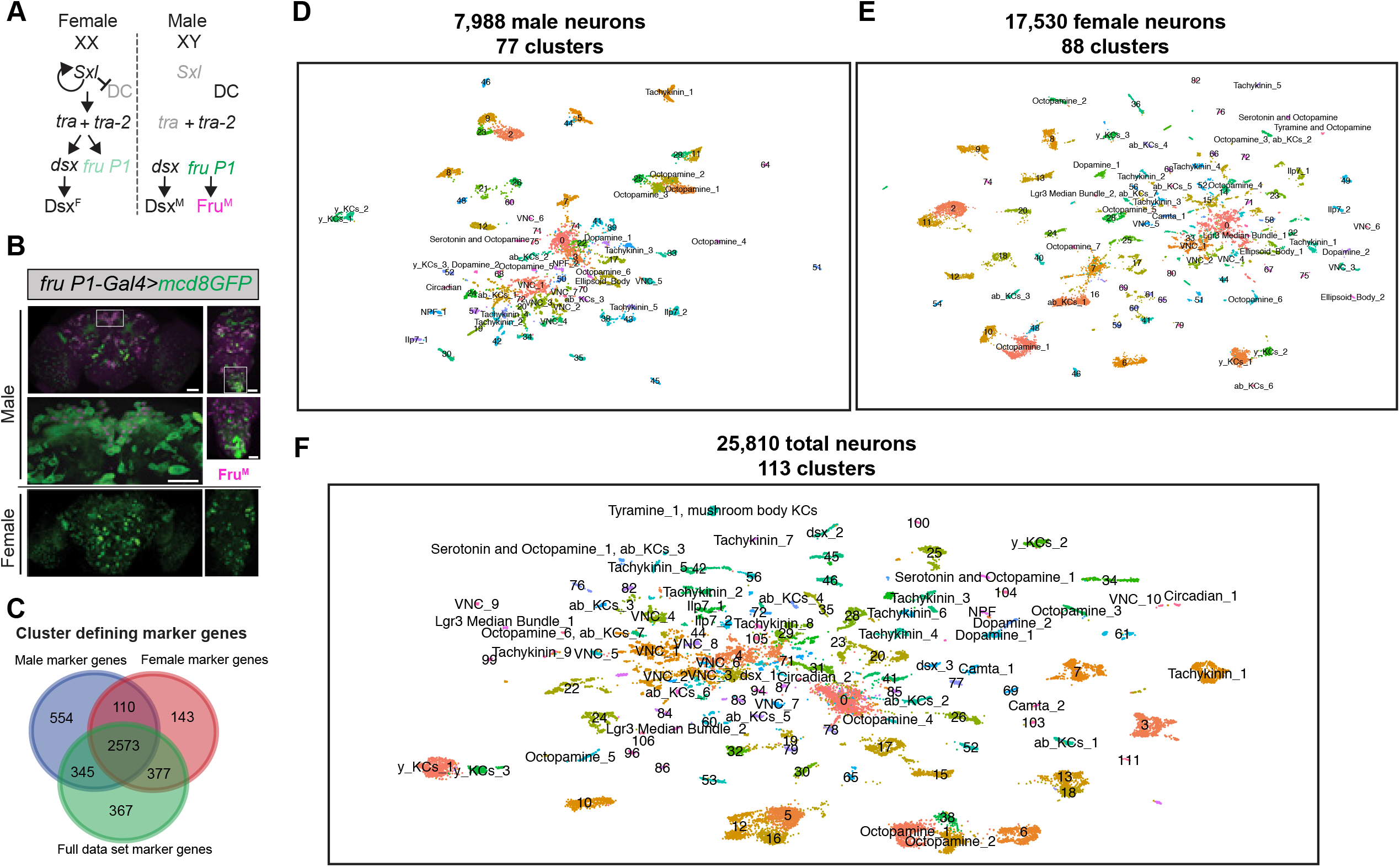
scRNA-sequencing of *fru P1* neurons from males and females at 48hr APF. **(A)** The *Drosophila* sex determination hierarchy is an alternative pre-mRNA splicing cascade that generates sex-specific transcription factors that regulate somatic sexual differentiation. The pre-mRNA splicing regulators are encoded by *sxl*, *tra* and *tra-2*. Early production of Sxl in females results in continued production of functional Sxl and Tra in females. Tra and Tra-2 regulate the sex-specific splicing of *doublesex* (*dsx*) and *fruitless* transcripts from the *P1* promoter (*fru P1*). In females, female-specific Dsx (Dsx^F^) is produced. In males, male-specific (Dsx^M^) and male-specific Fru (Fru^M^) are produced due to default pre-mRNA splicing. Sxl also regulates dosage compensation (DC) (reviewed in Andrew *et al*., 2019; Cline and Meyer, 1996) (**B**) Confocal maximum projections of 48hr APF male (top) and female (bottom) brains and ventral nerve cords (VNC) expressing membrane-bound GFP (*mcd8::GFP*) in *fru P1*-expressing neurons (green). Male tissues are also immunostained for male-specific Fru^M^ (magenta). Magnification of the boxed regions in male brain and VNC are below the male tissues (40x). Scale bars = 50 μm. (**C**) Venn diagram showing overlap of the full list of cluster marker genes from male (blue), female (red) and the full data set analyses (green). **(D)** UMAP plot of 7,988 *fru P1* cells, from male CNS at 48hr APF, grouped into 77 clusters (male data analysis). **(E)** UMAP plot of 17,530 *fru P1* cells, from female CNS at 48hr APF, grouped into 88 clusters (female data analysis). **(F)** UMAP plot of 25,530 total *fru P1* neurons, from both sexes (full data set), grouped into 113 clusters. For all UMAP plots, the annotations shown were determined using marker gene expression (**Supplemental Table 4)**. Clusters with numbers have additional annotations not shown and are listed in **Supplemental Table 4**.

Here, we focus on *fru P1* neurons that are found in both males and females, without large differences in their number or cell body positions (Manoli *et al*., 2005; Stockinger *et al*., 2005). *fru P1* was initially thought to be important for male courtship based on analyses of different *fru P1* mutant allele combinations (Anand et al., 2001; Gailey and Hall, 1988; Ryner *et al*., 1996; Villella et al., 1997). Additional studies showed that *fru P1* neurons are both necessary and largely sufficient for male courtship behaviors, based on experiments where *fru P1* neurons were either activated or silenced and behavior was examined, and by studies where male-specific Fru (Fru^M^) was produced in females in *fru P1* neurons (Clyne and Miesenbock, 2008; Demir and Dickson, 2005; Manoli *et al*., 2005; Stockinger *et al*., 2005). Female receptivity was later shown to be directed by *fru P1* neurons using neuronal silencing approaches (Kvitsiani and Dickson, 2006). *fru P1* neurons make up ∼2-5% of all neurons, in both sexes, with expression found in the periphery, brain and ventral nerve cord (Lee et al., 2000; Manoli *et al*., 2005; Stockinger *et al*., 2005). *fru P1* neurons in peripheral structures are important for detecting con-specific mates. In the central nervous system, *fru P1* is in regions important for higher-order processing of experience/sensation and regions involved in motor outputs for reproductive behaviors (reviewed in Auer and Benton, 2016)

The spatial positions of *fru P1* neurons in the brain and ventral nerve cord (VNC) have been assigned to named clusters, with the idea that they may share functions given their spatial proximity (Billeter and Goodwin, 2004; Lee *et al*., 2000; Manoli *et al*., 2005; Stockinger *et al*., 2005). However, it is not clear if all neurons within a cluster are functionally and/or molecularly similar, which we address here. While the position and number of neurons is similar between the sexes, there is dimorphism in cell number within several clusters, male-specific clusters, sex differences in neuronal projections, and differences in their physiology (reviewed in Auer and Benton, 2016; and Billeter and Goodwin, 2004; Cachero et al., 2010; Clowney et al., 2015; Datta et al., 2008; Kallman et al., 2015; Kimura, 2008; Kimura et al., 2005; Ruta et al., 2010; Yu et al., 2010). Sex differences in *fru P1* neurons begin to be established during metamorphosis, coinciding with when Fru^M^ is at its peak expression in males, during the mid-pupal phase (Arthur et al., 1998; Belote and Baker, 1987; Chen et al., 2021; Kimura *et al*., 2005; Lee *et al*., 2000; Stockinger *et al*., 2005). Therefore, a major remaining question is how does *fru* establish these sex differences and ultimately the potential for sex-specific behavior during development. This scRNA-seq study addresses what are the different molecular classes of *fru P1* neurons that arise during development in each sex.

Work from our lab and others have examined gene expression in *fru P1* neurons, Fru^M^ regulated expression, Fru^M^ binding, chromatin modifications in *fru P1* neurons, and open chromatin regions in *fru P1* neurons (Brovkina et al., 2021; Dalton et al., 2013; Goldman and Arbeitman, 2007; Neville et al., 2014; Newell et al., 2016; Palmateer et al., 2021; Vernes, 2014). Among these studies, only two have evaluated the mid-pupal stage (Neville *et al*., 2014; Palmateer *et al*., 2021). However, these studies used bulk sequencing approaches, thereby lacking the resolution to examine differences in gene expression across individual neurons. To determine the gene expression signatures for individual *fru P1* cells, we performed single-cell RNA-seq (scRNA-seq) during a critical developmental stage for establishing the potential for sex-specific behavior. This produced three cell atlases containing: *fru P1* male neurons, *fru P1* female neurons, and a full data set of male and female *fru P1* neurons co-analyzed. Here, we mainly focus on the full data set of 25,810 neurons from both males and females that formed 113 Uniform Manifold Approximation and Projection (UMAP) molecular clusters. We examine sex-differences in gene expression and neuron number within these clusters. We also manually annotated several clusters using the enriched gene expression signatures. We annotate clusters based on gene expression consistent with producing fast-acting neurotransmitters, neuropeptides, receptors, transcription factors, and several previously characterized *fru P1* neuronal populations. Additionally, we examine the *in vivo* spatial patterns of overlapping expression with *fru P1* and several marker genes to provide anatomical links to the UMAP space. Our identification of a population of neurons that express genes that regulate circadian behavior led to our discovery of a set of *fru P1* neurons that regulate sleep/activity/period length in a sex-specific manner. Altogether, our data sets show the molecular and functional heterogeneity of *fru P1* neurons, revealing the diversity of neurons required to build reproductive behaviors in males and females. While the data reveal sex-differences in gene expression within molecularly defined clusters, the observation that nearly all clusters are comprised of male and female neurons indicates that male and female *fru P1* neurons also share common gene expression repertoires, with sex-specific information overlaid on these core patterns.

## Results and Discussion

### scRNA-seq of *fru P1*-expressing cells in the pupal CNS reveals cell type diversity

To gain insight into the molecular and functional differences across *fru P1*-expressing neurons (*fru P1* neurons) in males and females, we performed scRNA-seq, using the 10x Genomics platform. This approach allows us to examine the transcriptomes of individual *fru P1* neurons. To enrich for *fru P1* neurons, fluorescence activated cell sorting (FACS) was performed on dissociated neurons from pupae CNS tissue, at 48hrs after puparium formation (APF), that expressed membrane-bound-GFP in *fru P1* neurons (*fru P1 > mCD8::GFP*) (Manoli *et al*., 2005) (**Figure 1B**). This is the pupal stage where Fru^M^ is at peak expression in males, based on immunostaining, and a critical period for when *fru P1* establishes the potential for behavior (Arthur *et al*., 1998; Belote and Baker, 1987; Chen *et al*., 2021; Lee *et al*., 2000). After 10x library construction, Illumina sequencing, read mapping and processing with the 10x Genomics CellRanger pipeline, we obtained gene-barcode matrices for further analyses.

These matrices were filtered to retain high quality cells: cells were removed that had >5% mitochondrial transcripts (dying cells), <200 genes detected (empty droplets), those expressing more than 4,000 genes (nfeatures), and/or possessing more than 20,000 unique molecular identifiers (UMIs; a metric for transcript counts) (potential doublets or triplets) (**Supplemental Figure 1**). After filtering, there were 7,988 male cells and 17,530 female cells in the pooled replicates (**Supplemental Figure 1**). The pooled replicates from males have 1,653 genes and 4,712 UMIs per cell (median values). The pooled replicates from females have 1,729 genes and 5,186 UMIs per cell (median values) (**Supplemental Table 1**). The combined filtered data from both sexes contains 25,810 *fru P1*-expressing cells (hereafter called the full data set). In the full data set, the cells have a median of 1,706 genes and 5,046 UMIs detected per cell, which is on par with or better coverage than other single cell atlas studies that examine the *Drosophila* CNS (Allen *et al*., 2020; Avalos et al., 2019; Croset *et al*., 2018; Davie *et al*., 2018; Li et al., 2021) (**Supplemental Table 1**). As expected, given that *fru P1* is expressed in neurons, 99% percent of the cells in the full data set are classified as neurons (25,509 cells). This is based on detecting expression of at least one of three genes with known neuronal expression (*embryonic lethal abnormal vision*, *elav*; *neuronal Synaptobrevin*, *nSyb*; *long non-coding RNA: noe*; **Supplemental Table 2**) (Diantonio et al., 1993; Kim et al., 1998; Robinow et al., 1988). Given this, the cells in the scRNA-seq dataset will be called neurons, hereafter.

We next scaled and normalized the expression data from the full data set, and also the male and female data sets separately, using a Seurat workflow (Stuart et al., 2019). To generate uniform manifold approximation and projection (UMAP) plots for visualization and analysis, we selected the number of significant principle components from the expression data, based on JackStraw analyses (Stuart *et al*., 2019) (**Supplemental Figure 2**). The male and female replicates show large overlap in UMAP space, indicating that there are minimal batch effects between replicates (**Supplemental Figure 1G-I**). To identify neurons that exhibit similarities in their gene expression profiles, we performed graph-based clustering analysis on each data set. To examine gene expression differences between the clusters in each data set, we performed an analysis to identify marker genes (**Figure 1C**). Marker genes are those that have significantly higher expression in one cluster compared to all other clusters within each analysis (**Supplemental Table 4**). A comparison of the complete list of marker genes from each analysis indicates that there are sex differences in the marker genes identified (**Figure 1C**). Therefore, we examine the UMAPs and clustering from each sex and the full data set going forward. The male data set has 77 clusters (**Figure 1D, Supplemental Figure 3A**), the female data set has 88 clusters (**Figure 1E, Supplemental Figure 3B**), and the full data set has 113 clusters (**Figure 1F, Supplemental Figure 4**). Nearly all clusters have cells from each replicate, further demonstrating that replicates show high concordance (**Supplemental Table 2**). For example, we found that 99% of clusters in the male data set and 100% of clusters in the female data set have cells from each replicate (**Supplemental Table 2**). Given that *fru P1* is expressed in ∼2000 neurons (Lee *et al*., 2000), it is possible we are examining the full repertoire of *fru P1* neurons (∼3.9X coverage for male cells and ∼8.8X coverage for female cells), assuming there are not biases in the ability to capture different *fru P1* neurons in the experimental workflow.

Next, we evaluate the molecular differences across the clusters by examining marker genes. There is a range in the number of marker genes for each cluster: 48-501 in the male data set, 74-555 in the female data set, and 61-610 in the full data set (**Supplemental Table 4**). In total, we find 3,582 marker genes in the male data set, 3,203 in the female data set, and 3,662 in the full data set (**Supplemental Table 4**). When we determine the overlap of the marker genes in the three analyses, the majority are shared (2,573 genes; **Figure 1C**), indicating that *fru P1* neurons from males and females have overlapping gene expression profiles (**Figure 1**), as we previously showed (Newell *et al*., 2016; Palmateer *et al*., 2021). However, each analysis reveals unique marker genes, with the most occurring in the male data set (**Figure 1C**). This is also consistent with our previous studies that demonstrated that there are more genes with male-biased expression in *fru P1*-expressing neurons, based on a cell-type-specific, bulk RNA-seq approach called Translating Ribosome Affinity Profiling (TRAP) (Newell *et al*., 2016; Palmateer *et al*., 2021; Thomas et al., 2012). When we examine the overlap in marker genes from the three scRNA-seq data sets with gene lists from our previous TRAP study that examined gene expression enriched in *fru P1* neurons at 48hr APF, we find significant overlap between the gene lists (**Supplemental Table 4**) (Palmateer *et al*., 2021). Gene expression heatmaps show the top five marker genes, based on log fold-change, for each cluster (**Supplemental Figure 4C-E**). The robust expression differences of the marker genes across clusters indicates that the clusters are different at the molecular level.

To understand the functions of neurons in the different clusters, we annotate the UMAP clusters, with a primary focus on the full data set (113 clusters, **Figure 1F**). To gain insight into which genes to focus on for annotation, we determined which gene functional groups are enriched in the marker genes lists, by performing Gene Ontology (GO) analyses. An examination of the top ten overrepresented terms for three GO categories: ‘molecular function’, ‘biological process’, and ‘cellular component’ (**Supplemental Figure 6A-C, Supplemental Table 5**), shows that they are shared across the three data set analyses (**Supplemental Figure 6A-C**), and are characteristic of developing neurons. In addition, we also performed protein domain and pathway enrichment analyses for these marker gene lists and find that the top enriched terms are also shared across the three data sets (**Supplemental Table 5**). We annotate each cluster by indicating the distribution of the marker genes from some of the enriched categories (GO, protein domain and pathways) and from a curated selection of genes that are biologically relevant (**Supplemental Table 4**). Additional annotation of the clusters is based on published expression data, gene ontology enrichments of marker genes from each cluster, and overlap with other single cell studies in the CNS (**Supplemental Table 4**) (Allen *et al*., 2020; Croset *et al*., 2018; Davie *et al*., 2018; Ma *et al*., 2021). We provide a name for 46 clusters and information at the level of marker gene expression for the remaining clusters.

### Examination of *fru P1* scRNA-seq clusters with sex-differences in neuron number

An examination of the full data set UMAP plot shows separation of male and female neurons within each cluster, with some clusters having more neurons from one sex (**Figure 2A, Supplemental Figure 1I**). This separation of the male and female cells is not due to the highly expressed male-specific long non-coding *RNA on the X* 1 and/or 2 (*roX1* and *roX2*), based on a full Seurat analysis performed when these genes are removed (**Supplemental Figure 5I-J**). We note that some of the separation may be due to developmental differences between males and females, given that females have a shorter pupal phase than males (Bakker and Nelissen, 1963). We next evaluated the clusters that have differences in neuron number between the sexes in our full data set. Given there are 2.19x more female neurons in the full data set we did several tests to determine the impact of this on clustering. After normalizing the number of neurons in each female cluster by dividing by 2.19x, we find that 52% of the clusters show a sex-bias in their neuron numbers. 24% show a >2-fold difference (sex-bias), 28% show >4-fold difference (strong sex-bias), and 3.5% of the clusters are sex-specific (**Figure 2B**, **Supplemental Table 3**). All references to sex-biased clusters in the full data set analysis are on normalized data values (**Supplemental Table 3)**.

**Figure 2.**
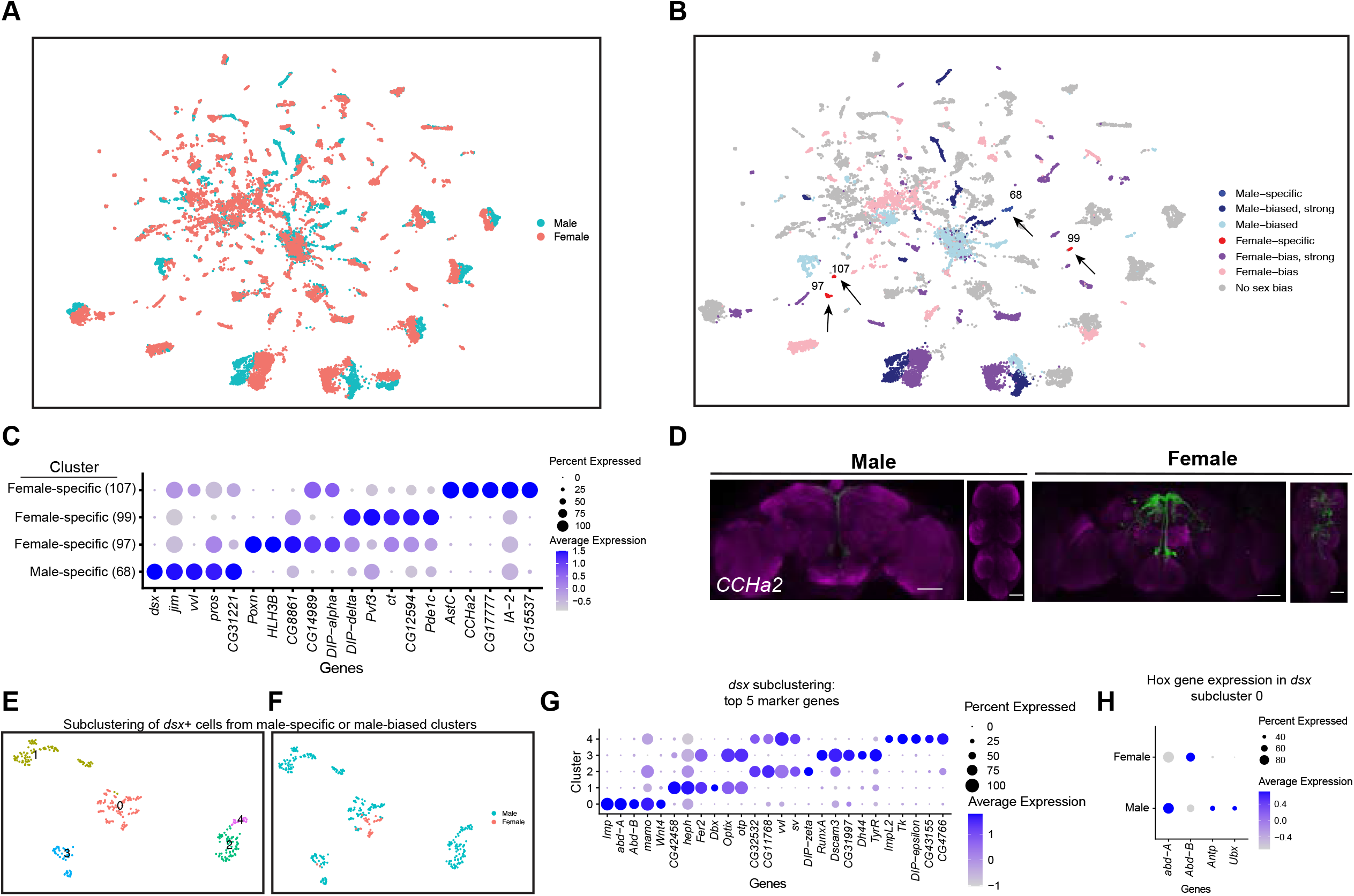
Sex differences in single-cell clustering. **(A-B)** UMAP plots of the 25,530 *fru P1* neurons from males and females (full data set) with all cells labeled by sex **(A)** and by sex-bias in cell number **(B)** (**Supplemental Table 3**). **(B)** Sex-biased classification is based on normalized number of female cells (legend on right, **Supplemental Table 3**). Cluster numbers and black arrows indicate female-specific clusters (red) and the male-specific clusters (dark blue). Female- (light pink) and male-biased (light blue) have 2-fold more neurons compared to the other sex. Female- (purple) and male-biased, strong (blue) have 4-fold more neurons compared to the other sex. **(C)** Dot plot of top 5 marker genes in each sex-specific cluster based on log fold-change in expression (**Supplemental Table 4**). **(D)** Brain and VNC confocal maximum projections from 0-24hr adults for *CCHa2* ∩ *fru P1*-expressing neurons, with intersecting expression shown in green for males (left) and females (right). Scale bars = 50 μm. (**E-F**) UMAP subcluster containing neurons from male-specific and male-biased clusters that express *dsx*, defining five subclusters (clusters are indicated by color and number). (**F**) The five subclusters are shown with sex indicated by color, with male cells in blue and female cells in pink. Female cells are present only in subclusters 0 and 3. **(G)** Dot plot of top 5 marker genes in each *dsx* subcluster based on log fold-change in expression (**Supplemental Table 4**). (**H**) Dot plot of Hox genes with known VNC expression in *dsx* subcluster 0, for both sexes. For all dot plot (**C, G and H**) panels the dot size indicates the percentage of neurons expressing each gene per cluster (Percent Expressed). Average normalized expression level is shown by color intensity (Average Expression).

We next determined if this difference is maintained when equal numbers of neurons between the sexes are present in the UMAP, by removing random subsets of female neurons to equal the number of male neurons, which was performed three independent times. We find that six clusters change their status after random removal across some of the independent cell removal analyses (**Supplemental Figure 7A, Supplemental Table 3**). Further, we repeated the full Seurat workflow by first removing female cells at random (downsampling) and generating new UMAP plots and clustering analysis for three independent downsampled analysis, with equal numbers of male and female cells. To examine the similarity of gene expression between the clusters, we use Pearson correlations in clustifyr (Fu et al., 2020). We match 80%-85% of the clusters in the downsampled analyses to those found in the full data set, and of those there is high concordance of sex-specific and sex-biased clusters (**Supplemental Figure 7B, Supplemental Table 3**). However, fewer female-specific clusters are present in the downsampled analyses and three additional male-specific clusters are resolved (**Supplemental Figure 7B, Supplemental Table 3**). Though differences are revealed in the analyses when female neurons are randomly removed to equal the number of male cells, most sex-biased clusters are preserved. Taken together, we proceed with the analysis of the UMAP clusters in the full data set, as the additional female cells provide information.

In the full data set there are four sex-specific clusters (three female-specific and one male-specific), with two maintained after the downsampling analysis (female-specific cluster 107 and male-specific cluster 68) (**Figure 2B,** arrows). An examination of the top five marker genes for the female-specific clusters that are not maintained across UMAP analyses (cluster 97 and 99) shows that each have a *Dpr Interacting Protein* (*DIP*) gene in the list (*DIP-alpha* or *DIP-delta*) (**Supplemental Table 4**). The *DIP*s encode members of the Immunoglobulin superfamily (IgSF), a family that is highly enriched among the full marker gene list (**Supplemental Table 5**). Previous studies have shown that the IgSFs are regulated by Fru^M^ and/or have enriched expression in *fru P1* neurons (Brovkina *et al*., 2021; Dalton *et al*., 2013; Goldman and Arbeitman, 2007; Neville *et al*., 2014; Newell *et al*., 2016; Palmateer *et al*., 2021; Vernes, 2014). Additionally, our analysis of adult *dpr*/*DIP*-expressing neurons that also express *fru P1* (*DIP* ∩ *fru P1*), showed that *DIP-alpha* ∩ *fru P1* and *DIP-delta* ∩ *fru P1* neurons have region-specific dimorphism in CNS neuron number and projection pattern (Brovero et al., 2021).

One of the top marker genes for the female-specific cluster 107 is *CCHamide-2* (*CCHa2*), which encodes a neuropeptide hormone that stimulates feeding (Ren et al., 2015). Cluster 107 neurons may be from the VNC, given that cluster 107 marker genes include Hox segment identity genes *Antennapedia* (*Antp*) and *Ultrabithorax* (*Ubx*). *Antp* and *Ubx* define VNC T1-T3 and T2-T3 thoracic segments, respectively (Baek et al., 2013; Jarvis et al., 2012). To determine if there are female-specific *CCHa2* neurons that also express *fru P1* (*CCHa2* ∩ *fru P1*) in the VNC, we used a genetic intersectional approach to visualize *CCHa2* ∩ *fru P1-*expressing neurons. The approach relies on the two-component Gal4/UAS system that drives expression of a Myc-tagged membrane-bound reporter protein (Nern et al., 2015). Expression of the UAS-reporter requires Flippase (FLP)-mediated removal of a stop cassette. The expression of FLP is directed by *fru P1* regulatory elements (Yu *et al*., 2010). Therefore, reporter gene expression is limited to *fru P1* neurons that also express Gal4. Throughout this study, we examine expression at 48 hr APF and 0-24 hour adults, given that the intersectional approach might be slow to report on expression given the requirement of a genomic FLP-mediated recombination event and production of several proteins. Additionally, *fru P1* expression is most thoroughly characterized in adults (Billeter and Goodwin, 2004; Cachero *et al*., 2010; Kimura *et al*., 2005; Lee *et al*., 2000; Manoli *et al*., 2005; Stockinger *et al*., 2005; Yu *et al*., 2010). *CCHa2* is also a marker gene for additional clusters, so we expect the expression pattern to include neurons not in cluster 107. There is no detectable reporter gene expression at 48hr APF. In 1-day old adults, we find female-specific *CCHa2* ∩ *fru P1-*expressing neurons in both the brain and VNC, and *CCHa2* ∩ *fru P1-*expressing neurons in the median bundle in both sexes (**Figure 2D**). There are no male *CCHa2* ∩ *fru P1-*expressing neurons in the VNC. This observation indicates that there are female-specific *CCHa2* ∩ *fru P1* neurons in the VNC, as suggested by the UMAP cluster analysis.

The top marker gene for the male-specific cluster 68 is the sex hierarchy gene *dsx*. This is consistent with the observation that there are male-specific populations of *dsx* ∩ *fru P1* neurons in the brain (pC1, pC2, SN, and SLN) and VNC (TN1 and abdominal ganglion) (Billeter et al., 2006; Ishii et al., 2020; Lee *et al*., 2002; Rideout et al., 2007; Sanders and Arbeitman, 2008). Given that cluster 68 does not express Hox segment identity genes that are expressed in the VNC, or other genes with known VNC expression, these neurons are likely from the brain (**Supplemental Figure 8B**). There are two additional clusters in the full data set that have *dsx* as a marker gene: one strongly male-biased (4x more male cells after normalization) and one male-biased cluster (2x more male cells after normalization) (**Supplemental Figure 8C, Supplemental Table 3, Supplemental Table 4**). To analyze these clusters more deeply, we subclustered these neurons, selecting for those with detectable *dsx* expression and identified marker genes for each subcluster (**Figure 2B**, **Supplemental Figure 8C**). This reanalysis resolved five subclusters, retaining the original male-biased cluster as one cluster (subcluster 0), and generating two clusters from the original male-specific cluster (subclusters 2 and 4) and two clusters from the original strongly male-biased cluster (subclusters 1 and 3; **Figure 2E-F**, **Supplemental Figure 8D**).

We propose that the male-biased subcluster 0 contains neurons from the VNC, given these neurons also express *Antp, Ubx, abdominal A* (*abd-A*), and *Abdominal B* (*Abd-B*), (**Figure 2E-F**, **Supplemental Figure 8E-H, Supplemental Table 4**), and the most significant Berkeley Drosophila Genome Project (BDGP) enrichment term for the marker genes is ‘ventral nerve cord’, based on analysis in Flymine portal (Lyne et al., 2007). In the VNC, both males and females have *dsx* ∩ *fru P1* neurons in the abdominal ganglion, and there are male-specific *dsx* ∩ *fru P1* neurons in the metathoracic ganglion called TN1 and TN2 neurons (Billeter *et al*., 2006; Lee *et al*., 2002; Rideout *et al*., 2007; Sanders and Arbeitman, 2008). When we examine the expression of Hox genes in subcluster 0, many of the male and female neurons show expression of *abd-A* and *Abd-B* (**Figure 2H**). Given this, we propose these neurons are the *dsx* ∩ *fru P1* neurons in the abdominal ganglion. There are also neurons in subcluster 0 only in males, with expression of *Antp* and *Ubx* (**Figure 2H**) that we propose are the male-specific metathoracic TN1 or TN2 neurons. Consistent with this, we previously showed TNI and TN2 neurons underwent cell death in females by 48hr APF (Sanders and Arbeitman, 2008).

We propose that the strongly male-biased and male-specific *dsx* and *fru P1*-expressing neuron clusters contain neurons from the brain, based on expression of Hox genes (**Supplemental Figure 8E-H**). Previous studies have shown *dsx* ∩ *fru P1* neurons in the brain in PC1 and PC2 regions, as well as small sets of neurons called SN and SLN (Lee *et al*., 2002; Rideout *et al*., 2007; Sanders and Arbeitman, 2008). The strongly male-biased cluster split into two subclusters, one that has two female neurons (subcluster 3), and the other is now male-specific (subcluster 1). The male-specific cluster split into subclusters 2 and 4 that are positioned closely together in the UMAP space (**Supplemental Figure 8D**). A recent study using genetic intersectional approaches found 1-2 pC1/pC2 female *dsx* ∩ *fru P1* neurons (Ishii *et al*., 2020), whereas we did not detect any (Sanders and Arbeitman, 2008), leaving open that possibility that neurons in subclusters 3 are from either pC1 or pC2 brain regions, given the detection of female cells. Also, it has been proposed that male pC2 *dsx*-expressing neurons are comprised of distinct populations, based on fasciculation patterns from either medial and lateral positions (pC2 renamed pC2m and pC2l) (Robinett *et al*., 2010). Thus, subclusters 1, 2, 3 and/or 4 may contain neurons that are from the pC1 or pC2, SN, SLN *dsx*-expressing regions in brain, given there are no known molecular markers that would distinguish these populations and there are predicted to be several distinct populations based on morphological and spatial features. An examination of marker genes unique to each subclusters suggests the following differences in functions (pathway enrichment, Benjamini Hochberg p < 0.1; **Supplemental Table 5**). Subcluster 1 is distinct because of unique marker genes enriched in the pathway ‘Acetylcholine binding and downstream events’. Subcluster 2 is distinct because of unique marker genes enriched in the pathways ‘Signal Transduction’, ‘Calcineurin activates NFAT’, ‘GPCR ligand binding’, ‘Amine ligand-binding receptors’, and ‘Class A/1 receptors’. Subcluster 3 did not have enough unique genes to detect pathway enrichments. Subcluster 4 is distinct because of unique marker genes enriched in the pathway ‘Neuronal Systems’ (**Supplemental Table 5**).

### Hox gene expression identifies *fru P1* neurons from the VNC

To annotate clusters from the VNC, we examined expression of Hox genes *Antp*, *Ubx, Abd-A*, and *abd-B* that are known to be expressed in an anterior to posterior pattern in the VNC (Baek *et al*., 2013). These genes show restricted expression in a subset of UMAP clusters (**Figure 3A**, **Supplemental Figure 9A-H)**. Additionally, we find several have more than one of these Hox genes as a marker gene (**Supplemental Table 4**). For example, male data set cluster 1 has all four genes as markers, which indicates that neurons from several thoracic and abdominal VNC segments may be present in some UMAP clusters. This also indicates that neurons from different segments may have shared gene expression profiles. Therefore, any cluster with one or multiple Hox marker genes was annotated as a “VNC” cluster (**Supplemental Table 4**), resulting in seven VNC clusters in both the male and female data set, and 11 in the full data set (**Figure 1D, F, G**, **Figure 3B**, **Supplemental Figure 9I-J**, **Supplemental Table 4**). In addition, we performed a correlation analysis to examine co-expression of *Antp*, *Ubx, Abd-A*, and *abd-B* within single neurons. All the data sets have significant positive correlations for expression of these genes, with the highest correlations found between *Abd-A*, and *abd-B,* consistent with these genes have the strongest overlap in their spatial expression (Baek *et al*., 2013) (**Figure 3C, Supplemental Figure 9K-L**). This finding differs from scRNA-seq on the adult VNC, which showed significant anti-correlations in expression between all pairings, with the exception of *Abd-A*, and *abd-B* (Allen *et al*., 2020). This suggests that during development there is a refinement of Hox gene expression that limits co-expression in VNC neurons.

**Figure 3.**
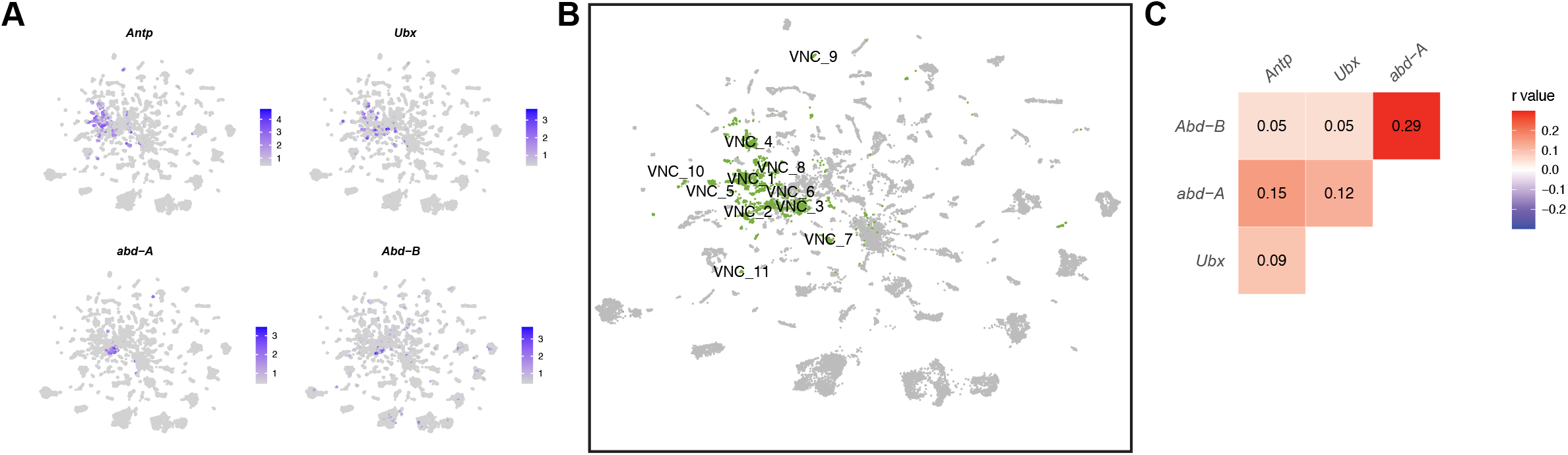
VNC cluster annotations based on *Hox* gene expression. **(A)** Gene expression feature plots showing neurons that express *Hox* genes with known expression in the VNC (*Antp*, *Ubx*, *abd-A*, and *Abd-B*), in the full data set UMAP. The gene expression feature plots show gene expression levels in purple in the UMAP, with color intensity proportional to the log normalized expression levels. **(B)** UMAP plot showing annotated VNC clusters in the full data set analysis. The VNC clusters are numbered and highlighted in green (**Supplemental Table 4**). **(C)** Heatmap of correlation of *Hox* gene expression in single neurons in the full data set. Pearson’s r values denoted on heatmap with color, according to legend (right).

### Identification of *fru P1* mushroom body Kenyon cell neuron populations

The mushroom body is a structure in the brain that is critical for *Drosophila* learning and memory (reviewed in Davis, 1993; Heisenberg, 2003). The mushroom body is comprised of intrinsic neurons called the Kenyon Cells (KCs), as well as extrinsic input and output neurons (Aso et al., 2014; Li et al., 2020). *fru P1* is expressed in the mushroom body γ and αβ KCs, whereas there are no reports of expression in the α’β’ KCs. Furthermore, previous work has demonstrated that when *fru P1* function is reduced by RNAi in γ or αβ Kenyon Cells (KCs) there are deficits in courtship learning (Manoli *et al*., 2005). To gain insight into *fru P1* MB KCs, we annotated clusters that are KC subtypes, based on marker gene analysis, using criteria similar to a previous scRNA-seq study of the *Drosophila* midbrain (Croset *et al*., 2018) (**Supplemental Table 4**). The marker gene criteria are schematically summarized (**Supplemental Figure 10A)**. Clusters that contain KCs can be identified based on having *eyeless* (*ey*) and/or *Dopamine 1-like receptor 2* (*Dop1R2*, also known as *damb*) as marker genes (Han et al., 1996; Kurusu et al., 2000)(**Supplemental Table 4**). To identify clusters that contain αβ KCs subtypes, we additionally required that *short neuropeptide precursor F* (*sNPF*), and/or *Fasciclin 2* (*Fas2*) are marker genes (Cheng et al., 2001; Crittenden et al., 1998; Johard et al., 2008). This identified one αβ KC cluster in male, four in female, and three in the full data set analyses (**Supplemental Table 4**). The clusters that contain γ KCs were identified as those that have *trio* as a marker gene, in addition to the marker genes described for identifying αβ KCs. *trio* encodes a Rho guanine nucleotide exchange factor that has high expression restricted to the γ KCs, at 48hrs APF (Awasaki et al., 2000). This allowed us to confidently annotate two γ KC clusters in each data set analysis (**Supplemental Table 4**). We also annotate potential γ KCs and αβ KC clusters based on having marker gene combinations consistent with containing KCs (indicated by *; **Supplemental Table 4)**. For example, we find one potential γ KC cluster in the female data set analysis that has only *Dop1R2* and *trio* marker genes. We also find several potential αβ KC clusters where *sNPF* and *Fas2* are marker genes, and there is expression of *ey* and/or *Dop1R2*, but these are not marker genes (see methods, **Supplemental Figure 10**). We could not confidently identify clusters containing *fru P1*-expressing α′β′ KCs, if they exist, as there are no known marker gene patterns that would distinguish this population at 48hr APF.

As several studies have implicated the αβ and γ lobes as being important for courtship memory in males, we next wanted to further examine the six high confidence KC clusters identified in the full data set analysis by subclustering and assessing marker genes. This resulted in 13 subclusters (**Figure 4B**), where the original three αβ KC clusters separate into five subclusters and the two γ KC clusters separate into seven subclusters (**Supplemental Figure 12J**). When we examine these subclusters (**Figure 4C**), one male-specific and one female-specific subcluster of γ KCs are identified (subclusters 4 and 12, respectively). The significant marker genes for male-specific subcluster 4 are enriched for ‘biological function’ GO enrichment terms ‘learning or memory’, ‘cognition’, ‘memory’ and ‘behavior’ (**Supplemental Table 5, Supplemental Table 6**). The list of seven significant marker genes for the female-specific cluster 12 do not exhibit any GO enrichment. Interestingly, among the seven marker genes for cluster 12, two are odorant binding protein encoding genes, *Odorant-binding protein 99a* (*Obp99a*) and *Odorant-binding protein 44a* (*Obp44a*) (**Figure 4D**, **Supplemental Table 6**), which have previously been shown to be expressed in the mushroom body of adult females (Crocker et al., 2016). Across the subclusters, we see overlap in expression of the top five maker genes (top based on log-fold-change, **Figure 4D**). For example, *Dopamine 1-like receptor 1* (*Dop1R1*) is a top marker gene in 11 out of 13 subclusters and has previously been shown to be highly enriched in KC populations (Croset *et al*., 2018). Other broadly expressed marker genes across *fru P1* KC subclusters include *Ribosomal protein L35* (*RpL35*), *IA-2 protein tyrosine phosphatase* (*IA-2*) and the *GFP* transgene we expressed to FACS sort *fru P1* neurons (**Figure 4D)**. We also identify subcluster-specific gene expression. Subcluster 11, an αβ KC subcluster, shows high expression of *Vmat* and *SerT*, suggesting that this population of *fru P1* αβ KCs is serotonergic (**Figure 4D)**. Altogether, this analysis suggests the existence of sex-specific *fru P1* KC subtypes that might have a role in learning during reproductive behaviors.

**Figure 4.**
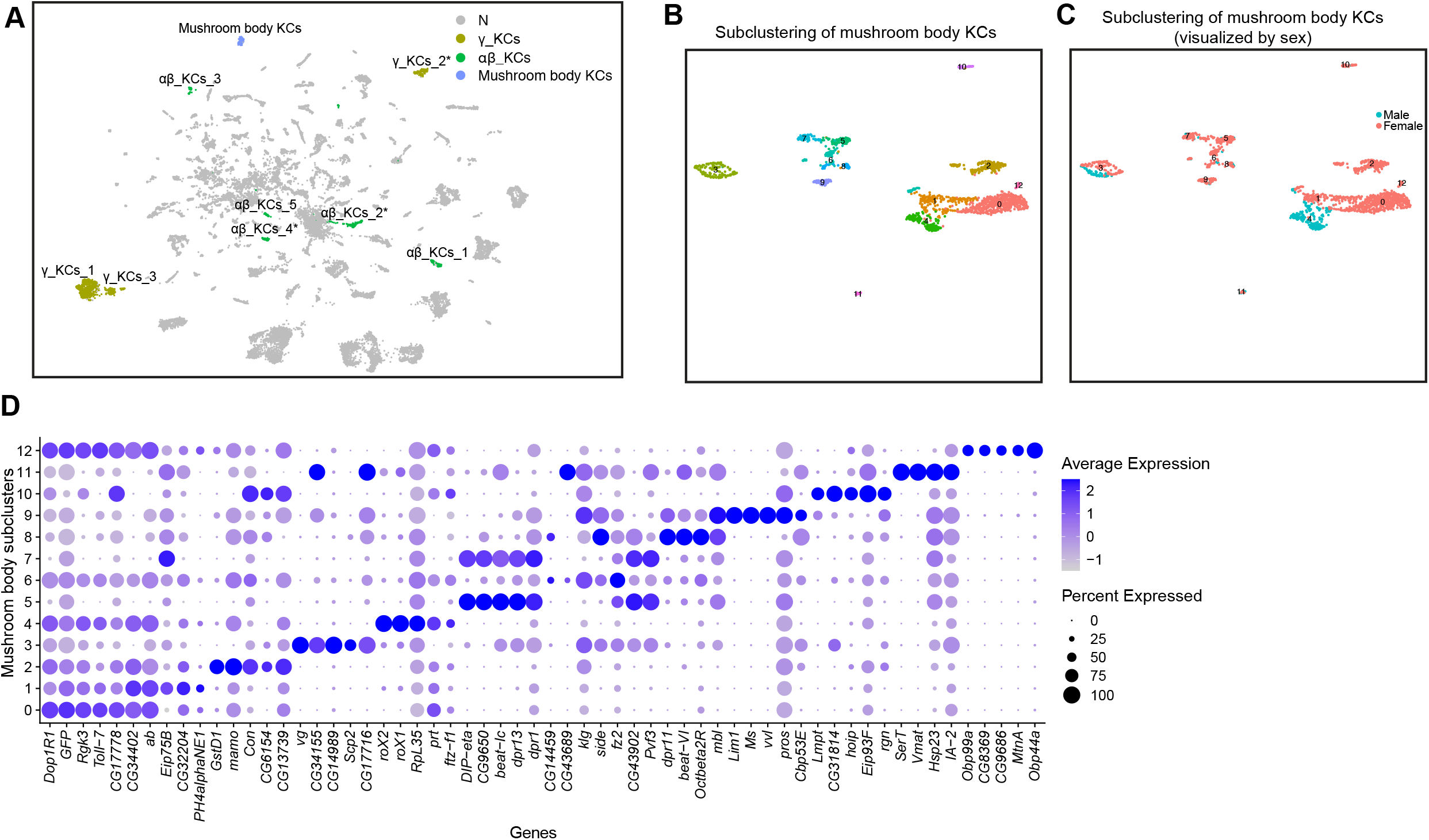
Annotation of mushroom body Kenyon cells. **(A)** UMAP plot highlighting Kenyon cell (KC) cluster annotations in the full data set analysis (**Supplemental Table 4**). N (grey) indicates clusters not meeting marker gene criteria used to annotate KC clusters (see methods and **Supplemental Figure 10A**). **(B)** UMAP of subclustering analysis of neurons from mushroom body KC clusters (colored clusters from panel **A**), creating 13 subclusters. (**C**) The 13 subclusters are shown with sex indicated by color, with male cells in blue and female cells in pink. (**D**) Dot plot of top 5 marker genes in each mushroom body KC subcluster based on log fold-change in expression (**Supplemental Table 4**). Dot size indicates the percentage of neurons expressing each gene per cluster (Percent Expressed). Average normalized expression level is shown by color intensity (Average Expression).

### Identification of *fru P1* neurons that produce fast-acting neurotransmitters

We next identify the *fru P1* neuronal clusters that produce the fast-acting neurotransmitters (FAN) acetylcholine, glutamate, and GABA. To annotate clusters, we used marker genes that encode products in a FAN biosynthetic pathway or encode transporters for the neurotransmitters, as previously done (Allen *et al*., 2020; Avalos *et al*., 2019; Croset *et al*., 2018; Davie *et al*., 2018). *Vesicular acetylcholine transporter* (*VAChT*) and *Choline acetyltransferase* (*ChAT*) were used to identify cholinergic neurons. *Vesicular glutamate transporter* (*VGlut*) was used to identify glutamatergic neurons. *Glutamic acid decarboxylase 1* (*Gad1*) and *Vesicular GABA Transporter* (*VGAT*) were used to identify GABAergic neurons (**Figure 5A**, **Supplemental Figure 11A and H**). The majority of UMAP clusters that have neurons that produce FANs are annotated as cholinergic. For example, in the full data set analysis there are 32 cholinergic clusters, 18 GABAergic, and 16 glutamatergic (**Figure 5A**, **Supplemental Table 4**). Only one full data set UMAP cluster has marker gene expression for more than one FAN (cholinergic and glutamatergic), and the male and female data set each have one UMAP cluster that has marker gene expression for more than one FAN (GABAergic and glutamatergic) (**Figure 5A**, **Supplemental Figure 11A and H, Supplemental Table 4**).

**Figure 5.**
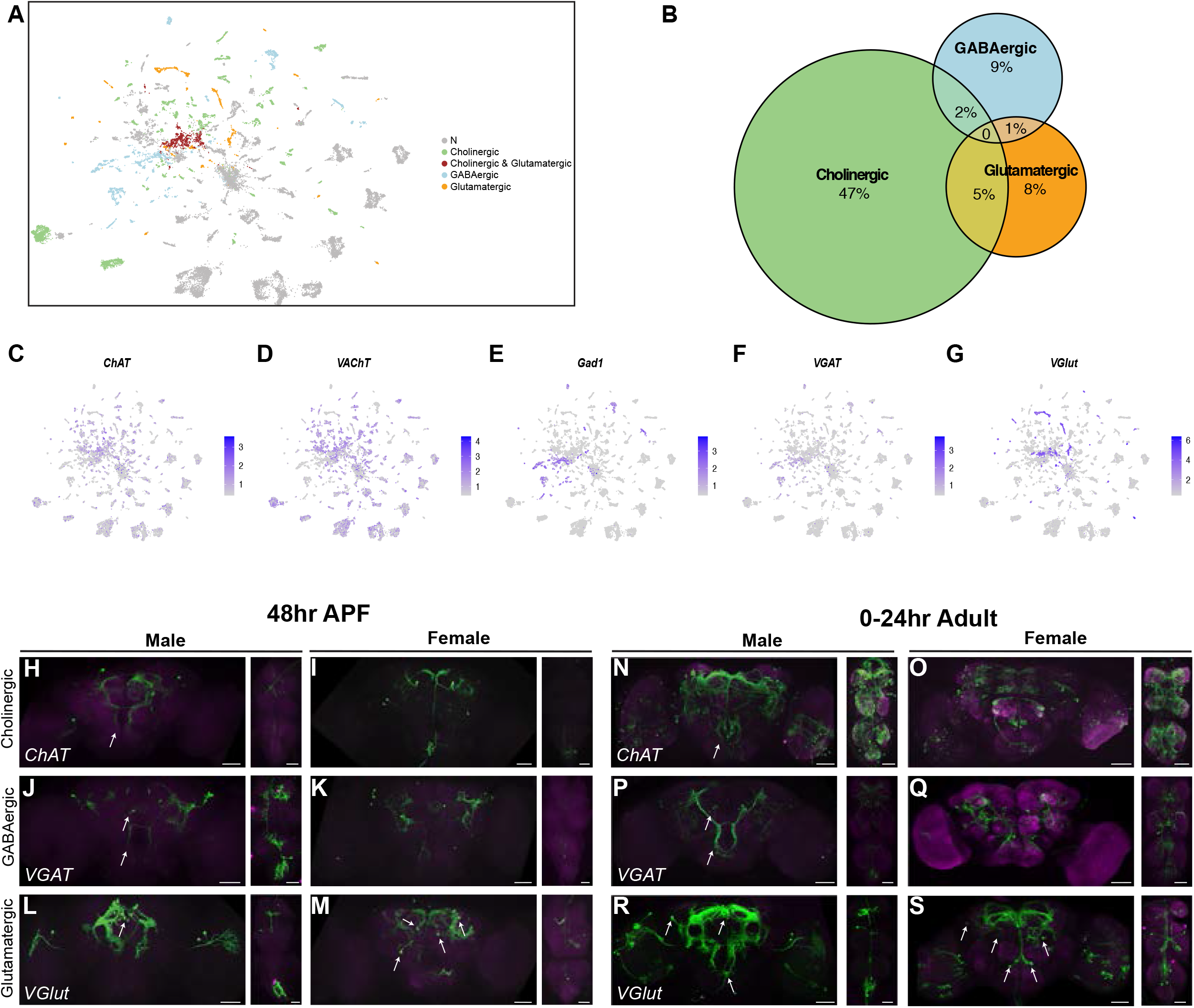
Annotation of neurons that produce fast-acting neurotransmitters (FANs). **(A)** UMAP plot showing annotated clusters with neurons that produce FANs, in the full data set analysis (**Supplemental Table 4**). In the color legend on right, N (grey) indicate clusters with no FAN marker gene expression. **(B)** Size proportional Euler diagram showing the percentage of cells with overlapping expression of genes indicative of cholinergic (*VAChT*), GABAergic (*Gad1*), and glutamatergic (*VGlut*) neurons, in the full data set. **(C-G)** Gene expression feature plots showing neurons that express genes indicative of FAN production: acetylcholine (*ChAT* and *VAChT*), GABA (*Gad1* and *VGAT*), and glutamate (*VGlut*), in the full data set UMAP. **(H-S)** Brain and VNC confocal maximum projections for FAN-producing ∩ *fru P1*-expressing neurons, with intersecting expression shown in green for males and females, as indicated. *ChAT* ∩ *fru P1* neurons*, VGAT* ∩ *fru P1* neurons, and *VGlut* ∩ *fru P1* neurons in 48hr APF and 0-24 hour adult are shown. Arrows point to regions with sexually dimorphic projects or cell bodies. **(T)** Dot plot of *prospero* (*pros*) expression in glutamatergic clusters from the full data set (**Supplemental Table 4**). Dot size indicates the percentage of neurons expressing each gene per cluster (Percent Expressed). Average normalized expression level is shown by color intensity (Average Expression). **(U)** Maximum confocal projection of male brain showing *VGlut* ∩ *fru P1* neurons (green) and anti-Prospero staining (magenta), imaged with ×20 objective. (**U’**) The Pars Intercerebralis (PI) reigon of the brain in white box in **U,** imaged with a ×40 objective. White arrows point of *VGlut* ∩ *fru P1* cell bodies in PI that do not show Prospero staining. Scale bars = 50 μm.

To determine if individual neurons express genes that would be indicative of a neuron producing multiple FANs, we examined expression of *VAChT* (cholinergic), *Gad1* (GABAergic), and *VGlut* (glutamatergic) at the neuron level, in each data set (**Figure 5B**, **Supplemental Figure 11B and I**). Similar to other single cell studies in the *Drosophila* CNS, this analysis revealed that *fru P1* neuronal populations largely produce only one FAN and therefore are mostly exclusive (**Figure 5B**, **Supplemental Figure 11B and I**) (Allen *et al*., 2020; Avalos *et al*., 2019; Croset *et al*., 2018). In contrast to these other scRNA-seq studies, no neurons in our data set express genes indicative of production of three FANs, and ≤8% of neurons co-express two genes indicative of production of two FANs (Allen *et al*., 2020; Avalos *et al*., 2019; Croset *et al*., 2018). The expression of these FAN biosynthetic pathway or transporter genes at the neuron level are also shown (**Figure 5C-G, Supplemental Figure 11C-G, J-N**), indicating that 62% are cholinergic (*VAChT* and/or *ChAT*), 19% are GABAergic (*Gad1* and/or *VGAT*), and 14% are glutamatergic (*VGlut*) (**Supplemental Table 2**).

To visualize the spatial patterns of *fru P1* neurons that produce different FANs we use a genetic intersectional strategy that relies on Gal4 transgenes that report on *ChAT* (cholinergic), *VGAT* (GABAergic), and *VGlut* (glutamatergic) (Deng et al., 2019; Nern *et al*., 2015; Yu *et al*., 2010). We can detect the three different *fru P1*-expressing FAN classes at 48hr APF in both sexes, based on reporter gene expression that labels the neuronal membrane of each FAN ∩ *fru P1*-expressing neuronal population. However, we do not find broad expression of *ChAT* at 48hr APF, which is not consistent with our single cell data that shows extensive expression of *ChAT* (**Figure 5C, Supplemental Figure 11C and J**). Examination of *ChAT* ∩ *fru P1*-expressing neuronal populations in 0-24 hour adults does reveal broader expression of *ChAT* compared to *VGAT* and *VGlut* (**Figure 5N-S**).

We find sexual dimorphisms in the morphology of *fru P1* neurons in the FAN ∩ *fru P1*-expressing neuronal populations, some that have been previously described (**Figure 5H-S**) (Cachero *et al*., 2010; Kimura *et al*., 2005; Yu *et al*., 2010). For example, in *CHAT* ∩ *fru P1* neurons, we observe male-specific expression in the tritocerebral loop (Yu *et al*., 2010). In *VGAT* ∩ *fru P1* neurons, we observe the sexually dimorphic mAL neurons, which have previously been shown to be GABAergic and suppress male courtship behavior in adults (**Figure 5J and P**) (Kallman *et al*., 2015; Kimura *et al*., 2005; Koganezawa et al., 2010). In addition, we find that the *VGlut* ∩ *fru P1* expression patterns exhibit sex differences in cell body positioning (**Figure 5L-M, R-S**). In the male brain, the majority of *VGlut* ∩ *fru P1* cell bodies are exclusively positioned near the Pars intercerebralis (PI), whereas in females, the cell bodies are additionally positioned throughout the midbrain (**Figure 5L-M, R-S,** white arrows).

There are 17 *VGlut* ∩ *fru P1* UMAP clusters, without a corresponding number of spatially distinct *VGlut* ∩ *fru P1* in the CNS in males and females (**Figure 5A, L-M, R-S**, **Supplemental Table 4**). This suggests that neurons that reside in similar spatial position may have different molecular identities, as has previously been found in other single cell studies of the *Drosophila* CNS (Ma *et al*., 2021). To interrogate gene expression differences, we further examined the top marker genes for these clusters. We found that *prospero* (*pros*), a gene which encodes a homeobox transcription factor protein that promotes neural differentiation (Doe et al., 1991), is a marker gene in three clusters (**Supplemental Table 4**). When we examined *pros* expression across all neurons in each glutamatergic UMAP cluster we find five additional clusters where *pros* is not a marker gene but is expressed in a high percentage of cells (**Figure 5T**). This analysis also showed that eight clusters exhibit little to no *pros* expression (**Figure 5T**). Next, we determined if neurons that are spatially close together are different at the molecular level, by examining Pros protein expression. In males, *VGlut* ∩ *fru P1* brain neurons are close together spatially, with the majority of the cell bodies positioned together near the Pars Intercerebralis region of the brain. These neurons are different molecularly, as only some have nuclear Pros staining, based on immunofluorescence analyses (**Figure 5U’**, white arrows, **Supplemental Figure 11O-S**). Earlier in development, *pros* has a role in neuroblast differentiation (Doe *et al*., 1991). However, by 48hr APF only mushroom body Kenyon cell neuroblasts are thought to be dividing (Ito and Hotta, 1992). The *VGlut* ∩ *fru P1* neurons are not KC neurons, leaving open the role of *pros* in the *fru P1* neurons evaluated here. Overall, the UMAP cluster analyses may reveal the molecular heterogeneity in closely positioned neurons in the brain, especially in the well-defined spatial clusters that the field has defined (Lee *et al*., 2000; Manoli *et al*., 2005; Stockinger *et al*., 2005).

### Aminergic populations of *fru P1* neurons

Subsets of *fru P1* neurons produce the biogenic amines dopamine, serotonin, tyramine, and octopamine (Billeter and Goodwin, 2004; Certel et al., 2010; Certel et al., 2007; Jois et al., 2018; Lee and Hall, 2001; Lee et al., 2001; Yu *et al*., 2010; Zhang et al., 2016). These aminergic *fru P1* neurons have roles regulating male mating drive (dopamine, Zhang *et al*., 2016), mate discrimination (octopamine/tyramine, Certel *et al*., 2010; Certel *et al*., 2007), and copulation duration (serotonin, Jois *et al*., 2018; Lee and Hall, 2001). To identify clusters with aminergic-releasing properties, we use marker gene expression of *vesicular monoamine transporter* (*Vmat*), that encodes a transporter for vesicular packaging of biogenic amines (Greer et al., 2005). *Vmat* is a marker gene in three clusters in the full data set, two clusters in the female data set, and is not detected as a marker gene in the male data set (**Figure 6A**, **Supplemental Table 4**), though it is expressed in restricted clusters of neurons in the male data set analysis (**Supplemental Figure 12A**). This restricted expression pattern of *Vmat* is consistent with other scRNA-seq analyses (Allen *et al*., 2020; Croset *et al*., 2018; Davie *et al*., 2018).

**Figure 6.**
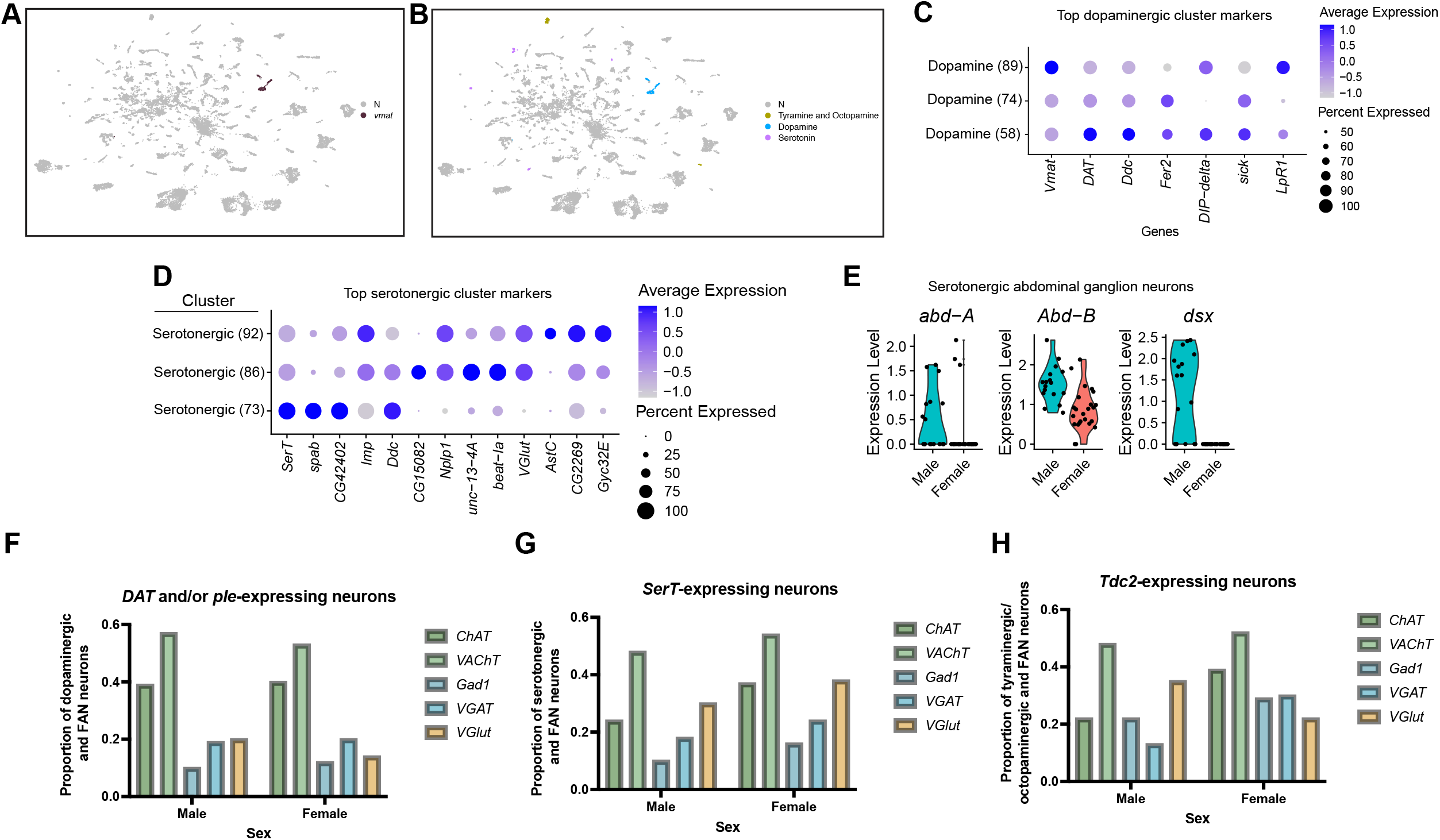
Annotation of neurons that produce aminergic neurotransmitters. (**A**) UMAP plot showing clusters where *Vmat* is a marker gene. (**B**) UMAP plot showing clusters of neurons that produce aminergic neurotransmitters are identified based on marker gene expression: Tyramine and Octopamine (*Tdc2*), Dopamine (*Dat* and/or *ple*), Serotonin (*SerT*), in the full data set (**Supplemental Table 4**). The N (grey) indicates clusters with no aminergic marker genes expression. (**C-D**) Dot plots of top 5 genes for the three dopaminergic clusters (**C**) and three serotonergic clusters (**D**), based on log fold-change in gene expression. Dot size indicates the percentage of neurons expressing each gene per cluster (Percent Expressed). Average normalized expression level is shown by color intensity (Average Expression). (**E**) Violin plots showing expression of *abd-A*, *Abd-B*, and *dsx* in male and female serotonergic cells predicted to be from the abdominal ganglion VNC neurons. These cells were identified by *SerT*, *Abd-A* and/or *abd-B* expression, resulting in 41 neurons (17 male and 24 female). (**F-H**) Proportion of *fru P1* dopaminergic neurons (*DAT* and/or *ple*-expressing) (**F**) serotonergic neurons (SerT-expressing) (**G**), and octopaminergic neurons (*Tdc2*-expressing) (**H**) that co-express genes indicative of FAN-expression in males and females. The proportion that are aminergic and cholinergic (*ChAT* and *VAChT*) are shown with green bars, GABAergic (*Gad1* and *VGAT*) are shown with blue bars, and glutamatergic (*VGlut*) are shown by orange bars. Legend to right of each plot.

Next, we examined the spatial expression patterns of *Vmat* ∩ *fru P1* neurons (**Supplemental Figure 12D-G**). At 48hr APF in both sexes, we find limited expression in the brain and no expression in the VNC (**Supplemental Figure 12D-E**). In 0-24hr adults, we also observe limited expression in the brains of both sexes, with projections around the mushroom body γ lobes (**Supplemental Figure 12F-G**). Further, in 0-24hr adults, there is expression in the VNC of both sexes, consistent with previous reports of *Vmat* expression in the 5 day adult VNC (**Supplemental Figure 12F-G**)(Allen *et al*., 2020). The restricted *Vmat* ∩ *fru P1* expression is consistent with the limited expression in the UMAP clusters.

Below, we additionally annotate the aminergic clusters based on marker gene expression for genes that encode biosynthetic enzymes or transporters of specific aminergic transmitters (**Figure 6B, Supplemental Figure 12H-I, Supplemental Table 4**). Dopaminergic neurons were classified by marker gene expression of *Dopamine transporter* (*DAT*) and/or *pale* (*ple*). Serotonergic neurons were defined by maker gene expression of *Serotonin transporter* (*SerT*). Tyraminergic and octopaminergic were defined by marker gene expression of *Tyrosine decarboxylase 2* (*Tdc2*). Expression of these genes was also visualized at the single neuron level (**Supplemental Figure 12A-C**), revealing that these genes are also expressed beyond where they are considered cluster marker genes.

### Dopaminergic populations of *fru P1* neurons

The three *Vmat* clusters from the full data set analysis have *DAT* and *ple* among their top enriched marker genes. *Dat* and *ple* are expressed >96% of neurons in these clusters, so these three clusters are annotated as dopaminergic neurons (Clusters 58, 74, and 89, **Figure 6B**, **Supplemental Figure 12C, Supplemental Table 4**). All three dopaminergic clusters annotated here have marker genes previously shown to be in adult midbrain and VNC dopaminergic neurons: *PDGP- and VEGF-related growth factor* (*Pvf3*), *kekkon 1* (*kek1*) and *homothorax* (*hth*) (Allen *et al*., 2020; Croset *et al*., 2018). The three clusters do not have Hox marker genes that are expressed in the VNC, however a small population of neurons in clusters 74 and 89 have *Abd-B* expression (**Supplemental Figure 12J**). The top marker genes are largely overlapping between the three clusters, with *Vmat*, *DAT*, and *ple* among the top 5 (**Figure 6C**). Clusters 58 and 74 also share the IgSF gene *DIP-delta* as a top non-unique marker gene (**Figure 6C**, **Supplemental Table 4**). We showed that *DIP-delta* ∩ *fru P1* neurons in the adult brain have restricted spatial expression pattern, with broader expression in the VNC (Brovero *et al*., 2021). This indicates that some *DIP-delta* ∩ *fru P1* neurons are likely dopaminergic and are likely those previously visualized in the brain (Brovero *et al*., 2021). Further, when we examine *Vmat* ∩ *fru P1* expression patterns in the brain of both sexes (**Supplemental Figure 12D-G**), their expression pattern is reminiscent of *DIP-delta* ∩ *fru P1* neurons, with projections around the mushroom body γ lobes (Brovero *et al*., 2021).

Next, we examined the unique marker genes between each of these three clusters and evaluated their GO enrichment (**Supplemental Table 5**). The unique marker genes for clusters 58 and 74 both show protein domain enrichment for “Immunoglobulin-like domain”. This shows that beyond their shared expression of the IgSF encoding gene *DIP-delta*, they also have enrichment for different repertoires of IgSF encoding genes (**Supplemental Table 5**). In contrast, cluster 89 shows no protein domain enrichment but instead shows pathway enrichment for “signaling by WNT”, “Phosphorylation of ARR”, and “Assembly of receptor complex” (see below for WNT; **Supplemental Table 5**). Additionally, we found that all three of the dopaminergic clusters are strongly female-biased in neuron number (>4 fold more female neurons in normalized data), with all having ≤11 male neurons (cluster 58: 11 neurons, cluster 74: 2 neurons, cluster 89: 1 neuron, **Supplemental Table 3**).

It is known that male dopaminergic *fru P1* neurons have a role in mating drive and copulation duration (Jois *et al*., 2018; Zhang *et al*., 2016), so we next evaluated male dopaminergic clusters identified in the male data set analysis (male clusters 56 and 67, **Supplemental Figure 12H, Supplemental Table 4**). A small population of neurons in both clusters express *Abd-B* and *Ubx*, suggesting that they are from the VNC (**Supplemental Figure 12K**), potentially capturing previously identified neurons involved in male copulation duration (Jois *et al*., 2018). Unlike the dopaminergic clusters identified in the full data set analysis, these two male clusters show little overlap in their marker genes with only five shared marker genes (Supplemental Table 4). To compare these clusters, we visualized expression of their top marker genes, revealing that cluster 56 has *DIP-beta* (**Supplemental Figure 12L, Supplemental Table 4**). In addition, we find cluster 67 has *VGlut* as a top marker gene, suggesting that populations of male glutamatergic *fru P1* neurons are also dopaminergic (**Supplemental Figure 12L**). Further, when we perform GO enrichment on the unique marker genes, we find that both are enriched for “Immunoglobulin-like domain superfamily” containing genes. This shows the male clusters are also defined by different enriched subsets of IgSF domain encoding genes, as we saw above in dopaminergic neurons in the full data set analysis (**Supplemental Table 5**).

### Serotonergic populations of *fru P1* neurons

Next, to identify clusters with serotonergic neurons, we examined the clusters that have *SerT* as a marker gene in the full data set. We identify three serotonergic clusters (Clusters 73, 86, and 92). However, all three clusters also contain a substantial number of neurons without *SerT* expression, with all clusters having less than 70% of their neurons expressing *SerT* (cluster 73: 26%, cluster 86: 26%, and cluster 92: 70%) (**Supplemental Figure 12A-C, Supplemental Table 4**). These clusters have additional marker genes that have been previously identified in serotonergic neurons in the adult midbrain and VNC (Allen *et al*., 2020; Croset *et al*., 2018), which provides further evidence that they should be considered serotonergic clusters. All three serotonergic clusters have *IGF-II mRNA-binding protein* (*Imp*), and two have *Jim Lovell* (*lov*), as marker genes (Croset *et al*., 2018). We also find marker genes encoding transcription factors *ventral veins lacking* (*vvl*) and *Lim3* for clusters 86 and 92, as well as *juvenile hormone inducible 21* (*JhI-21*) as a marker gene in all three clusters. These three genes have previously been shown to be marker genes in adult VNC serotonergic neurons (Allen *et al*., 2020). Further, our data suggests that neurons from two serotonergic clusters have autoregulatory control, as one cluster has marker gene expression of 5HT receptors *5-hydroxytryptamine (serotonin) receptor 1A* and *B* (*5-HT1A* and *5-HT1B,* cluster 73) and another has *5-hydroxytryptamine (serotonin) receptor 2B* (*5-HT2B*, cluster 92) (**Supplemental Table 4**), as previously shown in adults (Allen *et al*., 2020).

To examine expression across these serotonergic clusters, we evaluate the top five marker genes for each serotonergic cluster. We find that all three have *SerT*, *Imp*, *Dopa decarboxylase* (*Ddc*), and *CG2269* as marker genes that are expressed in a high percentage of neurons (**Figure 6D, Supplemental Table 4**). The serotonergic clusters 86 and 92 share enriched expression of *Neuropeptide-like precursor 1* (*Nplp1*) and *VGlut*, suggesting that these serotonergic neurons are also glutamatergic and peptidergic (**Figure 6D, Supplemental Table 4**). This analysis also reveals specific and highly expressed markers for these neuron populations: cluster 86 expresses *CG15082* and cluster 92 expresses the neuropeptide encoding gene *Allatostatin C* (*AstC*) (**Figure 6D, Supplemental Table 4**), indicating molecular differences.

Serotonergic *fru P1* neurons in the abdominal ganglion of the VNC are sexually dimorphic, with ∼8-10 male-specific neurons in this region that have a role in sperm transfer (Billeter and Goodwin, 2004; Lee and Hall, 2001; Lee *et al*., 2001). To identify these neurons in our data set we searched for *SerT*-expressing neurons that also expresses *Abd-A* and/or *abd-B*, from the full data set analysis, resulting in 41 neurons (17 male and 24 female). To investigate sex-specific molecular differences, we examined sex-differentially expressed genes in these 41 neurons (**Supplemental Table 5**). *dsx* is a top marker gene in male neurons (**Supplemental Table 5**). 11 of the 17 male serotonergic abdominal ganglion neurons identified here express *dsx*, whereas the 24 female cells exhibit no *dsx* expression (**Figure 6E**). *dsx* expression is required for the formation of male-specific *fru P1* serotonergic neurons in the abdominal ganglion (Billeter *et al*., 2006), some of which are involved in sperm transfer (Tayler et al., 2012), suggesting that we may have identified these male-specific neurons in our data set.

### Tyraminergic and octopaminergic populations of *fru P1* neurons

Tyraminergic and/or octopaminergic *fru P1* neurons have been shown to have a role in mate discrimination. Artificially activating octopaminergic *fru P1* neurons or using RNA-mediated interference against *fru P1* transcripts in octopaminergic neurons increases male-male courtship (Certel *et al*., 2010; Certel *et al*., 2007). The gene encoding *tyrosine decarboxylase 2* (*Tdc2*), an enzyme involved in synthesizing the amino acid tyrosine to tyramine, is expressed in both tyraminergic and octopaminergic neurons (Roeder, 2005). We identify tyraminergic/octopaminergic neurons using the *Tdc2* marker gene. This results in two tyraminergic/octopaminergic clusters in the full data set and one in the female data set (**Supplemental Table 4)**. *Tdc2* is not detected as a marker gene in the male data set but is expressed in a small population of male cells (**Supplemental Figure 12A**). Interestingly, in the full data set, one of the tyraminergic/octopaminergic clusters, cluster 51, is also identified as a mushroom body KC cluster (**Supplemental Table 4**). This suggests that some previously identified *Tdc2*-expressing mushroom body neurons may also include *fru P1*-expressing neurons (Aso *et al*., 2014). However, to our knowledge, no reports of *fru P1* and *Tdc2* expression have been shown in the mushroom body.

We did not detect *Tdc2* ∩ *fru P1* neurons using the genetic intersectional strategy, with a recently published *Tdc2-Gal4* driver (Deng *et al*., 2019). There are previous reports of overlap based on Fru^M^ antibody staining and a different *Tdc2*-*Gal4* driver (Certel *et al*., 2010; Certel *et al*., 2007), suggesting that there may be limitations to our visualization approach. One additional possibility is that there are *Gal4* expression differences between these *Tdc2-Gal4* driver lines. Further, this may be a Fru^M^-expressing population where *fru^FLP^* is not expressed, given there are Fru^M^-expressing neurons that are *fru^FLP^* negative (Yu *et al*., 2010).

### *fru P1* neurons that produce both aminergic and fast-acting neurotransmitters

We find that one male dopaminergic cluster and two serotonergic clusters have *VGlut* marker gene expression (**Figure 6D**, **Supplemental Figure 12G**, and **Supplemental Table 4**). This suggests that there are neurons that may be both aminergic and FAN producing. This prompted us to evaluate at the neuron level co-expression of genes indicative of biogenic amine and FAN production or transport (**Figure 6F-H**). Here, aminergic neurons were classified based on *DAT* and/or *ple* (dopaminergic), *SerT* (serotonergic), or *Tdc2* (tyraminergic/octopaminergic) expression and we asked about their overlapping expression with genes we previously used to identify FAN-producing neurons. For the three classes of aminergic neurons the majority are also cholinergic, based on their co-expression of either *ChAT* or *VAChT* (**Figure 6F-H**, green bars). The male tyraminergic/octopaminergic neurons have a higher proportion of glutamatergic (*VGlut*) co-expressing neurons compared to GABAergic (*VGAT* or *Gad1*), whereas the opposite is seen in females (**Figure 6H**). Overall, we find a range of *fru P1* aminergic neurons show co-expression with genes indicative of producing/transporting FANs (10-54% of aminergic neurons). This suggests that populations of *fru P1* neurons likely release at least two neurotransmitters.

### Identification of *fru P1* neurons that express circadian clock genes

Research of circadian/sleep behaviors over many decades have identified sexual dimorphisms (reviewed in Andretic and Shaw, 2005; Ho and Sehgal, 2005; King and Sehgal, 2020; Shafer and Keene, 2021), with *fru P1* neurons and mating behavior implicated in mediating sex-differences (Fujii and Amrein, 2010; Fujii et al., 2007; Hanafusa et al., 2013; Sakai and Ishida, 2001; Tauber et al., 2003). GO enrichment analyses on marker genes identified two clusters in the full data set, clusters 108 and 109, enriched with genes that have circadian functions (**Figure 1C, Figure 7A**). The marker genes include *period* (*per*), *timeless* (*tim*), *Clock* (*Clk*), *vrille* (*vri*), and *Pdp1-epsilon* (*Pdp1*) (**Supplemental Table 4**). We also annotated other clusters with circadian maker genes (**Supplemental Table 4**), but 108 and 109 are the only clusters with more than one marker gene with known circadian functions in the full data set. The male data set has one UMAP cluster with *per, tim, Clk, vri*, *Pdp1* marker genes (**Supplemental Table 4**), whereas there are no female UMAP clusters with multiple marker genes with known circadian functions. In the full data set, cluster 109 is male-biased in cell number (>2-fold more after normalization), though both clusters 108 and 109 contain cells from males and females (**Supplemental Table 3**). An examination of the expression of *per, tim, Clk, vri*, *Pdp1* in all neurons did not reveal any additional clusters that are likely to be involved in circadian functions that might have been missed when only examining marker genes in clusters (**Supplemental Figure 13A-C**).

**Figure 7.**
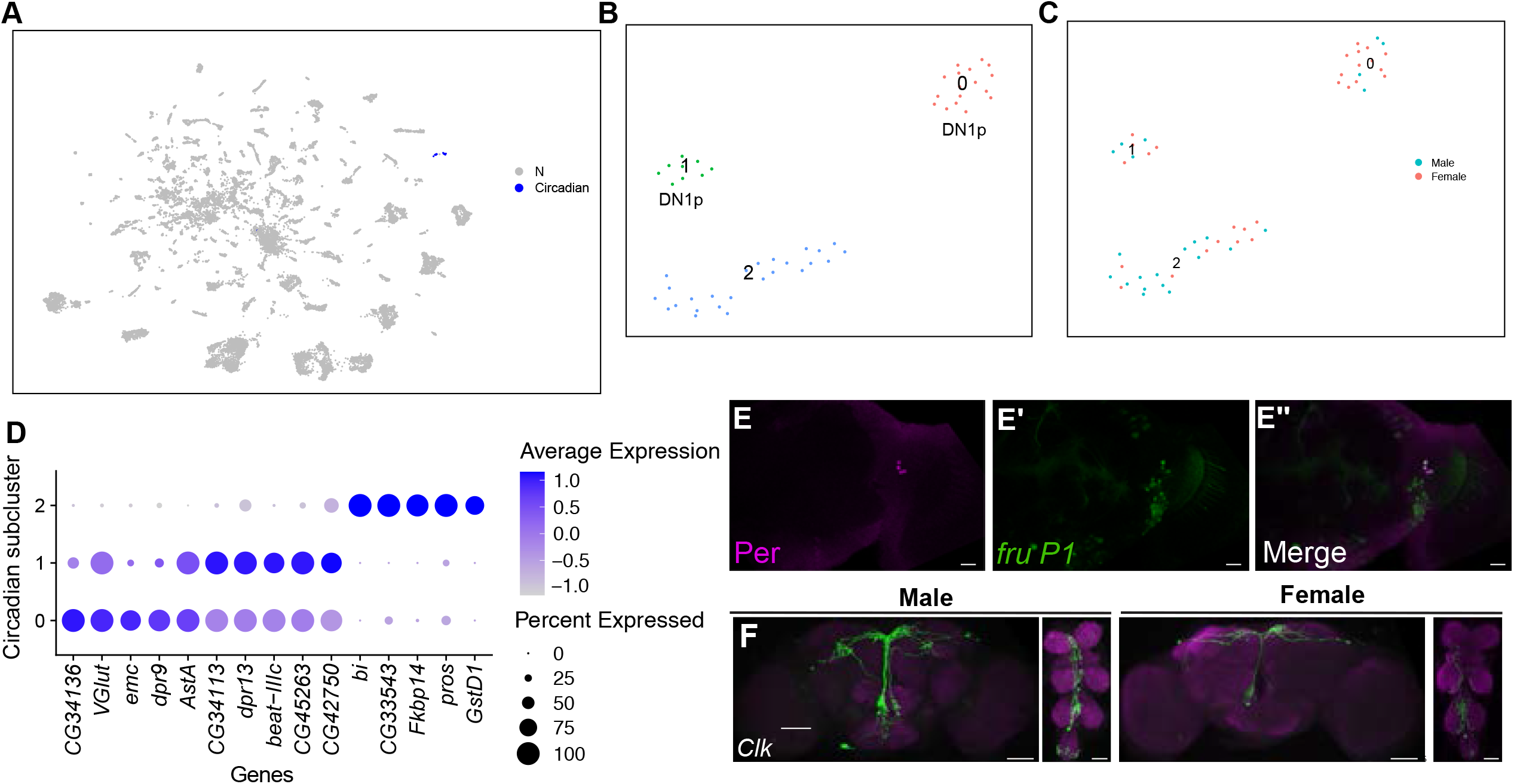
Annotation of neurons that express circadian clock genes. (**A**) UMAP plot showing annotated as circadian clock neurons (blue), in full data set (**Supplemental Table 4**). (**B**) UMAP of subclustering analysis of neurons from annotated circadian clusters (colored clusters from panel **A**), defining 3 subclusters. (**C**) The 3 subclusters are shown with sex indicated by color. The male cells are in blue and female cells are in pink. (**D**) Dot plot of top 5 marker genes in each circadian subcluster based on log fold-change gene expression (**Supplemental Table 4**). Dot size indicates the percentage of neurons expressing each gene per cluster (Percent Expressed). Average normalized expression level is shown by color intensity (Average Expression). (**E-E”**) Confocal maximum projections of brains immunostained for Per (magenta) and *fru P1* >*mcd8::GFP* neurons, from 48hr APF male pupae, at 40x magnification. The right brain hemisphere is shown. **(F)** Brain and VNC confocal maximum projections for *Clk856* ∩ *fru P1* expression in 0-24hr adult males (left) and females (right). Scale bars = 50 μm.

To further examine the neurons in cluster 108 and 109, we performed a subclustering analysis, resulting in three subclusters that each have male and female neurons (**Figure 7B-C**). Expression of *per, tim, Clk, vri*, *Pdp1* is different in the three subclusters (**Supplemental Figure 13D**), as is expression of the top five maker genes, (based on log-fold-change in expression, **Figure 7D**), suggesting the three subclusters are comprised of different circadian regulatory neurons, or the neurons could have distinct gene expression profiles due to slight differences in time of day when the neurons were processed for 10x Genomics library preparation. Based on expression data from a recent scRNA-seq study of clock neurons (Ma *et al*., 2021), we postulate that subcluster 0 is most similar to DN1p neurons given this cluster has *gl*, *VGlut*, *Rh7*, and *Dh31* as marker genes (**Figure 7B**, **Supplemental Table 6**). We postulate that subcluster 1 has similar expression to DN1p neurons, due to having *Rh7* as a marker gene (Kistenpfennig et al., 2017; Ma *et al*., 2021). Consistent with these observations, DN1s have been shown to express Fru^M^ in adults and DN1ps show a sex-difference in cell number in adults (Fujii and Amrein, 2010; Hanafusa *et al*., 2013). Marker gene expression in Subcluster 2 indicates that neurons in this cluster are similar to s-LNvs or l-LNvs clock neurons, because *cry* is a marker gene (Yoshii et al., 2008). Previous studies of developing clock neurons, based on *per* and *tim* expression, indicate that two DN1 neurons, four small ventral LNs (s-LNvs), and two large ventral LNs (l-LNvs) are present at 48hr APF (Kaneko et al., 1997). However, we are not able to visualize DN1s or l-LNv neurons at 48hr APF in either sex with Per antibody staining. We do identify four s-LNvs in both sexes with Per antibody staining at 48hr APF, three of which overlap with *fru P1* expression (**Figure 7E-E’’, Supplemental Figure 13E-E’’**). This supports that subcluster 2 may be comprised of developing s-LNvs. However, there is no expression of *pdf* in subcluster 2, leaving open the possibility that these neurons may be a different subpopulation of circadian neurons that are present in pupae (Helfrichforster, 1995; Kaneko *et al*., 1997).

To visualize circadian neurons that overlap with *fru P1* neurons, we use the genetic intersection approach to visualize *Clk856* ∩ *fru P1* neurons in the CNS. The *Clk856-Gal4* transgene was recently used to classify clock neuron expression in adults, using scRNA-seq (Ma *et al*., 2021). We did not detect *Clk856* ∩ *fru P1* > *sm.GDP.Myc* neurons at 48hr APF in either sex, perhaps due to a lag in reporter gene expression, as noted above. When we examined 0-24hr adults, we find *Clk856* ∩ *fru P1* > *sm.GDP.Myc* neurons that have cell bodies positioned in the DN1 and DN3 circadian regions. Additional cell bodies are detected in suboesophageal zone (SEZ) with projections in the SEZ or in the median bundle, and cell bodies are in several regions of the VNC (**Figure 7F**). When we quantified the number of *Clk856* ∩ *fru P1* > *sm.GDP.Myc* cell bodies in the brain, we observe that males have significantly more DN3 neurons at 0-24hrs (**Supplemental Table 7)**. The male-biased number of *fru P1*-expressing DN3s has not been previously described, to our knowledge (**Figure 7F, Supplemental Table 7**). We note that it is uncertain if DN3 neurons would be present in our 48hr APF scRNA-seq data set, as DN3 neurons are thought to arise later in pupal development (Kaneko *et al*., 1997). In 4-7 day adults, this sex-difference in the number of *Clk856* ∩ *fru P1* DN3 neurons does not persist, but we find a larger number of DN3s in both males and females. Now, we find a significant male-bias in the number of *Clk856* ∩ *fru P1* DN1 neurons at 4-7 days. This is in agreement with previous studies that have quantified *fru*-expressing DN1s and found a larger number in males (**Supplemental Figure 13F-I, Supplemental Table 7**) (Hanafusa *et al*., 2013). In addition, in 4-7 adults, now equal numbers of LNds are detected with *Clk856* ∩ *fru P1* > *sm.GDP.Myc* in both sexes.

To examine the behavioral role of *Clk856* ∩ *fru P1* neurons, we activated these neurons using the intersectional genetic approach, with *UAS<stop<TrpA1^Myc^* (*Clk856* ∩ *fru P1* > *TrpA1^Myc^*) (von Philipsborn et al., 2011). We assayed the effect on locomotor activity patterns using the *Drosophila* activity monitor (DAM, Trikinetics) system, in 12hr light: 12hr dark (LD) conditions. The TrpA1 channel is Myc-tagged, so we further visualized *Clk856* ∩ *fru P1* neurons in 4-7 day adults. Here, we see a significantly larger number of male DN1s, LNds, and both types of SEZ neurons, as compared to females (**Supplemental Figure 13, Supplemental Table 7)**. Generally, when we find significant sex-differences in *Clk856* ∩ *fru P1* > *TrpA1^Myc^* or *Clk856* ∩ *fru P1* > *sm.GDP.Myc* the effect size is small.

To examine locomotor activity, we raised flies at 19°C, a temperature where the TrpA1 channel is inactive, and recorded activity in the DAM system at 25°C, a temperature where the TrpA1 channel is activated (von Philipsborn *et al*., 2011). When we compare activity between the sexes across genotypes, we find that CS males are significantly more active than females (**Supplemental Figure 14A**). This is not observed in the experimental comparisons of *Clk856* ∩ *fru P1* > *TrpA1^Myc^* males and females, where the trend is that females are more active (**Supplemental Figure 14B**). However, when we examine sleep, females have significantly reduced sleep only in the comparisons of *Clk856* ∩ *fru P1* > *TrpA1^Myc^* males and females. A large difference in daytime sleep is observed in *Clk856* ∩ *fru P1* > *TrpA1^Myc^* males and females, where males have a large amount of sleep and females very little (**Supplemental Figure 14C**). There are no significant differences between males and females in nighttime sleep for any genotype (**Supplemental Figure 14D**). *Clk856* ∩ *fru P1* > *TrpA1^Myc^* females have significantly reduced sleep bout length, and significantly increased bout number, compared to males (**Supplemental Figure 14E-F**). We also observe the most significant differences between overall day and night activity in *Clk856* ∩ *fru P1* > *TrpA1^Myc^* females, with no significant differences observed in *Clk856* ∩ *fru P1* > *TrpA1^Myc^* males, consistent with the changes in sleep in *Clk856* ∩ *fru P1* > *TrpA1^Myc^* females (**Supplemental Figure 14G**). To evaluate activity across the 24hr day, we examined actograms for all genotypes (**Supplemental Figure 14 H-I**). *Clk856* ∩ *fru P1* > *TrpA1^Myc^* males exhibit little daytime activity (**Supplemental Figure 14 H**). *Clk856* ∩ *fru P1* > *TrpA1^Myc^* females show robust activity in the early afternoon compared to all other genotypes (**Supplemental Figure 14I**). This suggests that the *Clk856* ∩ *fru P1* neurons have sleep-reducing roles in females, and sleep-promoting roles in males. These neurons may normally direct the dimorphism in daytime sleep, which has been referred to as the male daytime siesta (Andretic and Shaw, 2005; Ho and Sehgal, 2005). Furthermore, it has been shown DN1 neurons have a role in promoting the daytime siesta (Guo et al., 2016), which are activated in our *Clk856* ∩ *fru P1* neurons.

Next, we determined if *Clk856* ∩ *fru P1* neurons had a role in regulating circadian period. To test this, we conducted the DAM assay in 12hr dark:12hr dark (DD) conditions after entraining the flies in LD. Consistent with the LD data, actograms containing two days of LD data show reduced daytime activity in *Clk856* ∩ *fru P1* > *TrpA1^Myc^* males and increased activity in the early afternoon in *Clk856* ∩ *fru P1* > *TrpA1^Myc^* females (**Supplemental Figure 15A-B**). Remarkably, these actograms also show that *Clk856* ∩ *fru P1* > *TrpA1^Myc^* males have an observable activity shift over the 10 DD days, indicative of a shortening circadian period (**Supplemental Figure 15A-B**). To further evaluate the circadian period in DD for all genotypes, we examined periodograms, period peaks, and period strength (**Supplemental Figure 15C-E**). Strikingly, this analysis reveals a shortened period in *Clk856* ∩ *fru P1* > *TrpA1^Myc^* males (mean period=22.8hrs), and a longer period in *Clk856* ∩ *fru P1* > *TrpA1^Myc^* females (mean period=24.6hrs, with a small effect size), compared to control genotypes (**Supplemental Figure 15C-D**). When we examine period strength we find that *Clk856* ∩ *fru P1* > *TrpA1^Myc^* also exhibit the strongest period strength compared to all other genotypes. Altogether, we find that activating *Clk856* ∩ *fru P1* > *TrpA1^Myc^* neurons has sexually dimorphic effects on circadian period, with males having a shortened period and females having a lengthened period. This could be due to quantitative sex-differences in number, sex-differences in the connectivity, or sex-differences in their intrinsic physiology.

### Neuropeptide and neuropeptide receptor expression in *fru P1* neurons

Out of the 49 annotated neuropeptide-encoding genes in the Drosophila genome (Larkin et al., 2021), we find that 47 genes are expressed in *fru P1* neurons in the full data set analysis (**Figure 8A**), and 18 genes are identified as marker genes (**Supplemental Table 4**). In addition, 64% of the clusters in the full data set (72 clusters) have at least one neuropeptide-encoding gene as a marker gene (**Supplemental Figure 4**). Here, we focus on those with known roles in reproductive behaviors in females and males.

**Figure 8.**
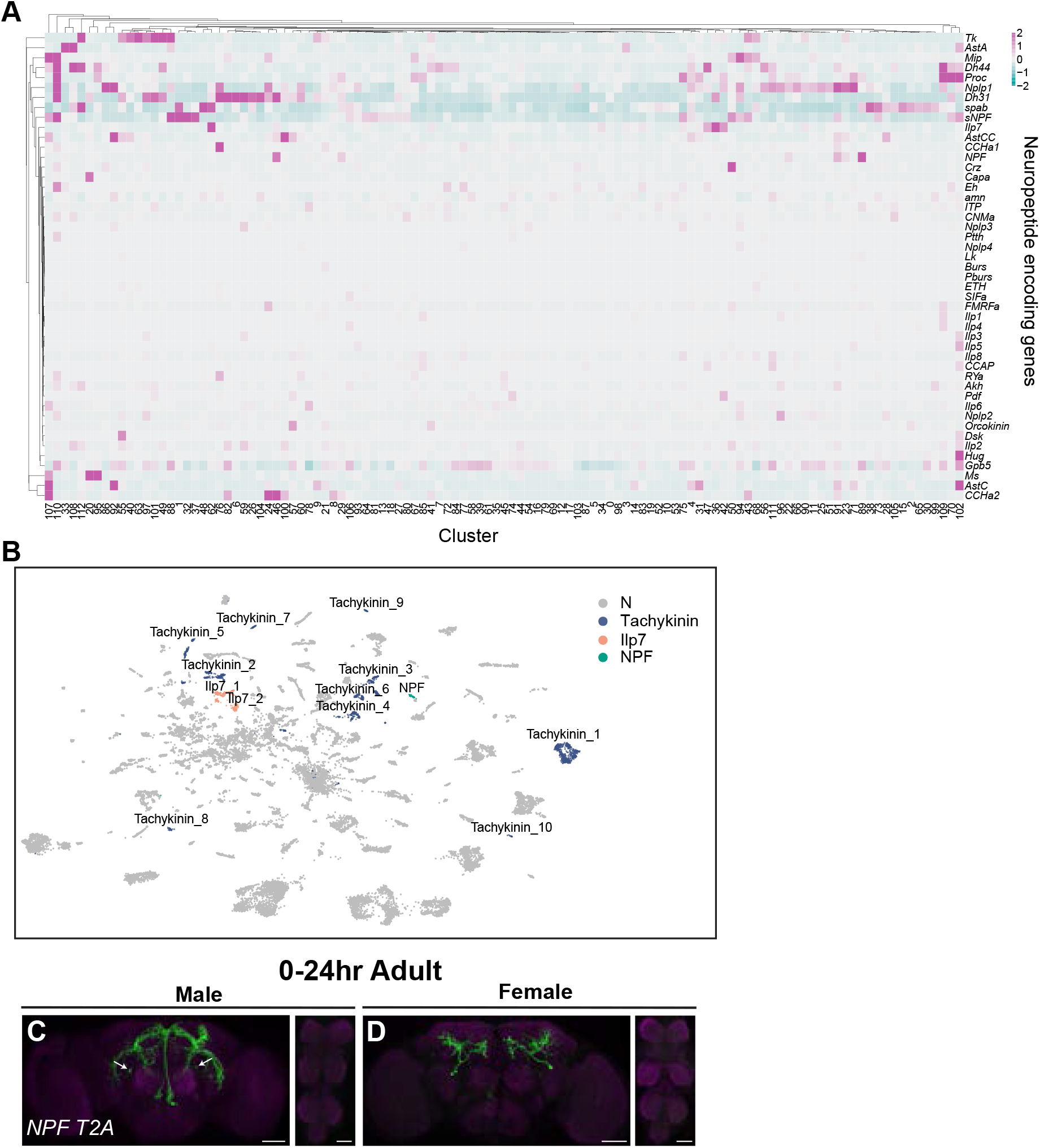
Annotation of neurons that express neuropeptides. (**A**) Heatmap of log-normalized gene expression for neuropeptide encoding genes. Gene expression values were mean-scaled and log-normalized. (**B**) UMAP plot showing annotated clusters with neurons that produce neuropeptides with known roles in *fru P1* neurons for directing reproductive behavior, in the full data set (**Supplemental Table 4**). **(C-D)** Confocal maximum projections of brains and VNC showing *NPF* ∩ *fru P1* expression in 0-24hr adult males and females, using an *NPF-T2A-Gal4* driver (Deng *et al*., 2019). White arrows on male brain indicate male-specific *NPF* ∩ *fru P1* neurons that have been previously identified (Liu *et al*., 2019). Scale bars = 50 μm.

We first characterize clusters that express genes that encode neuropeptides with known roles in female-specific reproductive behaviors. For example, *Insulin-like peptide 7-* (*Ilp7*) expressing neurons are involved in oviduct function, and *Diuretic hormone 44* (*Dh44*)- expressing neurons have a role in sperm ejection (Castellanos et al., 2013; Garner et al., 2018; Lee et al., 2015). We find *Ilp7* is a marker gene in two clusters, in all three data sets (**Figure 8B**, **Supplemental Figure 16A-B**). *Ilp7* has been shown to be expressed in *fru P1* neurons in both sexes, with female-specific neurons identified in the abdominal ganglion of the VNC, due to cell death in males (Garner *et al*., 2018). However, neither cluster with *Ilp7* as a marker gene in the full data set is female-specific nor show a sex-bias in cell number (**Supplemental Table 3**), so it is possible that we did not identify the female-specific *Ilp7* neurons. *Dh44* has been shown to have a role in female sperm storage post-mating, with six cholinergic *Dh44*-expressing neurons in the Pars Intercerebralis of the female brain shown to have a role in sperm ejection behavior (Lee *et al*., 2015). We find *Dh44* is a marker gene in two clusters in both the male and female data sets and six clusters in the full data set (**Supplemental Table 4**). In the full data set, three of these clusters with *Dh44* as a marker gene also have marker genes that indicated they are cholinergic (*ChAT* and/or *VAChT* are marker genes, **Supplemental Table 4**). This previous work showed co-expression of *fru P1* with the gene that encodes the *Dh44* receptor (*Dh44-R1*) but did not examine if *Dh44-*expressing neurons were *fru P1*-expressing (Lee *et al*., 2015). Using a *T2A*-*Gal4* knock-in in the *Dh44* gene and the genetic intersectional approach (Deng *et al*., 2019), we visualized the spatial expression of *Dh44* ∩ *fru P1* neurons, finding expression at 48hr APF and in 0-24hr adults in both sexes (**Supplemental Figure 16C-F**). In 0-24hr adults, this *Dh44* ∩ *fru P1* expression pattern includes neurons in the Pars Intercerebralis, which may contain the neurons implicated to have a role in female sperm ejection.

*fru P1* neurons that produce neuropeptides also have known roles in male courtship, copulation, and aggression behaviors (Asahina et al., 2014; Liu et al., 2019; Tayler *et al*., 2012; Wohl et al., 2020; Wu et al., 2019). For example, a subset of male-specific *Tackykinin* (*Tk*) and *fru P1* co-expressing neurons in the brain promote male aggression, though a larger number of *Tk*-expressing neurons have been identified in both sexes (Asahina *et al*., 2014; Nassel et al., 2019; Wohl *et al*., 2020). Here, we find that *Tk* is a marker gene in 10 clusters in the full data set, and five in both the male and female data set analyses (**Figure 8B, Supplemental Figure 16A-B, Supplemental Table 4**). This suggests that beyond the male-specific *Tk-Gal4* and *fru P1* co-expressing neurons that have been well characterized for their role in aggression (Asahina *et al*., 2014; Wohl *et al*., 2020), there are additional *Tk*-expressing neurons that co-express *fru P1* in both sexes.

Male-specific *Neuropeptide F* (*NPF*) and *fru P1* co-expressing neurons have a role in regulating courtship drive, by inhibiting courtship in sexually satiated males (Liu *et al*., 2019). We annotate two clusters where *NPF* is a marker gene in both male and full data set analyses (**Figure 8B, Supplemental Figure 16A, Supplemental Table 4**). *NPF* is not a marker gene in the female data set (**Supplemental Table 4**). To spatially visualize *NPF* ∩ *fru P1* neurons, we used the genetic intersection approach with an *NPF T2A*-*Gal4* knock-in (Deng *et al*., 2019). There were no visible *NPF* ∩ *fru P1* neurons at 48hr APF in either sex. In 0-24hr adult brains, we identify dramatically different sexually dimorphic neuronal projections, with some cell bodies in similar positions (**Figure 8C-D**). Based on location of the neurons, we potentially find the previously described male-specific *NPF* neurons in the brain using the intersectional approach (**Figure 8C-D**, white arrows). However, the *NPF* ∩ *fru P1* expression pattern differs from previous reports that used different *NPF*-*Gal4* or *LexA* insertions (Liu *et al*., 2019). Additionally, when we examined expression of all genes that encode neuropeptides, we find additional clusters that express the neuropeptides *Corazonin* (*Crz*) and *Drosulfakinin* (*Dsk*), which have been shown to be expressed in subpopulations of *fru P1* neurons and have impacts on male behaviors, but are not considered marker genes in our data sets (**Figure 8A**) (sperm transfer, Tayler *et al*., 2012; courtship inhibition, Wu *et al*., 2019).

Neuropeptide receptors are also widely expressed in *fru P1* neurons, with 104 clusters having at least one gene encoding a neuropeptide receptor as a marker gene in the full data set (**Supplemental Table 4**). Previous studies have found that *fru P1* neurons that express neuropeptide receptors have roles in male courtship behaviors. These include *Corazonin receptor* (*CrzR*) (sperm transfer, Tayler *et al*., 2012), *Cholecystokinin-like receptor at 17D3* (*CCKLR-17D3*) (courtship inhibition, Wu *et al*., 2019), and *SIFamide receptor* (*SIFaR*) (increased male-male courtship, Sellami and Veenstra, 2015). In addition, the neuropeptide receptor *Leucine-rich repeat-containing G protein-coupled receptor 3* (*Lgr3*) has been shown to expressed in *fru P1* neurons and have a role in female behavior (female receptivity and fecundity, Meissner et al., 2016). All these aforementioned neuropeptide receptors are found as marker genes in our data set (**Supplemental Table 4**). We visualized *CrzR* ∩ *fru P1* and *Lgr3* ∩ *fru P1* neurons using *T2A*-*Gal4* knock-in driver lines (Deng *et al*., 2019) (**Supplemental Figure 16G-N**). *CrzR* ∩ *fru P1* neurons in the abdominal ganglia have a known role in male copulation behavior (Tayler *et al*., 2012). We find that these neurons are in similar positions as previously shown in males and additionally visualize sexually dimorphic neurons in the brain as well as a small set of neurons in the female abdominal ganglia (Tayler *et al*., 2012) (**Supplemental Figure 16G-J**). The 48hr APF *Lgr3* ∩ *fru P1* neurons show expected median bundle expression, with more GFP-expression in females compared to males, consistent with *Lgr3* expression being regulated by Fru^M^ (**Supplemental Figure 16K-L**) (Meissner *et al*., 2016). However, in 0-24hr adult males, we observe male-specific *Lgr3* ∩ *fru P1* neuronal projections in the lateral protocerebral complex and in the VNC that were not previously reported (**Supplemental Figure 16M-N**), which may be due to differences in visualization strategies. Given that 109 of the 113 clusters in the full data set express at least one neuropeptide or neuropeptide receptor as a marker gene, this indicates that nearly all *fru P1*-expressing neurons use neuropeptides for signaling (**Supplemental Table 4**).

### *nAChR* receptor subunit genes expressed in *fru P1* neurons

We find nicotinic acetylcholine receptor (nAChR) subunits are marker genes for the majority of clusters in the full data set (91 clusters out of 113) (**Supplemental Table 4**). Nicotinic acetylcholine receptor are pentameric receptors that assemble with combinations of 10 subunits [seven α nAChR subunits (1-7) and three β nAChR subunits (1-3)] (Rosenthal and Yuan, 2021). Of these 10 *nAChR* subunit genes, we find that nine are marker genes and all 10 are expressed in the full data set (**Supplemental Table 4).** Furthermore, they are broadly expressed with at least 97% of *fru P1* neurons expressing at least one *nAChR* receptor. Out of the 10 subunits, *nAChRα5* is the most widely expressed in all three data sets and *nAChR*β*3* has the narrowest expression (**Supplemental Table 8**). The different subunit combinations confer different physiological binding properties (Perry et al., 2021). To evaluate subunit co-expression, we examined the pairwise expression correlation between the 10 subunits for the three data set analyses and show significant correlations (**Figure 9A-C**). For all data sets, we find that the overall correlation trends are similar, with some additional significant correlations observed for the female data set (**Figure 9Q-S**). We find the highest positive expression correlations are between: *nAChRα5* and *nAChR*β*1*, *nAChRα6* and *nAChRα7*, and *nAChRα1* and *nAChRα2* for the full data set analysis (**Figure 9C**). In contrast to the analysis of subunit correlations performed in the adult mid-brain data set, we find anti-correlations between subunit expression data (**Figure 9A-C**) (Croset *et al*., 2018). These negative correlations occur between *nAChRα5* and *nAChRα6* and *nAChRα5* and *nAChRα7*, with the strongest negative correlations in the female data set analysis (**Figure 9A-C**). Given that *nAChRα5* and *nAChRα6* has the strongest expression correlation in the adult midbrain data set and we observe this pair as the strongest expression anti-correlation points to the importance of examining different neuronal populations using single cell approaches.

**Figure 9.**
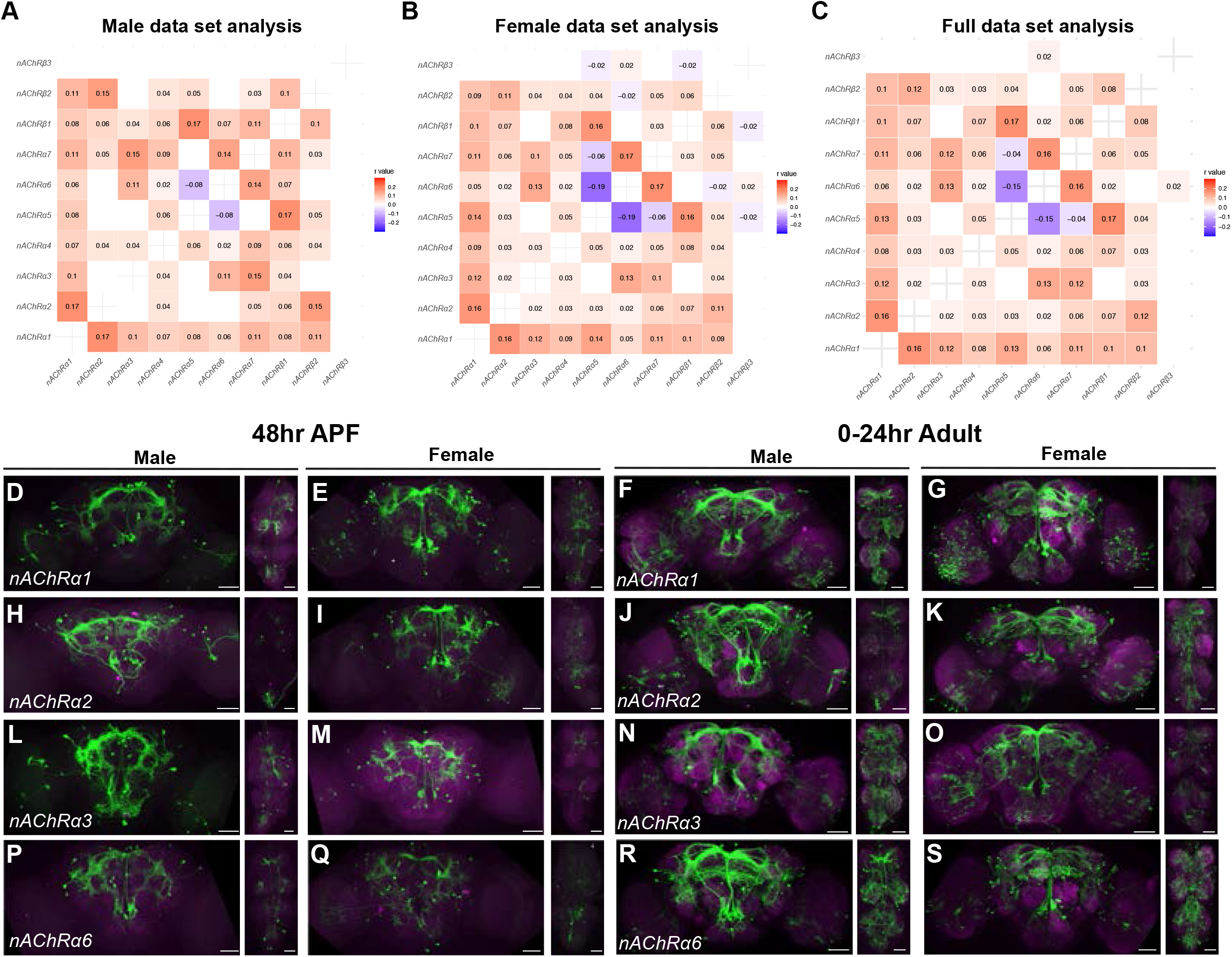
*nAChR* subunit gene expression in *fru P1* neurons. (**A-C**) Heatmap showing correlation of *nAChR* subunit gene expression in neurons in the male data set (**A**), female data set (**B**), and full data set (**C**). Pearson’s r values denoted on heatmap with color, according to legends. (**D-S**) Confocal maximum projections of brain and VNC for *nAChRα* ∩ *fru P1*-expressing neurons, with intersecting expression shown in green for males and females, as indicated. Male and female animals from 48hr APF and 0-24 hour adults are shown, as indicated. Scale bars = 50 μm.

We next visualized the spatial expression patterns for some *nAChRα* that have available Gal4 lines. We examined nAChR*α1*, *nAChRα2*, *nAChRα3*, and *nAChRα6* ∩ *fru P1* neurons and find broad expression at 48hr APF and in 0-24hr adults of both sexes, which is consistent with our scRNA-seq expression data (**Figure 9D-S**). All *nAChR* ∩ *fru P1* spatial populations examined have similar expression patterns within each sex, which is consistent with their correlated co-expression (**Figure 9D-S**). The *nAChR* ∩ *fru P1* spatial populations are in all segments of the VNC in both sexes (**Figure 9D-S**). We also find that spatial expression of *nAChRα1*, *nAChRα2*, and *nAChRα6* ∩ *fru P1* neurons in 0-24hr adult male brains contain previously characterized sexually dimorphic projections in the tritocerebral loop (**Figure 9D-S**). Notably, we find that *nAChRα3* ∩ *fru P1* expression is absent from the mushroom body in both sexes, whereas we observe mushroom body expression for all other *nAChR*s ∩ *fru P1* examined (**Figure 9D-S**). We also find that *nAChRα3* is not a marker gene in mushroom body clusters identified in our data sets (**Supplemental Table 4**), consistent with previous reports indicating its absence from the mushroom body (Croset *et al*., 2018; Shih et al., 2019). Our expression findings here, and the identification of several *nAChR*s in genomic studies examining regulation by Fru^M^, expression in *fru P1* neurons, and expression changes post-mating, suggest that *nAChR*s are important for *Drosophila* reproductive behaviors (Brovkina *et al*., 2021; Dalton *et al*., 2013; Neville *et al*., 2014; Newell *et al*., 2016; Newell et al., 2020; Palmateer *et al*., 2021; Vernes, 2014).

### *Wnt* expression in *fru P1* neurons

Wnts have been shown to have a range of function in neurons, including axon guidance, synapse formation, neuronal plasticity, morphogenesis, and signaling in several species (He et al., 2018; Kolodkin and Tessier-Lavigne, 2011; Salinas and Zou, 2008; Sanes and Yamagata, 2009). Here, we further examine *Wnt* expression, given the GO enrichment analysis of marker genes in the full data set identified several wingless pathway terms including: ‘WG ligand binding’, ‘wingless pathway’, ‘Beta-catenin independent WNT signaling’, and ‘signaling by WNT’ (**Supplemental Table 5**). The analyses of dopaminergic clusters also identified cluster 89 as having marker genes enriched with the GO term ‘signaling by WNT’ (see above; **Supplemental Table 5**). Furthermore, our previous genome-wide-association study showed four WNT pathway signaling genes had variants associated with differences in female remating behavior (Newell *et al*., 2020), adding further impetus to evaluate the Wnt pathway. We found three *Wnt* genes were marker genes, *Wnt4* (14 clusters), *Wnt5* (2 clusters) and *Wnt10* (4 clusters) (**Supplemental Table 4**).

We visualized the spatial expression patterns for *Wnt4* ∩ *fru P1* and *Wnt10* ∩ *fru P1* neurons and observe several sexual dimorphisms in their neuronal projection patterns at 48hr APF and in 0-24hr adults of both sexes (**Supplemental Figure 17A-H**). We additionally visualized *Wnt5* ∩ *fru P1* in 0-24hr adults (**Supplemental Figure 17I-J**). In 0-24hr adult males, *Wnt4* ∩ *fru P1* neuron projections look similar to previously identified sexually dimorphic vPR1/vPr-a ascending neurons (**Supplemental Figure 17C**, white arrows), which have been suggested to have a role in sensory integration and motor output to control wing song (Cachero *et al*., 2010; Yu *et al*., 2010). *Wnt10* ∩ *fru P1* neurons are observed only in the brain, and not the VNC, and show sexually dimorphic projections (**Supplemental Figure 17E-H**, white arrows). Male *Wnt10* ∩ *fru P1* neurons project across hemispheres at both 48hr APF and 0-24hr stages (**Supplemental Figure 17E and G**, white arrows), whereas in female brains no projections between the brain hemispheres were observed (**Supplemental Figure 17F and H**, white arrows). Sexually dimorphic midline crossing of gustatory sensory neurons was previously observed in the VNC and regulated by *fru* and *dsx* (Mellert et al., 2010; Possidente and Murphey, 1989). The observation that *Wnt10* ∩ *fru P1* neurons also have sexually dimorphic midline crossing in the brain suggest this may be a mechanism to generate sex differences in behavior. In 0-24hr adults, *Wnt5*∩ *fru P1* shows broader expression in the brain than *Wnt4* ∩ *fru P1* and *Wnt10* ∩ *fru P1* neurons, and also exhibits expression in all VNC segments (**Supplemental Figure 17I-J**).

### Sex differences in gene expression between male and female *fru P1* neurons

We next examined differential expression between the sexes within individual UMAP clusters in the full data set analysis. This analysis reveals 6,162 genes are significantly differentially expressed between males and females, at the cluster level (3,504 higher in males, 2,658 higher in females). When we reduce these gene lists to remove redundant genes (the same genes are differentially expressed in more than one cluster) and compare between males and females, we find 711 genes uniquely male-biased in expression (always show higher expression in males within at least one cluster), 586 are uniquely female-biased in expression (always show higher expression in females within at least one cluster) (**Supplemental Table 9**). There are 147 genes that are either male- or female-biased, depending on the cluster (**Supplemental Table 9**). When we compare these sex-biased genes to genes identified as sex-biased in our previous cell-type-specific TRAP study at 48hrAPF, we find significant overlap between gene lists (**Supplemental Table 4**) (Palmateer *et al*., 2021).

We next performed GO enrichment on the list of genes that have sex-biased expression (**Supplemental Table 5**). The 711 genes with uniquely male-biased expression are mainly enriched for ‘biological process’ terms related to ‘translation’ (**Supplemental Table 5**). Though, when we examine ‘pathway enrichment’ for these genes, we find ‘wingless pathway’ and several circadian clock related terms, which include ‘degradation of CRY’, ‘degradation of PER’, ‘degradation of TIM’, and ‘circadian clock pathway’ (**Supplemental Table 5**). When we examine the enrichment terms for the 586 genes with uniquely female-biased expression we find enrichment for terms related to ‘signaling’ and ‘neuron development’, such as ‘neurogenesis’, ‘axonogenisis’, and ‘synapse organization’ (**Supplemental Table 5**). Further, these uniquely female-biased genes are highly enriched for ‘immunoglobulin-like domain’ encoding genes (**Supplemental Table 5**). The genes that show sex-differential expression in either males or females, depending on the clusters, are also highly enriched for neuronal terms and ‘immunoglobulin-like domain’ encoding genes (**Supplemental Table 5**). This analysis shows that the same genes are re-used in different neuronal clusters to impart sexual dimorphism, given the redundancy in the gene lists found across the clusters.

### Immunoglobulin-like domain superfamily (IgSF) expression in *fru P1* neurons

A Flymine analysis of the marker gene lists to find ‘protein domain’ enrichments identifies ‘Immunoglobulin-like domain superfamily’ with the most significant enrichment in each of the three data sets (**Supplemental Table 5**) (Lyne *et al*., 2007). In the full data set analysis, we find that all clusters have at least one marker gene that is a member of the Immunoglobulin-like domain superfamily, with most expressing a large and unique repertoire (**Supplemental Table 4**). Using a heatmap we visualized expression of IgSF-encoding genes, which shows the average expression of a gene in a cluster, regardless of marker gene status (**Supplemental Figure 18**). Consistent with what we observe when we examine marker genes (**Supplemental Table 4**), each cluster exhibits unique expression combinations of IgSF encoding genes (**Supplemental Figure 18**).

Previously we examined expression of *dpr* and *DIP* genes in the male CNS from 48 hr APF using 10X single-cell genomic approaches, with *fru P1* neurons identified based on expression of *fru P1* and/or a GFP reporter (Brovero *et al*., 2021). Similar to our previous study on males, we find that in both sexes the majority of *dpr*s show broad expression, whereas *DIP*s exhibit more restricted expression (**Supplemental Figure 19**). With the addition of female data, we next determine if there are *dpr* and *DIP* expression differences between the sexes, for each cluster of the full data set analysis (**Supplemental Figure 19**). This analysis revealed that there are sex-differences in *dpr* and *DIP* expression for several clusters in our data set (**Supplemental Figure 19, Supplemental Table 9**). Of the *dpr*s and *DIP*s that show a significant sex-bias in expression within clusters, some show expression that is always higher in female neurons (uniquely female-biased; *dpr*s *1*, *5*, *8*, *10*, *11*, *12*, *14*, *15*, *16*, *17*, and *20, DIP-alpha* and *DIP-beta*), whereas fewer show expression that is always higher in males (uniquely male-biased; *dprs 7*, *18*, and *21*, *DIP*s *gamma*, *theta*, and *zeta*, **Supplemental Figure 19, Supplemental Table 9**). In contrast, *dpr*s *2*, *3*, *6*, *9*, and *13* and *DIP-delta* and DIP*-eta* show cluster dependent differences in their significant sex-biased expression (**Supplemental Figure 19, Supplemental Table 9**). Differences in *dpr*/*DIP* co-expression within individual *fru P1* neurons may offer a mechanism to generate different cell adhesion properties to mediate synaptic connections, as we previously proposed (Brovero *et al*., 2021).

### Transcription factor expression in *fru P1* neurons

During the stage of metamorphosis that we examined, cell fate decisions are being directed by transcription factor (TF) expression, with different combinations of TFs found in different neuronal populations (Doe, 2017; Shirasaki and Pfaff, 2002). Consistent with this, the top ‘molecular function’ GO enrichment terms for marker genes across all three data set analyses is ‘DNA-binding transcription factor activity’ (**Supplemental Figure 6, Supplemental Table 5**). Out of the 628 *Drosophila* genes encoding TFs (Flybase), 240 are marker genes in the male data set analysis, 228 in the female data set analysis, and 253 in the full data set analysis (**Supplemental Table 4**). We next determined the enriched TF subcategories in Flymine, evaluating subcategories that contained more than 25 genes (Lyne *et al*., 2007). This identified three subcategories of TF marker genes: homeobox-like domain superfamily (74 genes), zinc finger C2H2 (74 genes), and helix-loop-helix (HLH) DNA binding domain superfamily (25 genes). We next examined if certain subcategories have restricted or broad expression, to gain insight into which TFs might impart cluster-specific identities.

Most of the homeobox-like domain superfamily TF encoding genes, including *abd-A*, *Abd-B*, and *slouch* (*slou*), show cluster specificity, with high expression in a large percent of neurons in the cluster, in only a subset of clusters (**Supplemental Figure 20A**).There are some homeobox TF encoding genes, such as *pipsqueak* (*psq*), *extradenticle* (*exd*), and *CG16779*, that are expressed more broadly, in nearly all clusters (**Supplemental Figure 20A**), On the other hand, zinc finger C2H2 TF encoding genes, which includes *fru*, has most genes displaying broad expression (**Supplemental Figure 21**). Though, some zinc finger C2H2 TFs exhibit cluster-specific expression, such as *buttonhead* (*btd*), which is predominately expressed in cluster 101 (**Supplemental Figure 21**). Genes that encode HLH DNA binding domain superfamily TFs also have several showing broad expression across clusters (**Supplemental Figure 22**), and others with more restricted expression. These analyses suggests that the homeobox-like domain superfamily may have a large role in directing differences in cell-types among the *fru P1* neurons, given the overall number of genes with large differences in expression across the clusters (**Supplemental Figures 20-22**).

We next determined if the 253 marker genes that encode transcription factors display significant sex-differential expression. There are 69 genes that encode TFs with cluster-specific sex-differential expression. Seventeen genes are uniquely male-biased, 38 are uniquely female-biased, and 8 are either male- or female-biased depending on the cluster (**Figure 10A** and **Supplemental Table 9**). We used dots plots to visualize the expression of these TFs with cluster-specific differential expression between the sexes (**Figure 10B**). This shows that 49 clusters exhibit significant sex-differential expression of at least one marker gene TF, with several clusters showing differential expression of multiple TFs (**Figure 10B,** black circles). This differential TF expression between the sexes is likely a mechanism to direct sex differences in *fru P1* neurons.

**Figure 10.**
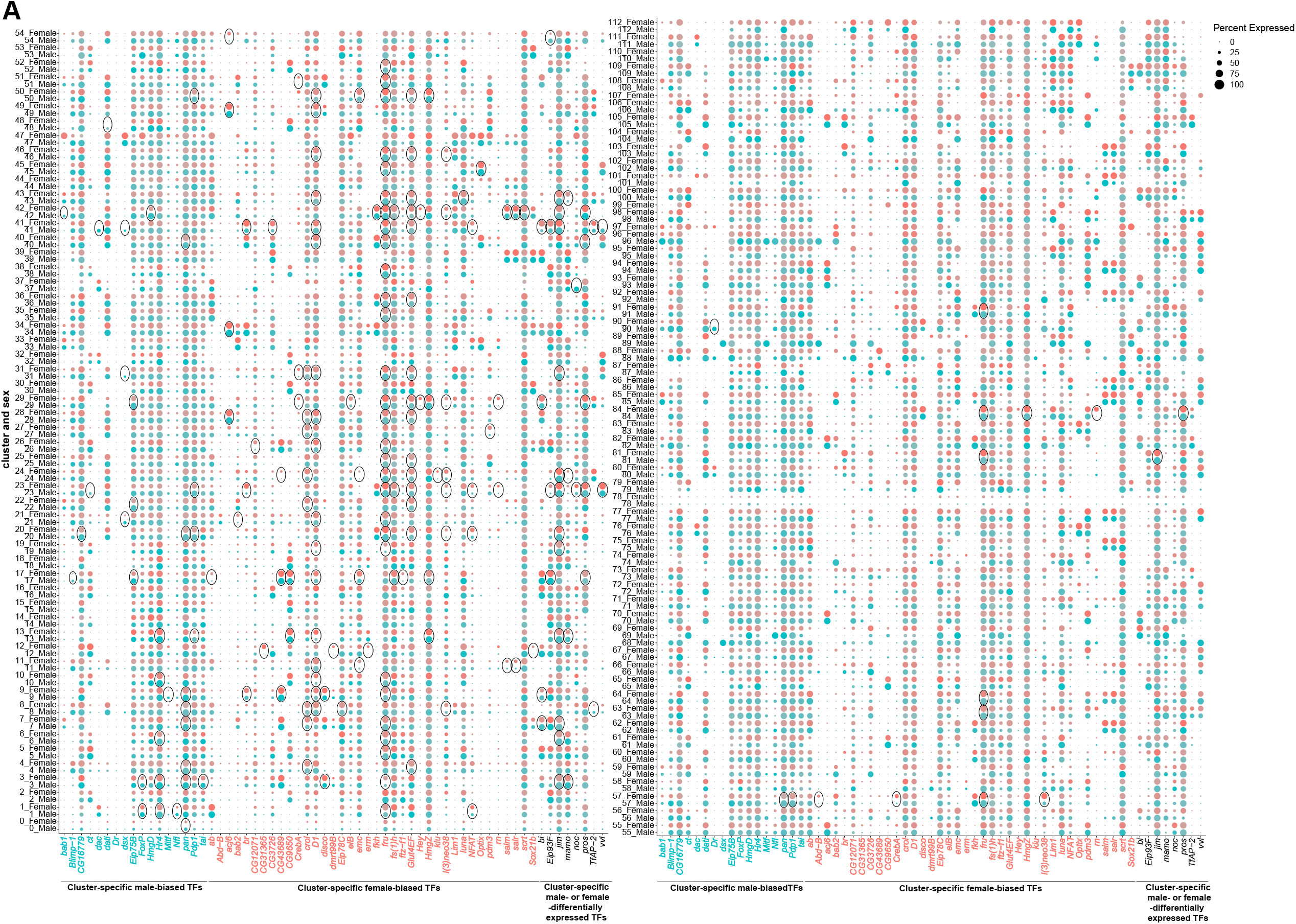
Sex differences in transcription factor (TF) expression in *fru P1* neurons. (**A**) Dot plot split by sex for TF genes that have cluster-specific sex-biased marker gene expression. On the X-axis, marker genes in blue only have male-biased, cluster-specific expression (17 genes). On the X-axis, marker genes in red only have female-biased, cluster-specific expression (38 genes). On the X-axis, marker genes in black have either male-biased expression in some cluster(s) and female-biased expression in other cluster(s) (8 genes). Dot size indicates the percentage of neurons expressing each gene per cluster (Percent Expressed). Male expression is shown in blue and female expression is shown in pink. Black circles indicate clusters where TFs are sex-differentially expressed (**Supplemental Table 9**).

## Conclusions

We performed scRNA-seq on pupal *fru P1* neurons from males and females to understand how sex differences are built into the nervous system to direct adult reproductive behaviors. The full data set analysis revealed the heterogeneity of *fru P1* neurons at 48hr APF with 113 different molecular clusters identified, as well as additional subtypes derived by subclustering analyses. This exceeds the number of previously designated spatial cluster and morphological classifications. This suggests that there are more functional differences in neurons than predicted by Fru^M^/*fru P1* spatial patterns (Billeter and Goodwin, 2004; Cachero *et al*., 2010; Lee *et al*., 2000; Manoli *et al*., 2005; Stockinger *et al*., 2005; Yu *et al*., 2010). The analyses also revealed that nearly all clusters have neurons from both males and females. This suggests that male and female neurons that direct vastly different reproductive behaviors, have a core gene expression program that are overlaid with sex differences in gene expression. This is consistent with our previous TRAP-seq data sets, where we postulated that shared molecular properties underlie common behavioral needs. Sex differences in gene expression are overlaid on a sex-shared baseline gene expression program to direct male and female behavioral differences, with consistently more male-biased genes identified in *fru P1* neurons across all of our studies (Newell *et al*., 2016; Palmateer *et al*., 2021).

## Materials and Methods

### Fly strains and husbandry

Flies for scRNA-seq experiments were the genotype: *w[*]; P[y[+t7.7] w[+mC]=10XUAS-IVS-mCD8::GF]attP40/UAS-Gal4;fru P1-Gal4/+.* This strain expressing membrane bound GFP in fru P1 neurons in males and females. The flies were sex-sorted at the white pre-pupal stage using the male gonad to distinguish between the sexes. We note that the *UAS-Gal4* present in this genotype was determined to be non-functional (Feng, 2017). The pupae were aged to 48 hours after puparium formation (APF) on 3% agar plates with Bromophenol blue for color contrast. All strains, unless otherwise indicated, are aged in a humidified incubator at 25°C on a 12 hour light and 12 hour dark cycle. Additional fly strains used in this study are listed in the Resources Table. The laboratory Drosophila media composition is: 33L H_2_O, 237g agar, 825g dried deactivated yeast, 1560g cornmeal, 3300g dextrose, 52.5g Tegosept in 270ml 95% ethanol and 60ml propionic acid.

### Dissociation of pupal CNS for single cell mRNA sequencing

The CNS (brain and ventral nerve cord) from twenty freshly dissected male or female pupae were used per replicate (n=2 replicates per sex). The dissected tissue was dissociated as previously described (Brovero *et al*., 2021). All dissections occurred within an hour after the lights turned on (ZT0). GFP positive cells (*fru P1*-expressing neurons) were enriched using fluorescent activated cell sorting (FACS) with a FACSAria SORP with BD FACSDiva software.

### 10xGenomics library preparation and sequencing

scRNA-seq libraries were generated using Single Cell 3’ Library & Gel Bead Kit v3, Chip B Kit, and the GemCode 10X Chromium instrument (10X Genomics, CA), according to the manufacturer’s protocol (Zheng et al., 2017). In brief, FACS enriched *fru P1*-expressing neurons were suspended in S2 medium with 10% FBS and the maximum volume of cell suspension (46.6μl) was mixed with single cell Master Mix and loaded into an individual chip channel for each sex and replicate. For all samples, this process occurred between four and five hours after lights on (ZT4-ZT5). The library preparation steps were conducted as previously described, with modifications to polymerase chain reaction (PCR) thermal cycling steps for v3 chemistry (Brovero *et al*., 2021). For cDNA amplification PCR settings were: 98°C for 3 min, 12 cycles of (98°C for 15s, 63°C for 20s), 72°C for 1 min, held at 4°C. For sample index PCR amplification the settings were: 98°C for 45s, 16 cycles of (98°C for 20s, 54°C for 30s, 72°C for 20s), 72°C for 1 min, and hold at 4°C. All single cell libraries were sequenced on an Illumina NovaSeq 6000 at Florida State University Translational Core Laboratory.

### scRNA-seq data processing and clustering in Seurat

CellRanger software (v.3.0.2) “cellranger count” command was used to align sequencing reads to a customized *Drosophila melanogaster* (BDGP6.92) STAR reference genome, which contained the sequence for *mCD8-GFP* cDNA to generate a multidimensional feature-barcode matrix for each replicate. Using the inflection point of the barcode rank plots, the CellRanger pipeline called the number of cells captured in each replicate (**Supplemental Table 1**). We captured 26,149 total cells across both sexes and replicates with 871,092,958 sequencing reads, reaching an average of 65% sequencing saturation (**Supplemental Table 1**). The feature-barcode matrices were next analyzed in Seurat (v3.2.2) (Stuart *et al*., 2019). Each expression matrix was filtered to obtain high quality cells using the following criteria: cells with >5% mitochondrial transcripts (dead/dying cells), <200 genes (empty droplets), those expressing more than 4,000 features (genes) and/or possessing more than 20,000 UMIs (potential doublets or triplets) were removed in each replicate. The male replicates and the female replicates were merged to form sex-specific datasets and additionally all replicates for both sexes were merged into a full data set. This yielded 7,988 male cells, 17,530 female cells and 25,510 cells in total. We used default normalization and scaling steps for each data set as outlined in the Seurat “Guided clustering tutorial” (https://satijalab.org/seurat/v3.2/pbmc3k_tutorial.html). The default “NormalizeData” function was used to normalize the data and the data was subsequently scaled using the default “ScaleData” function. We performed a Principal Component Analysis (PCA) using the top 2,000 highly variable features in each data set. We calculated statistically significant principal components (PCs) for each data set based on JackStraw analysis. We used the number of significant PCs, up to where the first non-significant PC was met, for each data set (**Supplemental Figure 2**). This resulted in 80, 104, and 107 principal components. We proceeded to use Seurat’s standard workflow to reduce dimensionality and cluster cells by default “FindNeighbors”, “FindClusters”, and “runUMAP” functions. We compared a range of cluster resolutions for each data set using the clustree R package (Zappia and Oshlack, 2018) before selecting an optimal resolution for each (resolution = 3.0, 0.7, and 1.2, **Supplemental Figures 3 and 4**).

### Marker gene identification and sex-differential expression

Marker genes were identified per cluster for each data set using the Seurat “FindAllMarkers” function, using the Wilcox rank sum test (min.pct=0.25, logfc.threshold=0.25). For each gene, the expression in a given cluster was compared with expression in neurons in the remaining clusters.. Differentially expressed genes between the sexes per cluster were identified using the Seurat “FindAllMarkers” function, using the Wilcox rank sum test (min.pct=0.25, logfc.threshold=0.25). For each gene, the expression in a male neurons within a cluster was compared with expression in female neurons within the same cluster. The p-values for marker gene and sex-differential expression were adjusted for multiple testing using the Bonferroni method.

### Sex differences in cell number within clusters

A cluster was considered “sex-specific” if only cells from one sex were present in that cluster. Given the data set contains an unbalanced number of cells between the sexes in the full data set, the female cell numbers per cluster were divided by a scaling factor of 2.19. For a cluster to be considered “sex-biased”, cells from one sex were required to be >2-fold higher in that cluster. For a cluster to be considered “sex-biased, strong” the number of cells was >4-fold higher.

### Downsampling and random cell removal analyses

To perform random cell removal analyses, a random subset of 7,988 female cells (to equal the number of male cells) was selected by the “sample” R function. The Seurat “Dimplot” function was used to visualize these data sets and the number of cells per sex in each cluster was quantified. To perform random downsampling analyses, a random subset of 7,988 female cells were selected from the 17,530 filtered female cells using the “sample” R function, as above. The random subset of female cells was then merged with the Seurat object containing the 7,988 filtered male cells into a new Seurat object using the “merge” function. This was performed three times to create three independent random downsampled data sets. Each downsampled data set underwent the Seurat workflow described above. Consistent with our analysis of the full data set, we used the number of significant principal components (PC) up to where the first non-significant PC was met. This resulted in 99, 98, and 99 PCs. We chose cluster resolutions that yielded ∼113 clusters, the number resolved in the analysis of the full data set, resulting in resolutions of 3.0, 3.0, and 3.1. To match these down sampled clusters with those in the full data set, we used the R package clustifyr (Fu *et al*., 2020). The “seurat_ref” function was used to make a cluster reference from the full data set and the default “clustify” wrapper function was used to perform Spearman correlations between the reference downsampled analysis clusters (**Supplemental Table 3**).

### Immunostaining and microscopy

Adult and 48hr APF brains were dissected and imaged as previously described (Palmateer *et al*., 2021). Primary antibodies used were as follows: rat *α*-Fru^M^ (1:200) (Sanders and Arbeitman, 2008), rabbit *α*-Myc (1:6050; abcam, ab9106), rabbit *α*-Per (1:500) (gift from Michael Rosbash), mouse *α*-Nc82 (1:20, DSHB), mouse *α*-Prospero (1:100, DSHB). The secondary antibodies were as follows: goat *α*-rat 488 (1:1000, Invitrogen A11006), goat *α*-rabbit 568 (1:500; Invitrogen, A11036), rabbit *α*-GFP 488 (1:600; Invitrogen A21311), goat *α*-mouse 633 (1:500; Invitrogen, A21052). Both primary and secondary antibodies were diluted in TNT (0.1M Tris-HCL, 0.3M NaCl, 0.5% Triton X-100). All confocal microscopy was performed on a Zeiss LSM 700 system, with Zeiss Plan-Apochromat 20x/0.8 and 40x/1.4 objectives. The z-stack slice interval for all images was 1.0μm. A 1 Airy Unit (AU) pinhole size was calculated in Zeiss Zen software (Black edition, 2012) for each laser line: 488nm: 38μm; 555nm: 34μm; and 639nm: 39μm. All images were acquired at 1024×1024 pixel resolution with bidirectional scanning.

*Clk856* ∩ *fru P1* > *sm.GDP.Myc* and *Clk856* ∩ *fru P1* > *TrpA1^Myc^* cell bodies were scored blinded, in each brain hemisphere for 0-24hr adults and 4-7 day adults of both sexes (**Supplemental Table 7**). Data from both left and right hemispheres for each region within sex were pooled for analysis for each genotype and time point. Statistical testing within each quantified region was performed between the sexes within each genotype and time point using a Mann-Whitney U-test in GraphPad Prism (9.3.0).

### Subclustering male-specific/biased *dsx*-expressing neuron clusters

All male-specific or male-biased that were enriched for *dsx* expression (**Supplemental Table 4**), clusters 21, 47, and 68 in the full data analysis were subset and re-analyzed. Only *dsx* expressing neurons in these clusters were maintained in this analysis (*dsx* expression > 0 UMIs). 27 significant principal components were used to perform new dimensionality reduction based on Jackstraw analysis. Cluster resolution was determined based on visual inspection of distinct UMAP populations being identified as individual clusters (resolution = 0.5). Differential expression was calculated between the clusters using the same *FindAllMarkers* criteria as above.

### Subclustering circadian neuron clusters

All clusters classified as “circadian” in our full data set UMAP analysis were subset and PCA was performed only on these cells. The two significant principal components were used to perform new dimensionality reduction based on Jackstraw analysis. Cluster resolution was determined based on visual inspection of distinct UMAP populations being identified as individual clusters (resolution = 2.5). Differential expression was calculated between the clusters using the same *FindAllMarkers* criteria as above.

### Annotating mushroom body KC populations and subclustering

Cell clusters were annotated based on the presence genes that have been shown to be expressed in the mushroom body KCs and those specific to sub-populations. All of these genes were manually annotated for their presence (**Supplemental Table 4**), however, a cluster was only confidently annotated as a mushroom body KC subtype if at least two genes were enriched in expression in that cluster. For example, if a cluster had a marker gene with known general KC expression present (*ey* and/or *Dop1R2*) and additionally had *Fas2*, *sNPF*, and/or *trio* as a marker gene, the cluster was annotated as a KC subtype. These subtypes were determined based on the presence of *trio*, which shows high expression in only the γ KCs at 48hr APF (ref). αβ KCs were annotated based on marker genes for *ey* and/or *Dop1R2* with *sNPF* and/or *Fas2* in the absence of *trio*. Further, we also found clusters with *Fas2* and *sNPF* as marker genes with expression of *ey* or *Dop1R2* in the cluster, but not present as a marker gene. We also find one cluster per data set that has marker gene expression of *trio* in addition to one other KC-expressed gene, either *Fas2* or *Dop1R2*. Clusters with incomplete enriched marker gene expression but suggestive of being KC subtypes are annotated as lower confidence ab_KC clusters (indicated by *).

All high confidence KC clusters in our full data set UMAP analysis were subset and PCA was performed only on these cells. The 21 first significant principal components were used to perform new dimensionality reduction based on Jackstraw analysis. Cluster resolution was determined based on visual inspection of distinct UMAP populations being identified as individual clusters (resolution = 0.5). Differential expression was calculated between the clusters using the same *FindAllMarkers* criteria as above.

### Locomotor activity assays and analysis

Wild-type Canton S (CS), transgene controls without a *Gal4* driver (*w; UAS<stop<TrpA1^Myc^; fru^FLP^/+*), and experimental flies with expression restricted to *Clk856* ∩ *fru P1* neurons (*w; UAS<stop<TrpA1^Myc^/Clk856-Gal4; fru^FLP^/+*) were collected 0-6hrs post-eclosion and loaded into glass tubes that contained our standard laboratory food. Tubes containing the flies were loaded into *Drosophila* Activity Monitors (DAM) (Trikinetics, Waltham, MA). The light:dark (LD) assay was conducted at 25°C on a 12hr:12hr LD cycle and beam cross activity was recorded in one min. bins for 15 days. Incubator lights turned on at 8am and off at 8pm. Flies were aged to 5 days during the assay, therefore the first five days of data were removed from the analysis. This resulted in 10 days of data analyzed for all genotypes in the LD assay. In the dark:dark (DD) assay, flies were first entrained and aged at 25°C on a 12hr:12hr LD cycle for 7 days, allowing for 2 days of LD data for analysis. Next, we switched to a 12hr:12hr DD cycle for 10 days (constant darkness for 10 days). Beam cross activity for the DD assay was recorded in one-minute bins. Data were analyzed using ShinyR-DAM for both assays (Cichewicz and Hirsh, 2018). Dead flies were considered those with less than 50 beam cross events per day and were removed from the analyses, resulting in an n=26-30 for all genotypes in LD and n=23-30 for all genotypes in DD. ShinyR-DAM analysis of sleep was only performed on LD assay data. ShinyR-DAM measures sleep events using a 5-minute sliding window, where 5 minutes of inactivity is considered a sleep event. Statistical tests were performed on ShinyR-DAM output data in R Studio and GraphPad Prism (9.3.0). One-way ANOVA with Tukey-HSD *post hoc* testing was performed for comparisons across all genotypes. Comparisons between daytime and nighttime sleep were statistically compared using a student’s t-test for daytime vs. nighttime sleep. Circadian period analysis was performed in ShinyR-DAM using the DD assay data. The default ShinyR-DAM parameters were used as follows: Chi-Sq testing range of 18-30hrs, a Chi-Sq period testing resolution of 0.2, and a rhythmicity threshold for filtering arhythmic individuals (Qp.act/Qp.sig) of 1. All ShinyR-DAM output data presented is provided in **Supplemental Table 7**.

### Data availability

The Gene Expression Omnibus accession number for all scRNA-seq the data is: GSE160370

## Supporting information

All Supplemental Figures

## Acknowledgements

We thank the Florida State University College of Medicine Translation Core for running the sequencing core. Stocks were also obtained from the Bloomington Drosophila Stock Center (NIH P40OD018537). Antibodies used in this study were obtained from the Developmental Studies Hybridoma Bank, created by the NICHD of the NIH and maintained at The University of Iowa, Department of Biology, Iowa City, IA 52242.

## Supplemental Figure Legends

**Supplemental Figure 1. Data filtering and replicate overlap in UMAP space. (A)** Violin plots with the number of total expressed genes (nFeature_RNA), number of transcripts (nCount_RNA) and percentage of mitochondrial transcripts expressed (percent.mt) in each replicate from both sexes in the unfiltered data. MR1 and MR2 indicates male replicates 1 and 2. FR1 and FR2 indicates female replicates 1 and 2. **(B)** Violin plots of unfiltered cell data with replicates combined by sex. **(C)** Violin plots with the filtered cell data from each replicate for the number of expressed genes (nFeature_RNA), number of transcripts (nCount_RNA) and percentage of mitochondrial transcripts expressed (percent.mt). Each replicate was filtered for cells that met the following criteria: >200 nFeature_RNA,>4000 and >20,000 nCount_RNA and <5% percent.mt. **(D)** Violin plots of cells meeting filtering criteria data for each replicate combined by sex. **(E)** Table with total numbers of cells per replicate prior to filtering (unfiltered cells) and the number meeting filtering criteria (filtered cells). **(F)** Table with total numbers of cells per sex prior to filtering (unfiltered cells) and the number meeting filtering criteria (filtered cells). **(G)** UMAP plot of the male data set presented in **Figure 1D** with male cells labeled by replicate. Male replicate 1 and 2 are indicated (MR1 and MR2). **(H)** UMAP plot of the female data set presented in **Figure 1E** with female cells labeled by replicate. Female replicate 1 and 2 are indicated (FR1 and FR2). **(I)** UMAP plot of the full data set presented in **Figure 1F** with all cells labeled by replicate. (MR1, MR2, FR1 and FR2)

**Supplemental Figure 2. Jackstraw plots used to determine number of significant principal components to use for dimensional reduction and clustering. (A)** Jackstraw plot for the number of principal components (PCs) for the male data set. 80 principal components were used for the male data set **(B)** Jackstraw plot for the number of principal components (PCs) for the female data set. 104 principal components were used for the female data set. **(C)** Jackstraw plot for the number of principal components (PCs) for the full data set. 107 principal components were used for the full data set. For all three analyses, the number was chosen up to where the first non-significant principal component was identified.

**Supplemental Figure 3. Test of differing cluster resolutions for male and female *fru P1* neuron data sets. (A)** UMAP plot of 7,988 male *fru P1* neurons at cluster resolution of 3, defining 76 clusters, used throughout the study. **(B)** Relationship between male data set clusters at different cluster resolutions. This analysis was performed at three resolutions. The color of the circles indicates cluster resolution, with red circles indicating resolution of 1, green circles indicating resolution of 2, and blue circles indicating resolution of 3. **(C)** UMAP plot of 17,530 female *fru P1* neurons at cluster resolution of 0.7, defining 87 clusters, used throughout the study. **(D)** Relationship between female data set clusters at different cluster resolutions. This analysis was performed at three resolutions. The color of the circles indicates cluster resolution, with red circles indicating resolution of 0.2, green circles indicating resolution of 0.5, and blue circles indicating resolution of 0.7. For **(B)** and **(D)**, the shade of the color of the arrows between the different cluster resolutions represent the proportion of the cluster present at the next highest resolution (in_prop). The color of the arrow indicates the count of cells within each cluster, between resolutions (count). The size of the circle indicates the number of cells in each cluster (size).

**Supplemental Figure 4. Test of differing cluster resolutions for full *fru P1* neuron data set and cluster defining marker gene expression for all data sets. (A)** UMAP plot of 25,530 *fru P1* neurons at cluster resolution (res) of 0.5, 0.7, and 1.2, each defining a different number of clusters. In the study we used resolution of 1.2, defining 113 clusters. **(B)** Relationship between data set clusters at different cluster resolutions. This analysis was performed at three resolutions. The color of the circles indicates cluster resolution, with red circles indicating resolution of 0.5, green circles indicating resolution of 0.7, and blue circles indicating resolution of 1.2. The shade of the color of the arrows between the different cluster resolutions represent the proportion of the cluster present at the next highest resolution (in_prop). The color of the arrow indicates the count of cells within each cluster, between resolutions (count). The size of the circle indicates the number of cells in each cluster (size). **(C)** UMAP plot of 77 clusters of male *fru P1* neurons and heatmap showing scaled and log-normalized expression of top 10 cluster marker genes in male cell clusters. **(D)** UMAP plot of 88 clusters of female *fru P1* neurons and heatmap showing scaled and log-normalized expression of top 10 cluster marker genes in female cell clusters. **(E)** UMAP plot of 113 clusters from full *fru P1* neuron data set analysis and heatmap showing scaled and log-normalized expression of top 10 cluster marker genes. Marker genes for all analyses are listed in **Supplemental Table 4**.

**Supplemental Figure 5. Visualization of GFP, *fru*, *roX1* and *roX2* expression. (A)** Gene expression features plots of GFP and **(B)** *fru* expression in the full data set UMAP. *roX1* and *roX2* expression in the male data set UMAP **(C-D)**, female data set UMAP **(E-F)**, and full data set UMAP **(G-H)**. The gene expression feature plots show gene expression levels in purple in the UMAP, with color intensity proportional to the log normalized expression levels. **(I-J)** UMAP plot of the full data set after removal of *roX1* and *roX2* expression data, with all cells labeled by sex **(I)** and **(J)** replicate (see methods).

**Supplemental Figure 6. GO enrichment analysis for marker genes lists. (A-C)** Gene Ontology (GO) enrichment analysis for male, female, and full data set marker gene lists. The GO categories are (**A**) molecular function, (**B**) biological process, and (**C**) cellular component. The GO terms shown in the plots are the top ten most significantly enriched terms for each marker gene list (Benjamini-Hochberg, p<0.05). The size of each dot indicates the number of genes in our marker gene list divided by the total number of genes belonging to the GO category (GeneRatio). The color indicates the p value (p.adjust). All GO information is listed in **Supplemental Table 5.**

**Supplemental Figure 7. Random removal and downsampling of female cells. (A)** From the full data set UMAP, a random set of 7,988 female cells were kept and the clusters were analyzed. This was performed three independent times. **(B)** A full reanalysis, using the standard Sueurat workflow, of the filtered cells was performed three times after downsampling the female data to 7,988 female cells. The UMAPs are shown. Comparisons across clusters are in Supplemental Table 3 (**A and B**).

**Supplemental Figure 8. Gene expression in sex-specific and sex-biased clusters**. (**A**) Gene expression feature plots showing gene expression in neurons from female-specific cluster 107. *CCHa2* and genes with known expression in the VNC are shown. (**B**) Feature plots showing gene expression in neurons from male-specific cluster 68. *dsx* and genes with known expression in the VNC are shown. **(C)** The full data set UMAP plot showing *dsx*-expressing clusters that are male-specific (dark blue), strongly male-biased (purple), or male-biased (light blue) in cell number. (**D**) A UMAP showing the clustering of five *dsx* subclusters, with the subclusters colorized by original sex-biased or sex-specific cluster assignment shown in (**C**). (**E-H**) Gene expression feature plots for *dsx* subclusters showing genes with known expression in the VNC. The gene expression feature plots show gene expression levels in purple in the UMAP, with color intensity proportional to the log normalized expression levels. The color legend is to the right in each panel.

**Supplemental Figure 9. VNC cluster annotations based on *Hox* expression for male and female data sets. (A-H)** Gene expression feature plots showing expression of *Hox* genes that have known expression in the VNC, in male UMAP (**A-D**) and female UMAP (**E-H**) data sets. **(I-J)** UMAP plot showing clusters annotated as having neurons from the VNC (highlight in green), in the male (**I**) and female (**J**) data set analyses. The cluster number is shown and corresponds to the annotation in **Supplemental Table 4**. **(K-L)** Pearson correlation heatmap of *Hox* gene expression, at the single neuron level, in the male (**K**) and female (**L**) data set analyses. Pearson’s r value denoted on heatmap and colorized according to legend (right). The gene expression feature plots show gene expression levels in purple in the UMAP, with color intensity proportional to the log normalized expression levels. The color legend is to the right in each panel.

**Supplemental Figure 10. Mushroom body Kenyon cell annotations. (A)** Schematic of criteria defining mushroom body Kenyon cell (KC) subtypes, based on gene expression of *ey*, *Dop1R2*, *sNPF*, *Fas2*, and *trio*. **(B-C)** Gene expression feature plots showing expression of *ey* (**B**) and *Dop1R2* (**C**) in KC annotated clusters from the full data set analysis. (**D**) UMAP plot showing Kenyon cell (KC) cluster annotations in the male data set analysis. The green and mustard highlights mushroom body clusters, and grey (N) indicates all other clusters (**Supplemental Table 4**). **(E-F)** Gene expression feature plots showing expression of *ey* (**E**) and *Dop1R2* (**F**) in KC annotated clusters from the male data set analyses. (**G**) UMAP plot showing Kenyon cell (KC) cluster annotations in the female data set analysis. The green and mustard highlights mushroom body clusters, and grey (N) indicates all other clusters (**Supplemental Table 4**). **(H-I)** Gene expression feature plots showing expression of *ey* (**H**) and *Dop1R2* (**I**) in KC annotated clusters from the female data set analyses. The gene expression feature plots show gene expression levels in purple in the UMAP, with color intensity proportional to the log normalized expression levels. The color legend is to the right in each panel. (**J**) UMAP of 13 mushroom body subclusters, with the subclusters colorized by original full data set analysis cluster assignment, shown in **Figure 4A** (legend on right).

**Supplemental Figure 11. Expression of genes consistent with production of fast-acting neurotransmitters (FAN). (A)** UMAP plot showing clusters annotated as producing FANs, in the male data set (**Supplemental Table 4**). Color legend on right indicate clusters with FAN marker gene expression, or no expression (grey, N). **(B)** Size proportional Euler diagram showing the percentage of cells with overlapping expression of genes indicative of cholinergic (*VAChT*), GABAergic (*Gad1*), and glutamatergic (*VGlut*) neurons in the male data set. **(C-G)** Gene expression feature plots showing distribution of neurons expressing genes indicative of FAN production: acetylcholine (*ChAT* and *VAChT*), GABA (*Gad1* and *VGAT*), and glutamate (*VGlut*), in the male data set. **(H)** UMAP plot showing clusters annotated as producing FANs, in the female data set (**Supplemental Table 4**). Color legend on right indicate clusters with FAN marker gene expression, or no expression (grey, N). **(I)** Size proportional Euler diagram showing the percentage of cells with overlapping expression of genes indicative of cholinergic (*VAChT*), GABAergic (*Gad1*), and glutamatergic (*VGlut*) neurons in the male data set. **(J-N)** Gene expression feature plots showing distribution of neurons expressing genes indicative of FAN production: acetylcholine (*ChAT* and *VAChT*), GABA (*Gad1* and *VGAT*), and glutamate (*VGlut*), in the female data set. The gene expression feature plots show gene expression levels in purple in the UMAP, with color intensity proportional to the log normalized expression levels. The color legend is to the right in each panel.

**Supplemental Figure 12. Expression of genes consistent with production of aminergic neurotransmitters.** (**A-C**) Gene expression feature plots for (**A**) full, (**B**) male and (**C**) female data set UMAPs showing gene expression of *Vmat* (aminergic neurons) and biosynthetic enzymes or transporters for specific aminergic transmitters: dopamine (*DAT* and *ple*), serotonin (*SerT*), and octopamine (*Tdc2*). **(D-G)** Confocal maximum projections for *Vmat* ∩ *fru P1* co-expressing neurons from males and females. Brain is on left and VNC is on right in each panel. (**D-E**) The images are from animals at 48hr APF and (**F-G**) 0-24hr adults. Scale bars = 50 μm. (**H-I**) UMAP plot showing aminergic cluster annotations in the male (**H**) and female (**I**) data sets (**Supplemental Table 4**). Color legend on right indicate clusters with aminergic marker gene expression, or no expression (grey, N). (**J**) Gene expression feature plots for dopaminergic neurons showing expression of *DAT, ple,* and *Abd-B*, in the full data set. (**K**) Gene expression feature plots for dopaminergic neurons showing expression of *Ubx* and *Abd-B*, in the male data set UMAP. The gene expression feature plots show gene expression levels in purple in the UMAP, with color intensity proportional to the log normalized expression levels. (**L**) Dot plots of top 5 expressed genes in dopaminergic clusters, based on log fold-change in gene expression, in the male data set. Dot size indicates the percentage of neurons expressing each gene per cluster (Percent Expressed). Average normalized expression level is shown by color intensity (Average Expression).

**Supplemental Figure 13. Visualization of circadian clock gene expression.** (**A-C**) Gene expression feature plots for full (**A**), male (**B**) and female (**C**) data set UMAPs showing expression of circadian genes (*per*, *tim*, *Clk*, *vri*, and *Pdp1*). The gene expression feature plots show gene expression levels in purple in the UMAP, with color intensity proportional to the log normalized expression levels. (**D**) Dot plot of expression levels of genes with functions in the circadian molecular clock, in the circadian subclusters. Dot size indicates the percentage of neurons expressing each gene per cluster (Percent Expressed). Average normalized expression level is shown by color intensity (Average Expression). (**E-E”**) Confocal maximum projections of brains immunostained for Per (magenta) and *fru P1* >*mcd8::GFP* neurons, from 48hr APF female pupae, at 40x magnification. The left brain hemisphere is shown. (**F-I**) Confocal maximum projections of *Clk856 ∩ fru P1* neurons in 4-7 day adult male (**F-G**) and female (H-I) brains and VNCs with *UAS<stop<smGDP.myc* transgene. Additional brains in **G** and **I** show the variation in expression. (**J-K**) Confocal maximum projections of *Clk856 ∩ fru P1* neurons expressing *UAS<stop<* TrpA1^Myc^, immunostained for Myc, in 4-7 day adult male (**F-G**) and female (**H-I**) brains and VNCs. Scale bars = 50 μm.

**Supplemental Figure 14. Activating *Clk856* ∩ *fru P1* neurons effects locomotor activity and sleep in LD**. (**A**) Boxplots showing average locomotor activity per individual (beam crosses), per day, for each genotype and sex (n=26-30). The top and bottom of the box in the boxplot indicate 25^th^ and 75^th^ percentiles. (**B-F**) Average total sleep (B), daytime (C), and nighttime sleep (D) for each genotype and sex. (E-F) Average sleep bout length (E) and sleep bout number (F) for each genotype and sex. Average sleep was calculated in ShinyRDAM (Cichewicz and Hirsh, 2018). (G) Average locomotor activity during the day (lights on) and night (lights off), for each genotype and sex. (**H-I**) Actograms showing average locomotor activity, over 24 hours, during day six of the 10 days analyzed for males (**H**) and females (**I**). Lights on is at 8hrs (ZT0) and lights off is at 20hrs (ZT12), in the light-dark cycle. The genotypes of the flies assayed are: Canton S (CS), *w; UAS<stop<TrpA1^Myc^; fru^FLP^/+* (*TrpA1 fruFLP*), and *w; UAS<stop<TrpA1^Myc^/Clk856-Gal4; fru^FLP^/+* (*Clk856 TrpA1 fruFLP*). Sex is indicated after genotype by “M” for male or “F” for female. One-way ANOVA with Tukey-HSD post hoc tests were used for statistical analyses for comparisons across genotype in panels **A-F.** Student’s t-tests were used to compare daytime vs. nighttime activity in panel **G**. *: p<0.05 *, p<0.01 **, p<0.001 ***.

**Supplemental Figure 15. Activating *Clk856* ∩ *fru P1* neurons impacts circadian period length and strength in DD**. (**A-B**) Actograms showing average locomotor activity across all flies for each genotype, over 24 hours, during two days of LD followed by 10 days of DD for males (**A**) and females (**B**). Lights on is at 8hrs (ZT0) and lights off is at 20hrs (ZT12), in the light-dark cycle. (**C**) Chi-square periodogram for all genotypes. Lines above and below the dotted line indicate standard error of the mean (SEM). (**D**) Circadian period peaks for individual flies of each genotype (indicated by a dot). (E) Circadian period strength for individual flies of each genotype. The genotypes of the flies assayed are: Canton S (CS), *w; UAS<stop<TrpA1^Myc^; fru^FLP^/+* (*TrpA1 fruFLP*), and *w; UAS<stop<TrpA1^Myc^/Clk856-Gal4; fru^FLP^/+* (*Clk856 TrpA1 fruFLP*). Sex is indicated after genotype by “M” for male or “F” for female.

**Supplemental Figure 16. Annotation of neurons that express neuropeptides in males and females.** (**A-B**) UMAP plot showing annotated clusters with neurons that produce neuropeptides with known roles in *fru P1* neurons for directing reproductive behavior, in the male data set UMAP **(A)** and female data set UMAP **(B)** (**Supplemental Table 4**). (**C-N**) Brain and VNC confocal maximum projections showing intersecting expression patterns (in green), for males and females, as indicated. The patterns of *Dh44* ∩ *fru P1*, *CrzR* ∩ *fru P1*, and *Lgr3* ∩ *fru P1* neurons are shown. Scale bars = 50 μm.

**Supplemental Figure 17. *Wnt* ∩ *fru P1* expression patterns.** (**A-J**) Brain and VNC confocal maximum projections showing intersecting expression patterns (in green), for males and females, as indicated. The patterns of *Wnt4* ∩ *fru P1*, *Wnt10* ∩ *fru P1*, and *Wnt5* ∩ *fru P1* neurons are shown. Scale bars = 50 μm. The *Wnt5* stock was not available to perform expression analyses at 48hr APF due to the stock being unhealthy.

**Supplemental Figure 18. IgSF gene expression in *fru P1* neurons.** (**A**) Heatmap of log-normalized gene expression for IgSF encoding genes. Gene expression values were mean-scaled and log-normalized.

**Supplemental Figure 19: Sex differences in *dpr*/*DIP* gene expression in *fru P1* neurons** (**A**) Dot plot split by sex for all *dpr* and *DIP* genes. Dot size indicates the percentage of neurons expressing each gene per cluster (Percent Expressed). Male expression is shown in blue and female expression is shown in pink. Black circles indicate clusters where *dpr/DIP* genes are sex-differentially expressed (**Supplemental Table 9**).

**Supplemental Figure 20: Homeobox-like domain superfamily TF expression in *fru P1* neurons.** (**A**) Dot plot of homeobox-like domain superfamily TF encoding marker gene expression in full data set analysis clusters. Dot size indicates the percentage of neurons expressing each gene per cluster (Percent Expressed). Average normalized expression level is shown by color intensity (Average Expression).

**Supplemental Figure 21: Zinc finger C2H2 TF expression in *fru P1* neurons. A**) Dot plot of homeobox domain TF encoding marker gene expression in full data set analysis clusters. Dot size indicates the percentage of neurons expressing each gene per cluster (Percent Expressed). Average normalized expression level is shown by color intensity (Average Expression).

**Supplemental Figure 22: Helix-loop-helix DNA binding domain superfamily expression in *fru P1* neurons. A**) Dot plot of helix-loop-helix (HLH) DNA binding domain TF encoding marker gene expression in full data set analysis clusters. Dot size indicates the percentage of neurons expressing each gene per cluster (Percent Expressed). Average normalized expression level is shown by color intensity (Average Expression).

## Supplemental Tables

**Supplemental Table 1. Sequencing metrics and barcode-rank plots for all replicates.**

Excel data table containing CellRanger barcode-rank plots and sequencing metrics for all replicates. After quality control filtering, the median number of genes per cell and median UMI counts per cell are presented.

**Supplemental Table 2. Distribution of cells per replicate and sex in UMAPs and select gene expression metrics.**

Excel data table containing cell number contributions of each individual replicate to UMAP clusters. This table also contains the number of cells and percent of total cells expressing *GFP*, *fru*, *roX1*, *roX2*, *elav*, *nSyb*, and *noe*.

**Supplemental Table 3. Assigning sex-bias to clusters in full and downsampled data sets.**

Excel data table containing cell number contributions by each sex per cluster. A scaling factor to normalize the number of male and female cells was applied to determine the sex-bias in cell number. The table includes assessments of cluster data with random female cells removed, and a full Seurat re-analyses of data with downsampled female cell numbers. The table includes the cluster comparisons of downsampled analyses using clustifyr matching.

**Supplemental Table 4. Marker genes and annotations.**

Excel data table containing marker gene lists and cluster annotations for all analyses. Statistical tests of marker gene list overlaps with lists of genes shown to be enriched or differentially expressed at 48hr APF in previous *fru P1* TRAP study (Palmateer *et al*., 2021).

**Supplemental Table 5. All Gene Ontology (GO) enrichment results.**

Excel data table containing GO enrichment results.

**Supplemental Table 6. Marker genes for subclustering analyses.**

Excel data table containing marker genes for *dsx*, Kenyon cell, and circadian clock neuron subclustering.

**Supplemental Table 7. Clk856 ∩ *fru P1* cell body counts in the adult brain and Drosophila activity monitor (DAM) data.**

Excel data table containing the number of *Clk856* ∩ *fru P1* cell bodies per region in adult male and female brains. The DAM data graphs and plots are showing in Supplemental Figure 14.

**Supplemental Table 8. *nAChR* expression.**

Excel data table containing the proportion of total neurons expressing *nAChR*s in all data sets.

**Supplemental Table 9. Sex differences in gene expression within cluster.**

Excel data table containing the differentially expressed genes between the sexes per cluster.

