## Supplementary material for "Single-cell transcriptome profiles of *Drosophila fruitless*-expressing neurons from both sexes": All Supplemental Figures

### Unfiltered Data

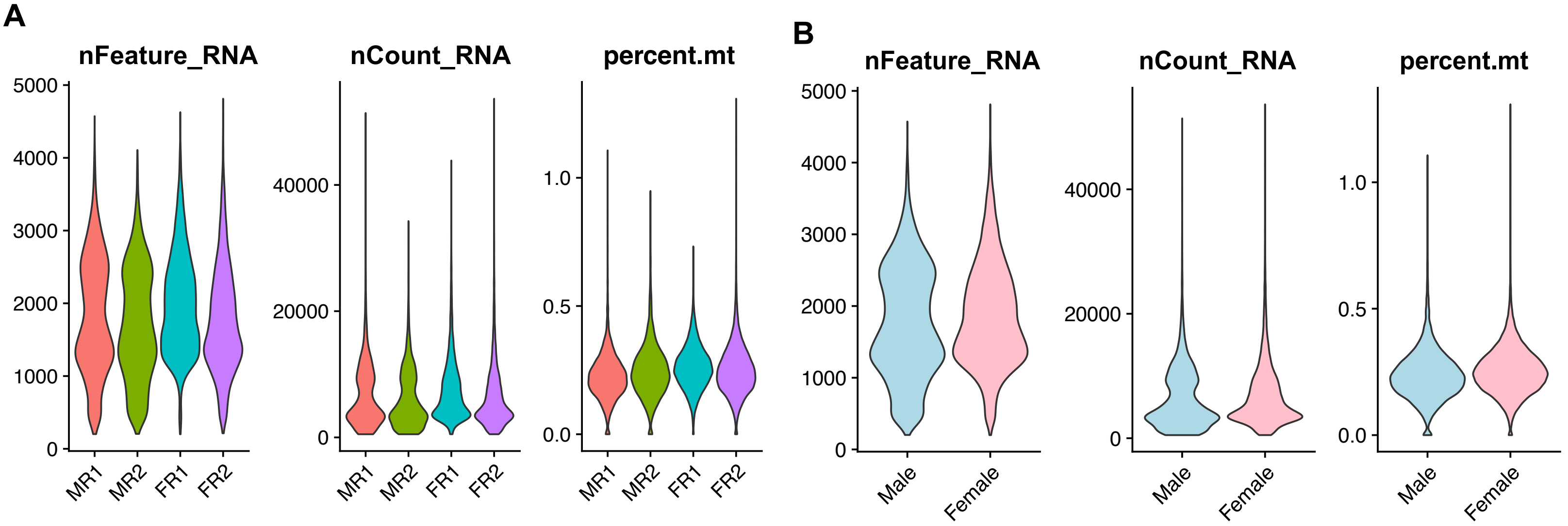


### Filtered Data

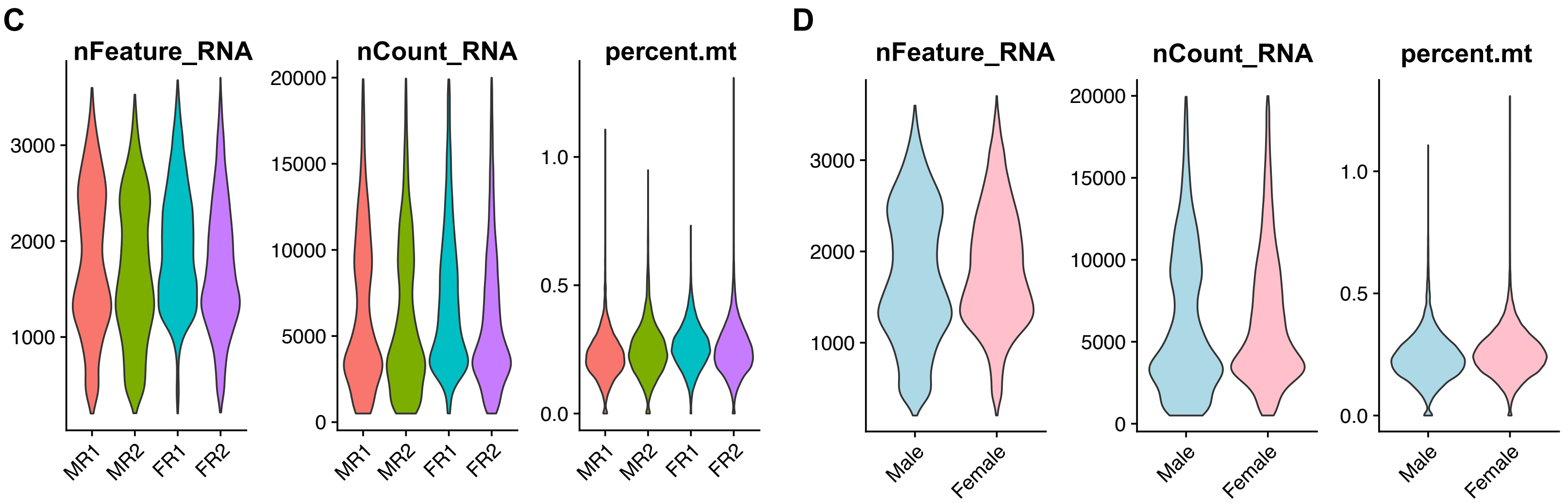


**E**

| Replicate | Unfiltered Cells | Filtered Cells |
|---|---|---|
| MR1 | 3,715 | 3,644 |
| MR2 | 4,388 | 4,344 |
| FR1 | 7,863 | 7,620 |
| FR2 | 10,149 | 9,910 |

**F**

| Sex | Unfiltered Cells | Filtered Cells |
|---|---|---|
| Male | 8,103 | 7,988 |
| Female | 18,012 | 17,530 |

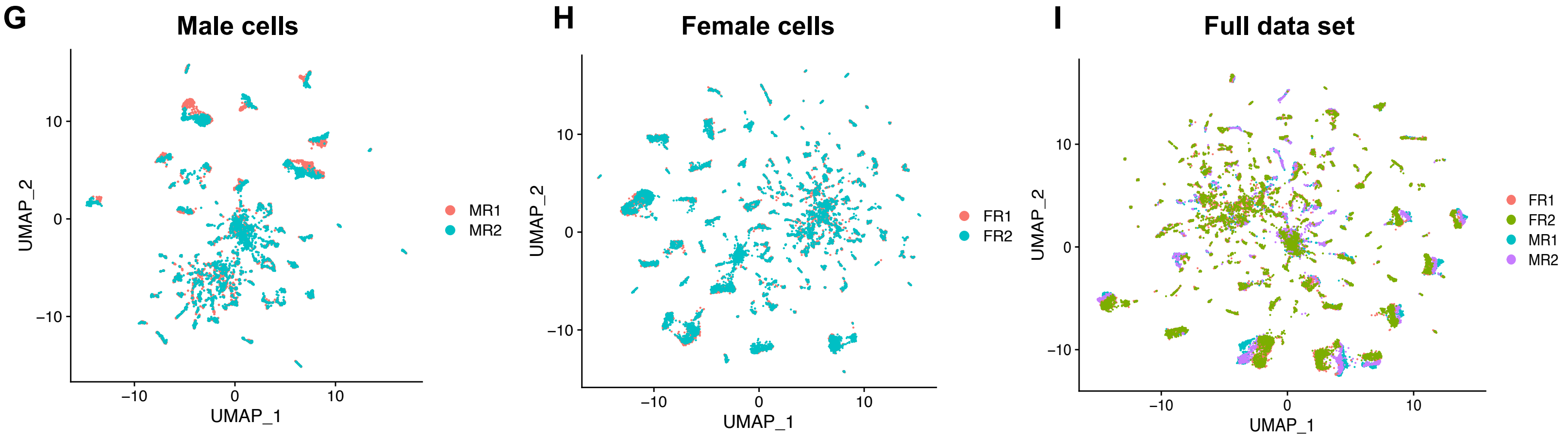

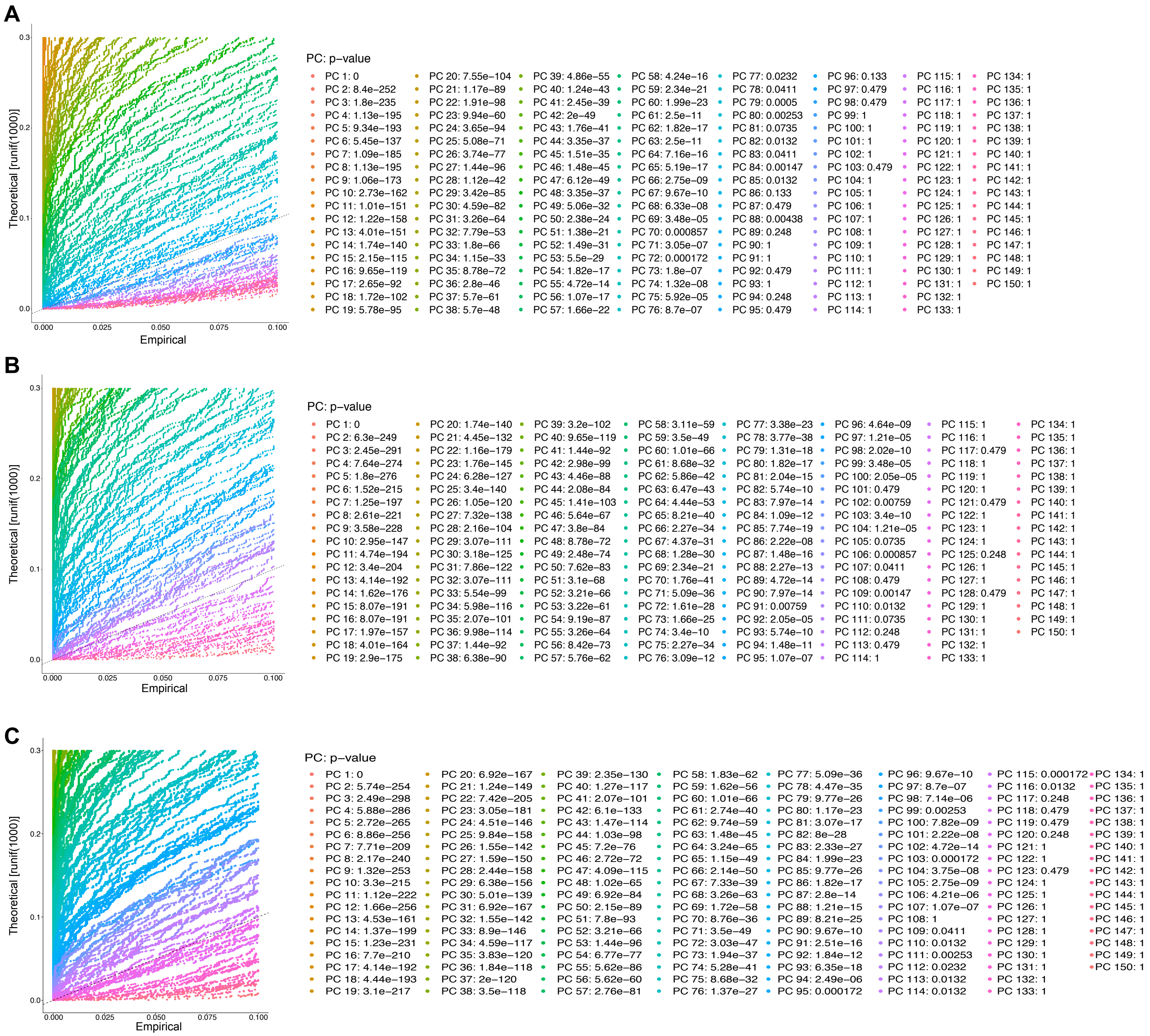

A
Theoretical [runif(1000)]
Empirical
0.0
0.1
0.2
0.3
0.000
0.025
0.050
0.075
0.100
PC: p−value
PC 1: 0
PC 2: 8.4e−252
PC 3: 1.8e−235
PC 4: 1.13e−195
PC 5: 9.34e−193
PC 6: 5.45e−137
PC 7: 1.09e−185
PC 8: 1.13e−195
PC 9: 1.06e−173
PC 10: 2.73e−162
PC 11: 1.01e−151
PC 12: 1.22e−158
PC 13: 4.01e−151
PC 14: 1.74e−140
PC 15: 2.15e−115
PC 16: 9.65e−119
PC 17: 2.65e−92
PC 18: 1.72e−102
PC 19: 5.78e−95
PC 20: 7.55e−104
PC 21: 1.17e−89
PC 22: 1.91e−98
PC 23: 9.94e−60
PC 24: 3.65e−94
PC 25: 5.08e−71
PC 26: 3.74e−77
PC 27: 1.44e−96
PC 28: 1.12e−42
PC 29: 3.42e−85
PC 30: 4.59e−82
PC 31: 3.26e−64
PC 32: 7.79e−53
PC 33: 1.8e−66
PC 34: 1.15e−33
PC 35: 8.78e−72
PC 36: 2.8e−46
PC 37: 5.7e−61
PC 38: 5.7e−48
PC 39: 4.86e−55
PC 40: 1.24e−43
PC 41: 2.45e−39
PC 42: 2e−49
PC 43: 1.76e−41
PC 44: 3.35e−37
PC 45: 1.51e−35
PC 46: 1.48e−45
PC 47: 6.12e−49
PC 48: 3.35e−37
PC 49: 5.06e−32
PC 50: 2.38e−24
PC 51: 1.38e−21
PC 52: 1.49e−31
PC 53: 5.5e−29
PC 54: 1.82e−17
PC 55: 4.72e−14
PC 56: 1.07e−17
PC 57: 1.66e−22
PC 58: 4.24e−16
PC 59: 2.34e−21
PC 60: 1.99e−23
PC 61: 2.5e−11
PC 62: 1.82e−17
PC 63: 2.5e−11
PC 64: 7.16e−16
PC 65: 5.19e−17
PC 66: 2.75e−09
PC 67: 9.67e−10
PC 68: 6.33e−08
PC 69: 3.48e−05
PC 70: 0.000857
PC 71: 3.05e−07
PC 72: 0.000172
PC 73: 1.8e−07
PC 74: 1.32e−08
PC 75: 5.92e−05
PC 76: 8.7e−07
PC 77: 0.0232
PC 78: 0.0411
PC 79: 0.0005
PC 80: 0.00253
PC 81: 0.0735
PC 82: 0.0132
PC 83: 0.0411
PC 84: 0.00147
PC 85: 0.0132
PC 86: 0.133
PC 87: 0.479
PC 88: 0.00438
PC 89: 0.248
PC 90: 1
PC 91: 1
PC 92: 0.479
PC 93: 1
PC 94: 0.248
PC 95: 0.479
PC 96: 0.133
PC 97: 0.479
PC 98: 0.479
PC 99: 1
PC 100: 1
PC 101: 1
PC 102: 1
PC 103: 0.479
PC 104: 1
PC 105: 1
PC 106: 1
PC 107: 1
PC 108: 1
PC 109: 1
PC 110: 1
PC 111: 1
PC 112: 1
PC 113: 1
PC 114: 1
PC 115: 1
PC 116: 1
PC 117: 1
PC 118: 1
PC 119: 1
PC 120: 1
PC 121: 1
PC 122: 1
PC 123: 1
PC 124: 1
PC 125: 1
PC 126: 1
PC 127: 1
PC 128: 1
PC 129: 1
PC 130: 1
PC 131: 1
PC 132: 1
PC 133: 1
PC 134: 1
PC 135: 1
PC 136: 1
PC 137: 1
PC 138: 1
PC 139: 1
PC 140: 1
PC 141: 1
PC 142: 1
PC 143: 1
PC 144: 1
PC 145: 1
PC 146: 1
PC 147: 1
PC 148: 1
PC 149: 1
PC 150: 1
B
Theoretical [runif(1000)]
Empirical
0.0
0.1
0.2
0.3
0.000
0.025
0.050
0.075
0.100
PC: p−value
PC 1: 0
PC 2: 6.3e−249
PC 3: 2.45e−291
PC 4: 7.64e−274
PC 5: 1.8e−276
PC 6: 1.52e−215
PC 7: 1.25e−197
PC 8: 2.61e−221
PC 9: 3.58e−228
PC 10: 2.95e−147
PC 11: 4.74e−194
PC 12: 3.4e−204
PC 13: 4.14e−192
PC 14: 1.62e−176
PC 15: 8.07e−191
PC 16: 8.07e−191
PC 17: 1.97e−157
PC 18: 4.01e−164
PC 19: 2.9e−175
PC 20: 1.74e−140
PC 21: 4.45e−132
PC 22: 1.16e−179
PC 23: 1.76e−145
PC 24: 6.28e−127
PC 25: 3.4e−140
PC 26: 1.05e−120
PC 27: 7.32e−138
PC 28: 2.16e−104
PC 29: 3.07e−111
PC 30: 3.18e−125
PC 31: 7.86e−122
PC 32: 3.07e−111
PC 33: 5.54e−99
PC 34: 5.98e−116
PC 35: 2.07e−101
PC 36: 9.98e−114
PC 37: 1.44e−92
PC 38: 6.38e−90
PC 39: 3.2e−102
PC 40: 9.65e−119
PC 41: 1.44e−92
PC 42: 2.98e−99
PC 43: 4.46e−88
PC 44: 2.08e−84
PC 45: 1.41e−103
PC 46: 5.64e−67
PC 47: 3.8e−84
PC 48: 8.78e−72
PC 49: 2.48e−74
PC 50: 7.62e−83
PC 51: 3.1e−68
PC 52: 3.21e−66
PC 53: 3.22e−61
PC 54: 9.19e−87
PC 55: 3.26e−64
PC 56: 8.42e−73
PC 57: 5.76e−62
PC 58: 3.11e−59
PC 59: 3.5e−49
PC 60: 1.01e−66
PC 61: 8.68e−32
PC 62: 5.86e−42
PC 63: 6.47e−43
PC 64: 4.44e−53
PC 65: 8.21e−40
PC 66: 2.27e−34
PC 67: 4.37e−31
PC 68: 1.28e−30
PC 69: 2.34e−21
PC 70: 1.76e−41
PC 71: 5.09e−36
PC 72: 1.61e−28
PC 73: 1.66e−25
PC 74: 3.4e−10
PC 75: 2.27e−34
PC 76: 3.09e−12
PC 77: 3.38e−23
PC 78: 3.77e−38
PC 79: 1.31e−18
PC 80: 1.82e−17
PC 81: 2.04e−15
PC 82: 5.74e−10
PC 83: 7.97e−14
PC 84: 1.09e−12
PC 85: 7.74e−19
PC 86: 2.22e−08
PC 87: 1.48e−16
PC 88: 2.27e−13
PC 89: 4.72e−14
PC 90: 7.97e−14
PC 91: 0.00759
PC 92: 2.05e−05
PC 93: 5.74e−10
PC 94: 1.48e−11
PC 95: 1.07e−07
PC 96: 4.64e−09
PC 97: 1.21e−05
PC 98: 2.02e−10
PC 99: 3.48e−05
PC 100: 2.05e−05
PC 101: 0.479
PC 102: 0.00759
PC 103: 3.4e−10
PC 104: 1.21e−05
PC 105: 0.0735
PC 106: 0.000857
PC 107: 0.0411
PC 108: 0.479
PC 109: 0.00147
PC 110: 0.0132
PC 111: 0.0735
PC 112: 0.248
PC 113: 0.479
PC 114: 1
PC 115: 1
PC 116: 1
PC 117: 0.479
PC 118: 1
PC 119: 1
PC 120: 1
PC 121: 0.479
PC 122: 1
PC 123: 1
PC 124: 1
PC 125: 0.248
PC 126: 1
PC 127: 1
PC 128: 0.479
PC 129: 1
PC 130: 1
PC 131: 1
PC 132: 1
PC 133: 1
PC 134: 1
PC 135: 1
PC 136: 1
PC 137: 1
PC 138: 1
PC 139: 1
PC 140: 1
PC 141: 1
PC 142: 1
PC 143: 1
PC 144: 1
PC 145: 1
PC 146: 1
PC 147: 1
PC 148: 1
PC 149: 1
PC 150: 1
C
Theoretical [runif(1000)]
Empirical
0.0
0.1
0.2
0.3
0.000
0.025
0.050
0.075
0.100
PC: p−value
PC 1: 0
PC 2: 5.74e−254
PC 3: 2.49e−298
PC 4: 5.88e−286
PC 5: 2.72e−265
PC 6: 8.86e−256
PC 7: 7.71e−209
PC 8: 2.17e−240
PC 9: 1.32e−253
PC 10: 3.3e−215
PC 11: 1.12e−222
PC 12: 1.66e−256
PC 13: 4.53e−161
PC 14: 1.37e−199
PC 15: 1.23e−231
PC 16: 7.7e−210
PC 17: 4.14e−192
PC 18: 4.44e−193
PC 19: 3.1e−217
PC 20: 6.92e−167
PC 21: 1.24e−149
PC 22: 7.42e−205
PC 23: 3.05e−181
PC 24: 4.51e−146
PC 25: 9.84e−158
PC 26: 1.55e−142
PC 27: 1.59e−150
PC 28: 2.44e−158
PC 29: 6.38e−156
PC 30: 5.01e−139
PC 31: 6.92e−167
PC 32: 1.55e−142
PC 33: 8.9e−146
PC 34: 4.59e−117
PC 35: 3.83e−120
PC 36: 1.84e−118
PC 37: 2e−120
PC 38: 3.5e−118
PC 39: 2.35e−130
PC 40: 1.27e−117
PC 41: 2.07e−101
PC 42: 6.1e−133
PC 43: 1.47e−114
PC 44: 1.03e−98
PC 45: 7.2e−76
PC 46: 2.72e−72
PC 47: 4.09e−115
PC 48: 1.02e−65
PC 49: 6.92e−84
PC 50: 2.15e−89
PC 51: 7.8e−93
PC 52: 3.21e−66
PC 53: 1.44e−96
PC 54: 6.77e−77
PC 55: 5.62e−86
PC 56: 5.62e−60
PC 57: 2.76e−81
PC 58: 1.83e−62
PC 59: 1.62e−56
PC 60: 1.01e−66
PC 61: 2.74e−40
PC 62: 9.74e−59
PC 63: 1.48e−45
PC 64: 3.24e−65
PC 65: 1.15e−49
PC 66: 2.14e−50
PC 67: 7.33e−39
PC 68: 3.26e−63
PC 69: 1.72e−58
PC 70: 8.76e−36
PC 71: 3.5e−49
PC 72: 3.03e−47
PC 73: 1.94e−37
PC 74: 5.28e−41
PC 75: 8.68e−32
PC 76: 1.37e−27
PC 77: 5.09e−36
PC 78: 4.47e−35
PC 79: 9.77e−26
PC 80: 1.17e−23
PC 81: 3.07e−17
PC 82: 8e−28
PC 83: 2.33e−27
PC 84: 1.99e−23
PC 85: 9.77e−26
PC 86: 1.82e−17
PC 87: 2.8e−14
PC 88: 1.21e−15
PC 89: 8.21e−25
PC 90: 9.67e−10
PC 91: 2.51e−16
PC 92: 1.84e−12
PC 93: 6.35e−18
PC 94: 2.49e−06
PC 95: 0.000172
PC 96: 9.67e−10
PC 97: 8.7e−07
PC 98: 7.14e−06
PC 99: 0.00253
PC 100: 7.82e−09
PC 101: 2.22e−08
PC 102: 4.72e−14
PC 103: 0.000172
PC 104: 3.75e−08
PC 105: 2.75e−09
PC 106: 4.21e−06
PC 107: 1.07e−07
PC 108: 1
PC 109: 0.0411
PC 110: 0.0132
PC 111: 0.00253
PC 112: 0.0232
PC 113: 0.0132
PC 114: 0.0132
PC 115: 0.000172
PC 116: 0.0132
PC 117: 0.248
PC 118: 0.479
PC 119: 0.479
PC 120: 0.248
PC 121: 1
PC 122: 1
PC 123: 0.479
PC 124: 1
PC 125: 1
PC 126: 1
PC 127: 1
PC 128: 1
PC 129: 1
PC 130: 1
PC 131: 1
PC 132: 1
PC 133: 1
PC 134: 1
PC 135: 1
PC 136: 1
PC 137: 1
PC 138: 1
PC 139: 1
PC 140: 1
PC 141: 1
PC 142: 1
PC 143: 1
PC 144: 1
PC 145: 1
PC 146: 1
PC 147: 1
PC 148: 1
PC 149: 1
PC 150: 1

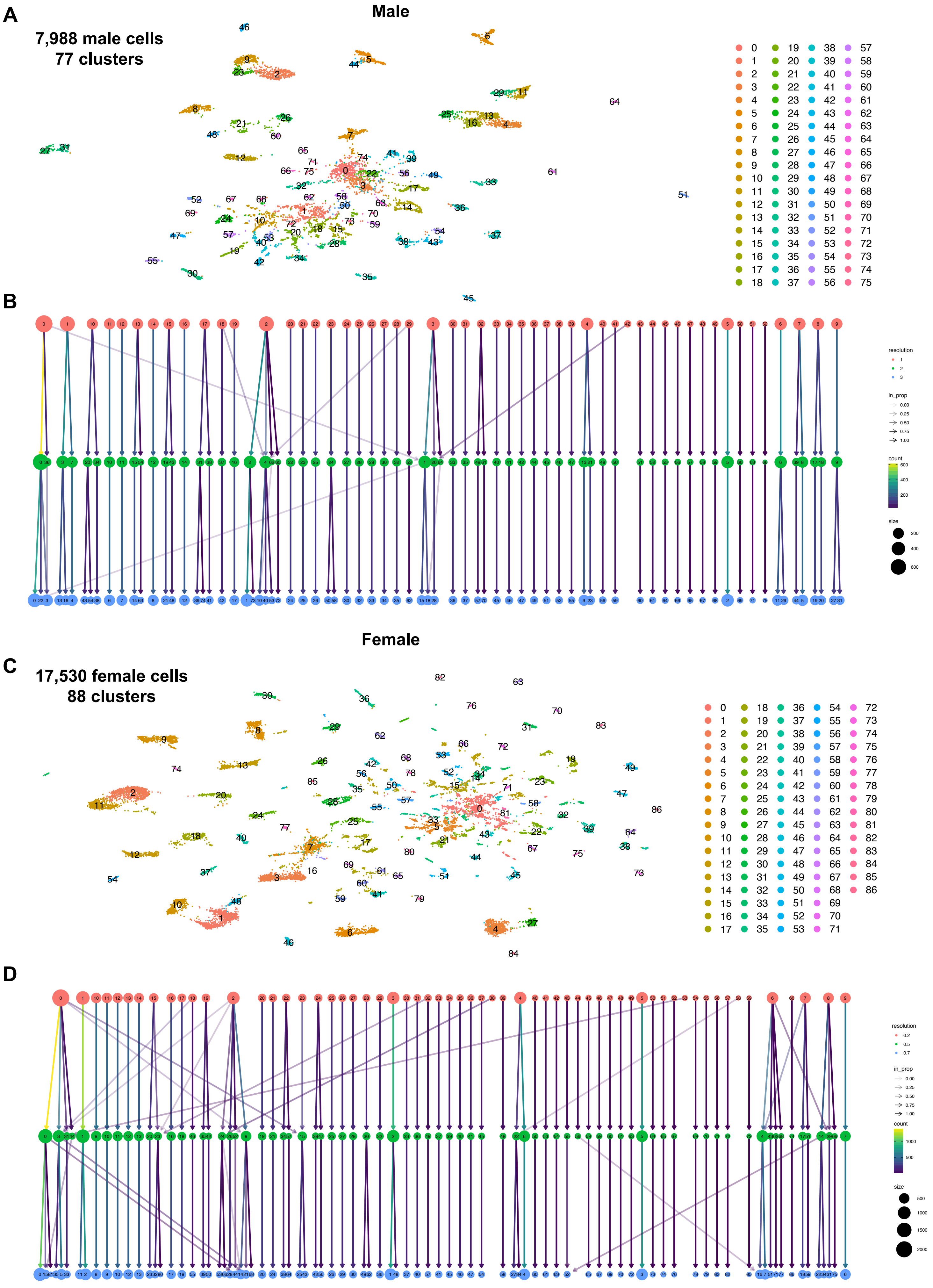
A
Male
7,988 male cells
77 clusters
B
resolution
in_prop
count
size
Female
C
17,530 female cells
88 clusters
D
resolution
in_prop
count
size

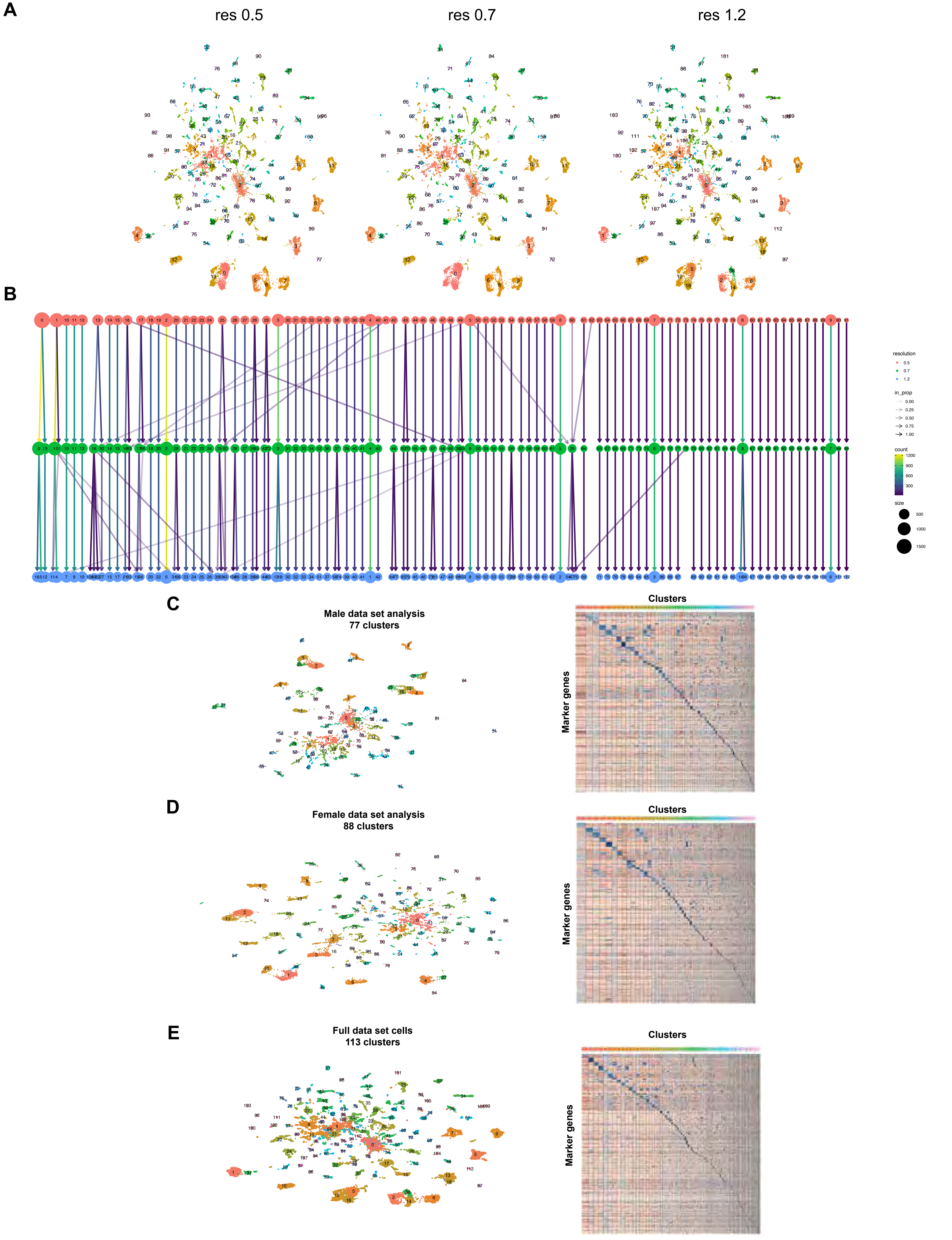
A
res 0.5
res 0.7
res 1.2
B
resolution
0.5
0.7
1.2
in_prop
0.00
0.25
0.50
0.75
1.00
count
1200
900
600
300
size
500
1000
1500
C
Male data set analysis
77 clusters
Clusters
Marker genes
D
Female data set analysis
88 clusters
Clusters
Marker genes
E
Full data set cells
113 clusters
Clusters
Marker genes

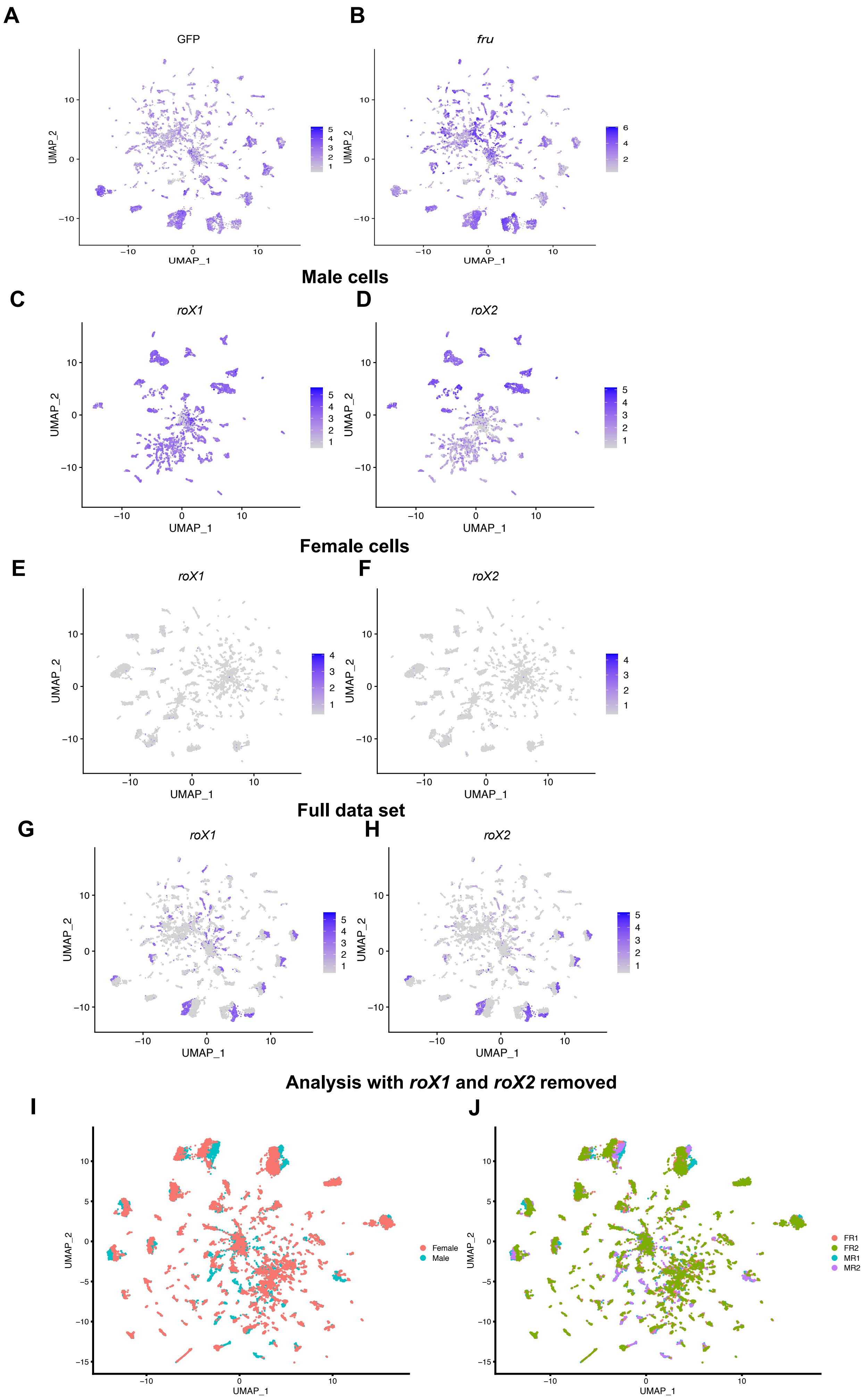
A
GFP
B
fru
UMAP_1
UMAP_2
Male cells
C
roX1
D
roX2
Female cells
E
roX1
F
roX2
Full data set
G
roX1
H
roX2
Analysis with roX1 and roX2 removed
I
Female
Male
J
FR1
FR2
MR1
MR2

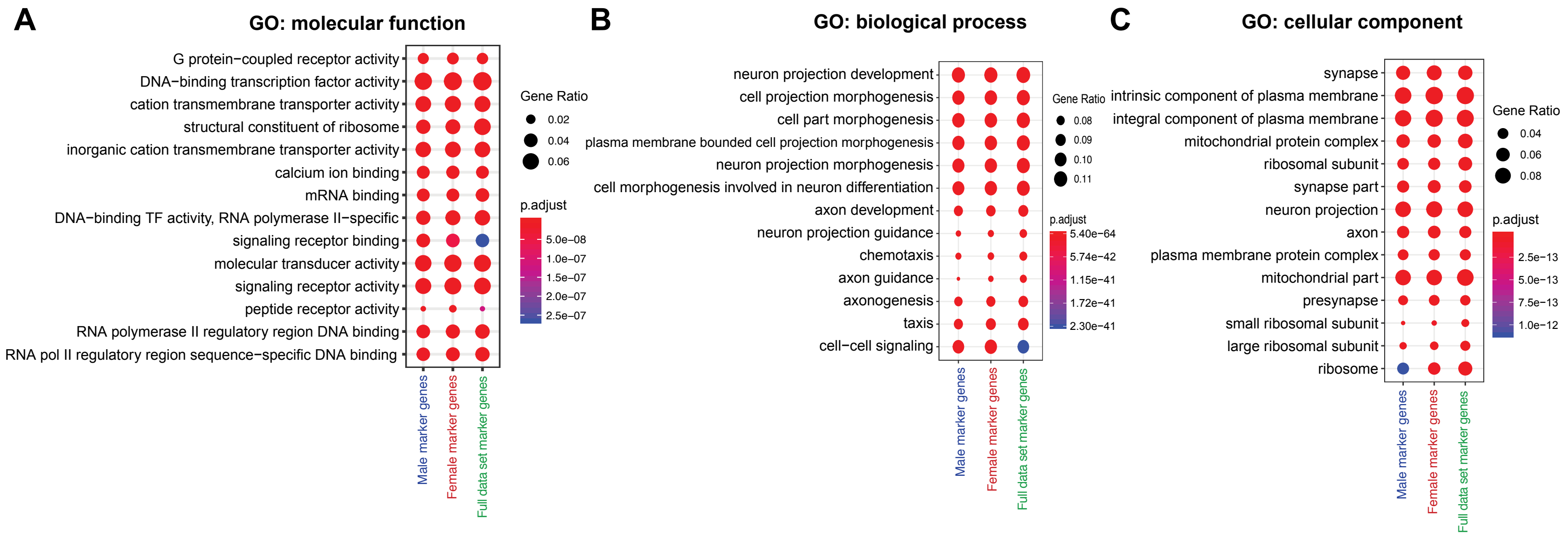
A
GO: molecular function
G protein−coupled receptor activity
DNA−binding transcription factor activity
cation transmembrane transporter activity
structural constituent of ribosome
inorganic cation transmembrane transporter activity
calcium ion binding
mRNA binding
DNA−binding TF activity, RNA polymerase II−specific
signaling receptor binding
molecular transducer activity
signaling receptor activity
peptide receptor activity
RNA polymerase II regulatory region DNA binding
RNA pol II regulatory region sequence−specific DNA binding
Male marker genes
Female marker genes
Full data set marker genes
Gene Ratio
0.02
0.04
0.06
p.adjust
5.0e−08
1.0e−07
1.5e−07
2.0e−07
2.5e−07
B
GO: biological process
neuron projection development
cell projection morphogenesis
cell part morphogenesis
plasma membrane bounded cell projection morphogenesis
neuron projection morphogenesis
cell morphogenesis involved in neuron differentiation
axon development
neuron projection guidance
chemotaxis
axon guidance
axonogenesis
taxis
cell−cell signaling
Male marker genes
Female marker genes
Full data set marker genes
Gene Ratio
0.08
0.09
0.10
0.11
p.adjust
5.40e−64
5.74e−42
1.15e−41
1.72e−41
2.30e−41
C
GO: cellular component
synapse
intrinsic component of plasma membrane
integral component of plasma membrane
mitochondrial protein complex
ribosomal subunit
synapse part
neuron projection
axon
plasma membrane protein complex
mitochondrial part
presynapse
small ribosomal subunit
large ribosomal subunit
ribosome
Male marker genes
Female marker genes
Full data set marker genes
Gene Ratio
0.04
0.06
0.08
p.adjust
2.5e−13
5.0e−13
7.5e−13
1.0e−12

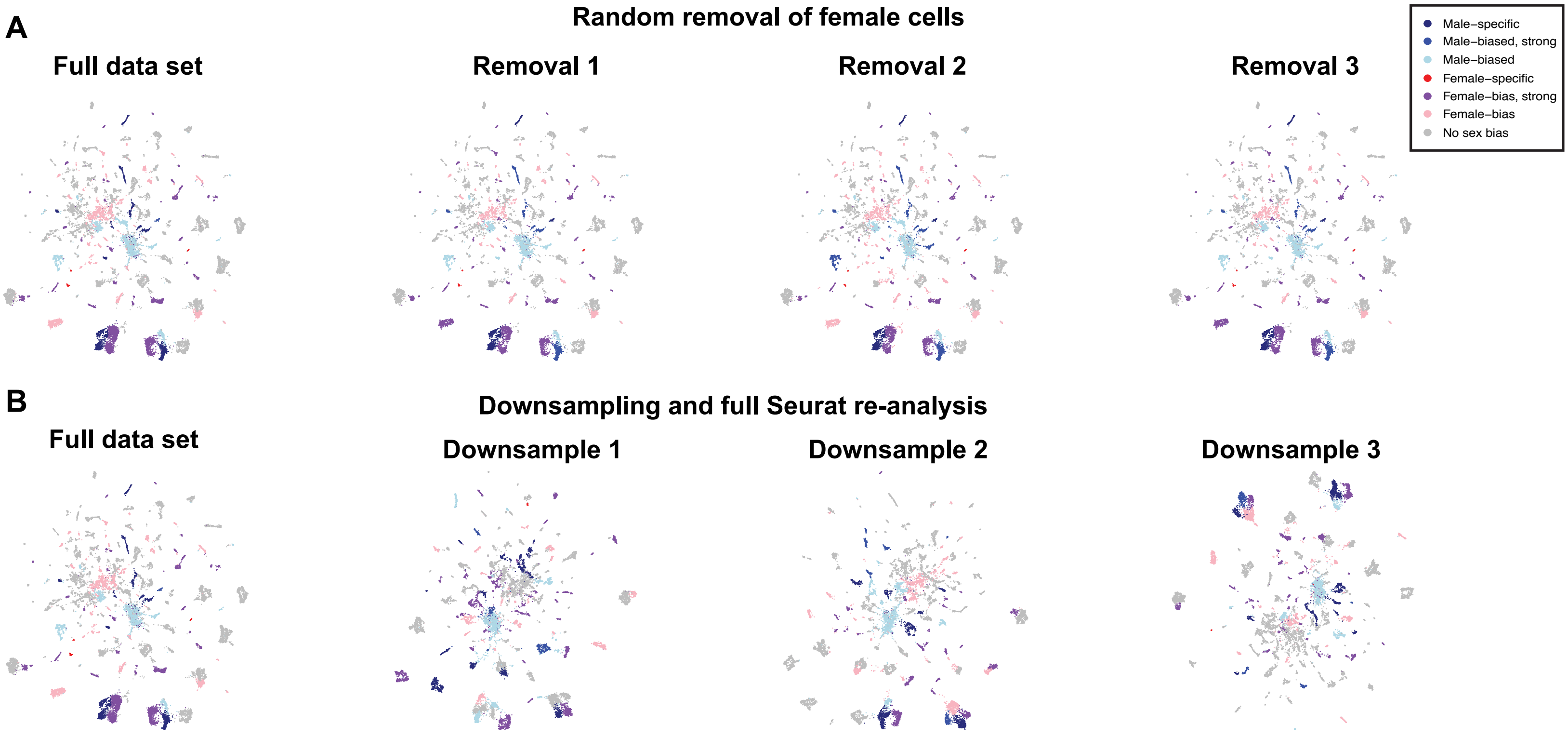

A
Random removal of female cells
Full data set
Removal 1
Removal 2
Removal 3
Male–specific
Male–biased, strong
Male–biased
Female–specific
Female–bias, strong
Female–bias
No sex bias
B
Downsampling and full Seurat re-analysis
Full data set
Downsample 1
Downsample 2
Downsample 3

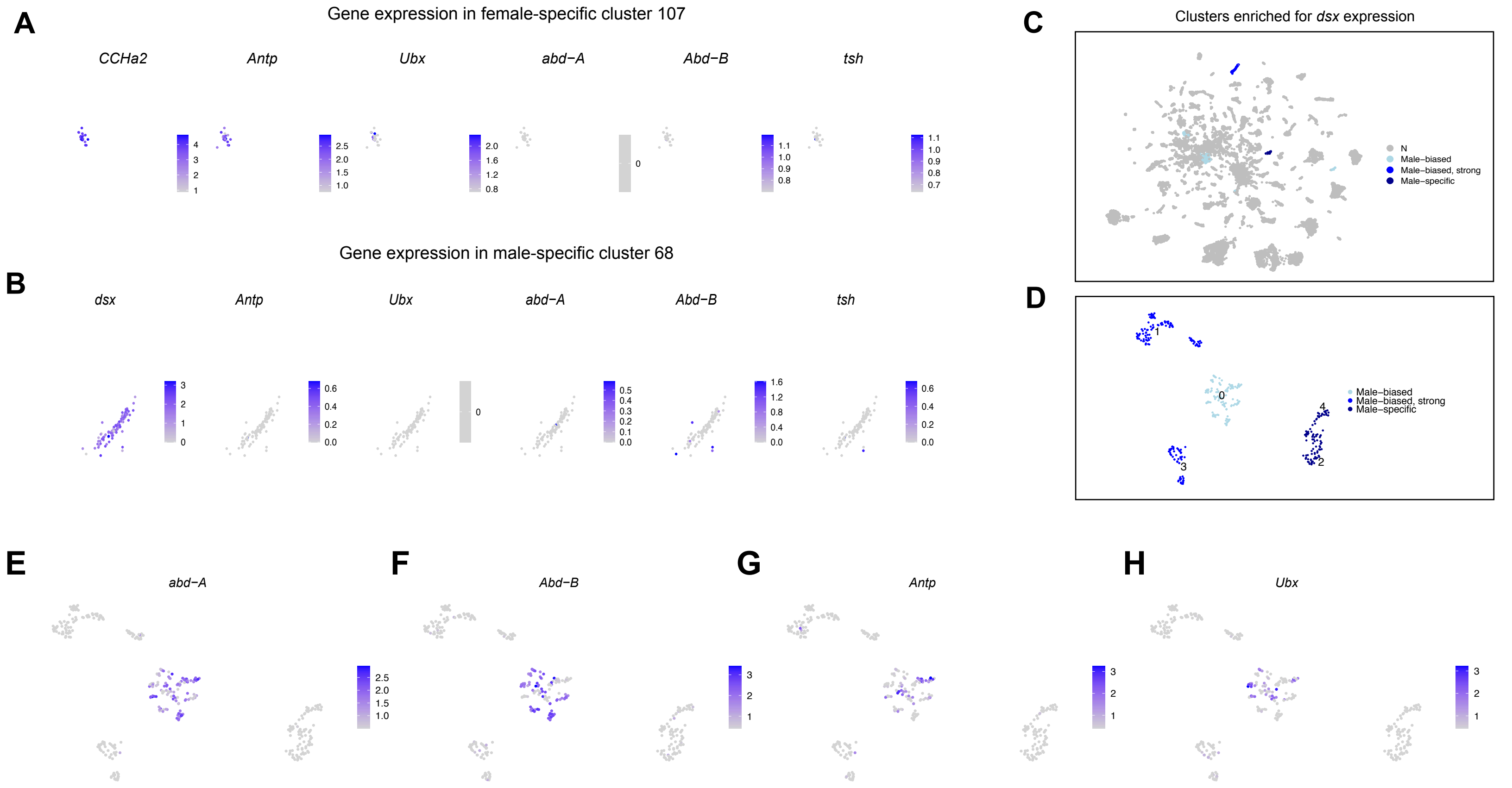
A
Gene expression in female-specific cluster 107
CCHa2
Antp
Ubx
abd−A
Abd−B
tsh
B
Gene expression in male-specific cluster 68
dsx
Antp
Ubx
abd−A
Abd−B
tsh
C
Clusters enriched for dsx expression
N
Male−biased
Male−biased, strong
Male−specific
D
Male−biased
Male−biased, strong
Male−specific
E
abd−A
F
Abd−B
G
Antp
H
Ubx

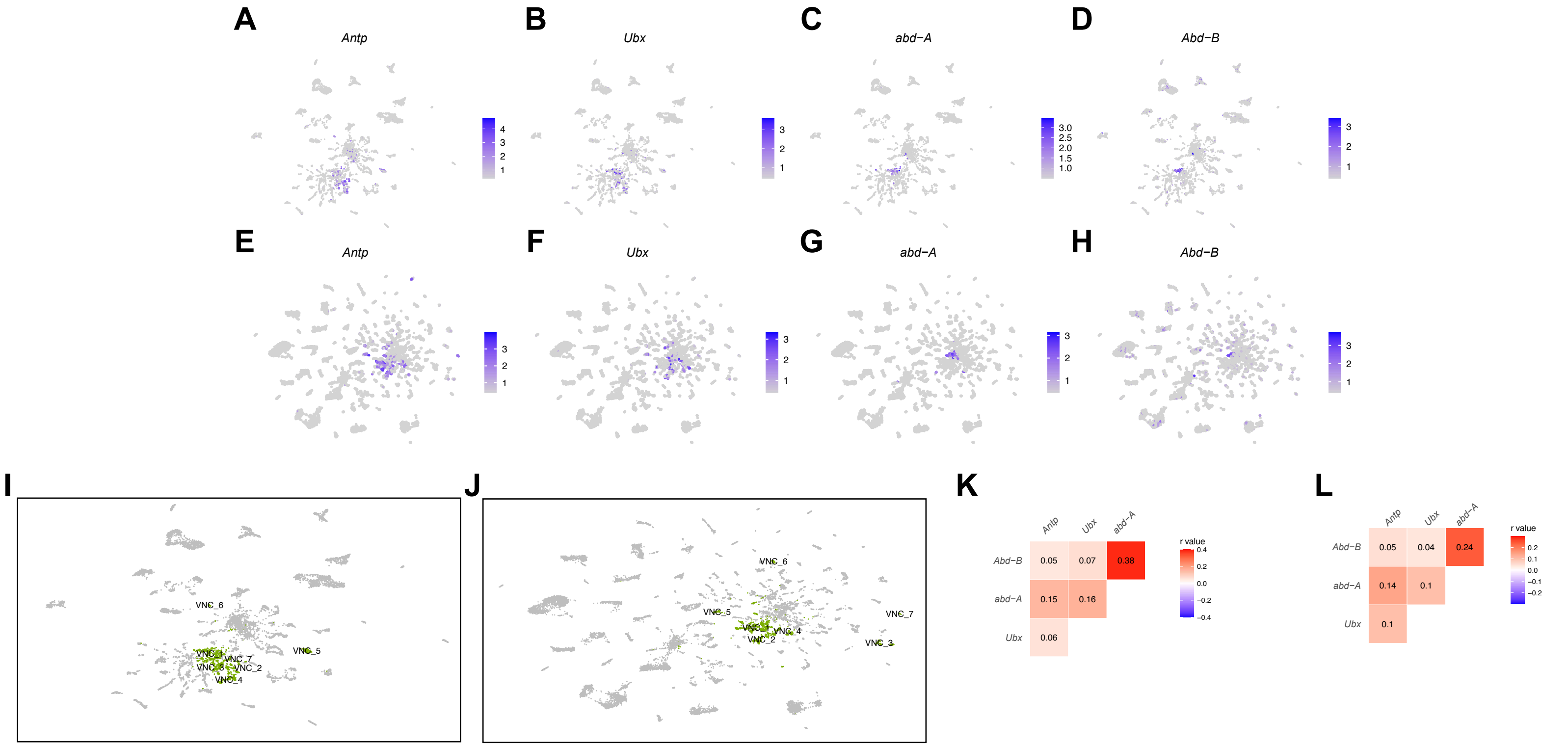
A
Antp
B
Ubx
C
abd−A
D
Abd−B
E
Antp
F
Ubx
G
abd−A
H
Abd−B
I
VNC_6
VNC_5
VNC_7
VNC_2
VNC_4
J
VNC_6
VNC_5
VNC_7
VNC_4
VNC_2
VNC_3
K
Antp
Ubx
abd−A
Abd−B
0.05
0.07
0.38
abd−A
0.15
0.16
Ubx
0.06
r value
L
Antp
Ubx
abd−A
Abd−B
0.05
0.04
0.24
abd−A
0.14
0.1
Ubx
0.1
r value

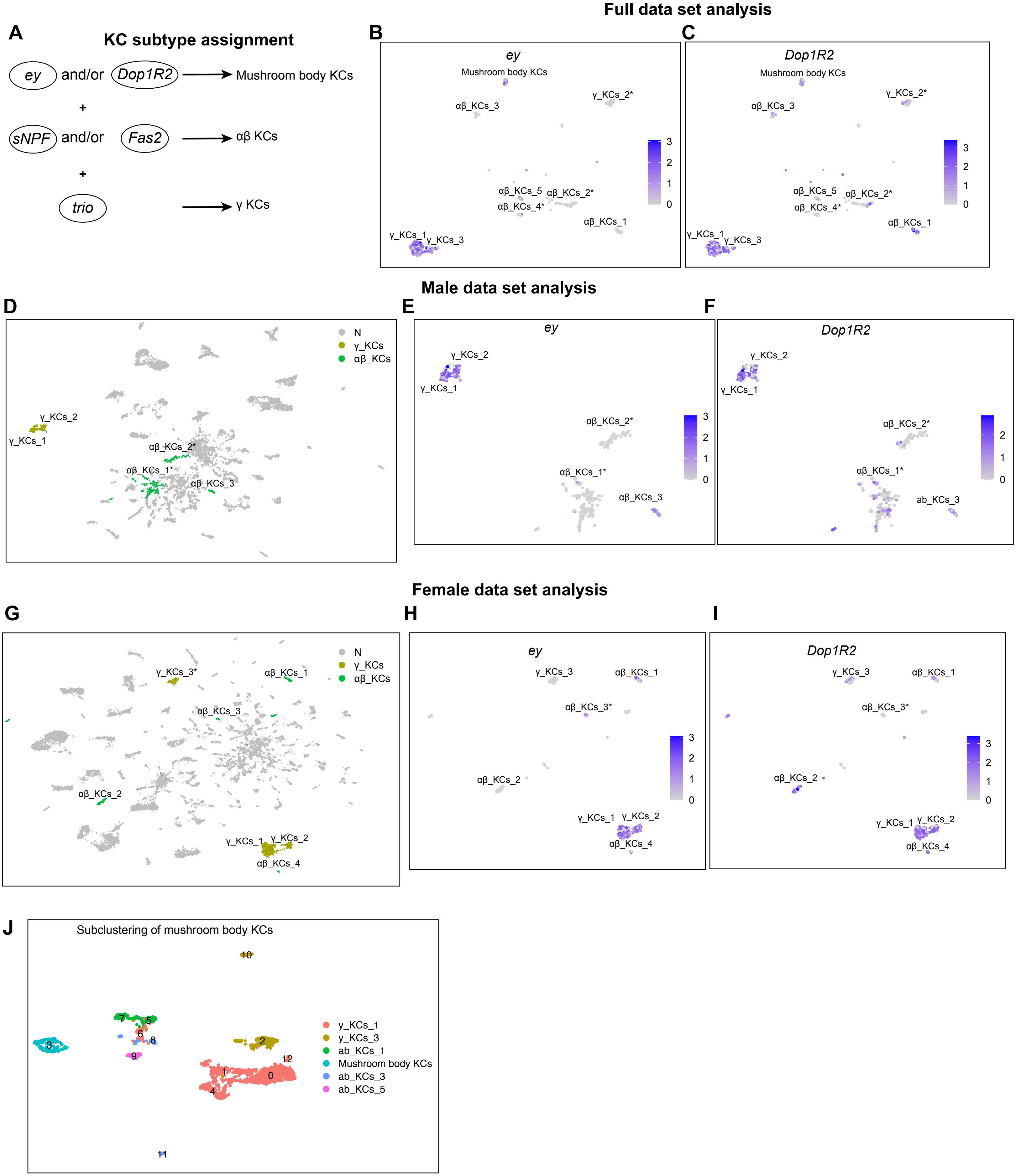

Full data set analysis
A
KC subtype assignment
ey and/or Dop1R2 → Mushroom body KCs
+
sNPF and/or Fas2 → αβ KCs
+
trio → γ KCs
B
ey
C
Dop1R2
Mushroom body KCs
γ_KCs_2*
αβ_KCs_3
αβ_KCs_5
αβ_KCs_2*
αβ_KCs_4*
αβ_KCs_1
γ_KCs_1
γ_KCs_3
Male data set analysis
D
E
ey
F
Dop1R2
N
γ_KCs
αβ_KCs
γ_KCs_2
γ_KCs_1
αβ_KCs_2*
αβ_KCs_1*
αβ_KCs_3
ab_KCs_3
Female data set analysis
G
H
ey
I
Dop1R2
γ_KCs_3*
γ_KCs_3
αβ_KCs_1
αβ_KCs_3
αβ_KCs_3*
αβ_KCs_2
γ_KCs_1
γ_KCs_2
αβ_KCs_4
J
Subclustering of mushroom body KCs
γ_KCs_1
γ_KCs_3
ab_KCs_1
Mushroom body KCs
ab_KCs_3
ab_KCs_5

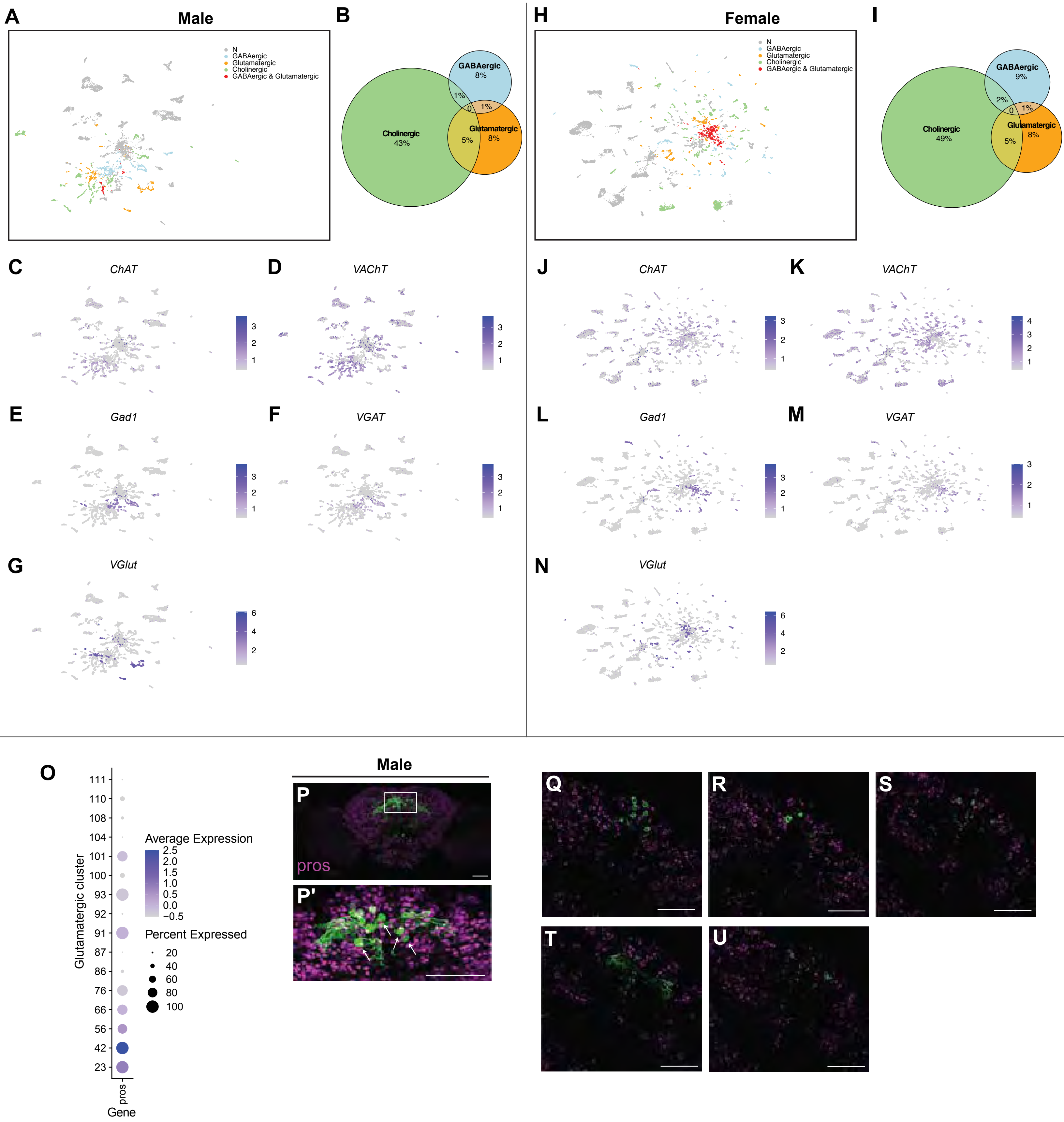
A
Male
N
GABAergic
Glutamatergic
Cholinergic
GABAergic & Glutamatergic
B
GABAergic
8%
1%
0
1%
Cholinergic
43%
Glutamatergic
5%
8%
C
ChAT
D
VAChT
E
Gad1
F
VGAT
G
VGlut
H
Female
N
GABAergic
Glutamatergic
Cholinergic
GABAergic & Glutamatergic
I
GABAergic
9%
2%
0
1%
Cholinergic
49%
Glutamatergic
5%
8%
J
ChAT
K
VAChT
L
Gad1
M
VGAT
N
VGlut
O
Glutamatergic cluster
111
110
108
104
101
100
93
92
91
87
86
76
66
56
42
23
Average Expression
2.5
2.0
1.5
1.0
0.5
0.0
−0.5
Percent Expressed
20
40
60
80
100
pros
Gene
Male
P
pros
P'
Q
R
S
T
U

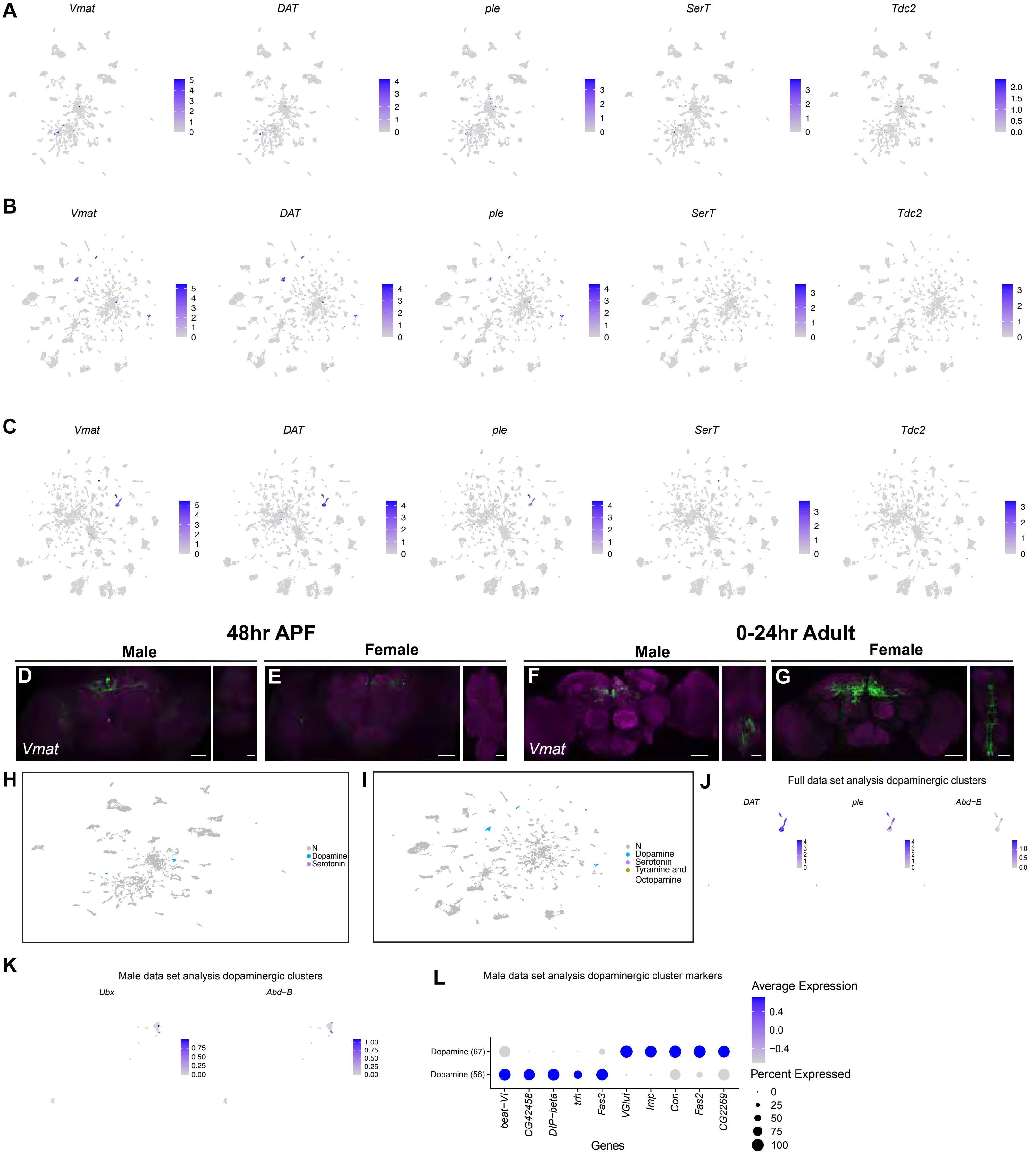
A
Vmat
DAT
ple
SerT
Tdc2
B
C
48hr APF
Male
Female
0-24hr Adult
D
E
F
G
Vmat
H
N
Dopamine
Serotonin
I
Tyramine and Octopamine
J
Full data set analysis dopaminergic clusters
Abd−B
K
Male data set analysis dopaminergic clusters
Ubx
L
Male data set analysis dopaminergic cluster markers
Average Expression
Percent Expressed
Dopamine (67)
Dopamine (56)
beat−VI
CG42458
DIP−beta
trh
Fas3
VGlut
Imp
Con
Fas2
CG2269
Genes

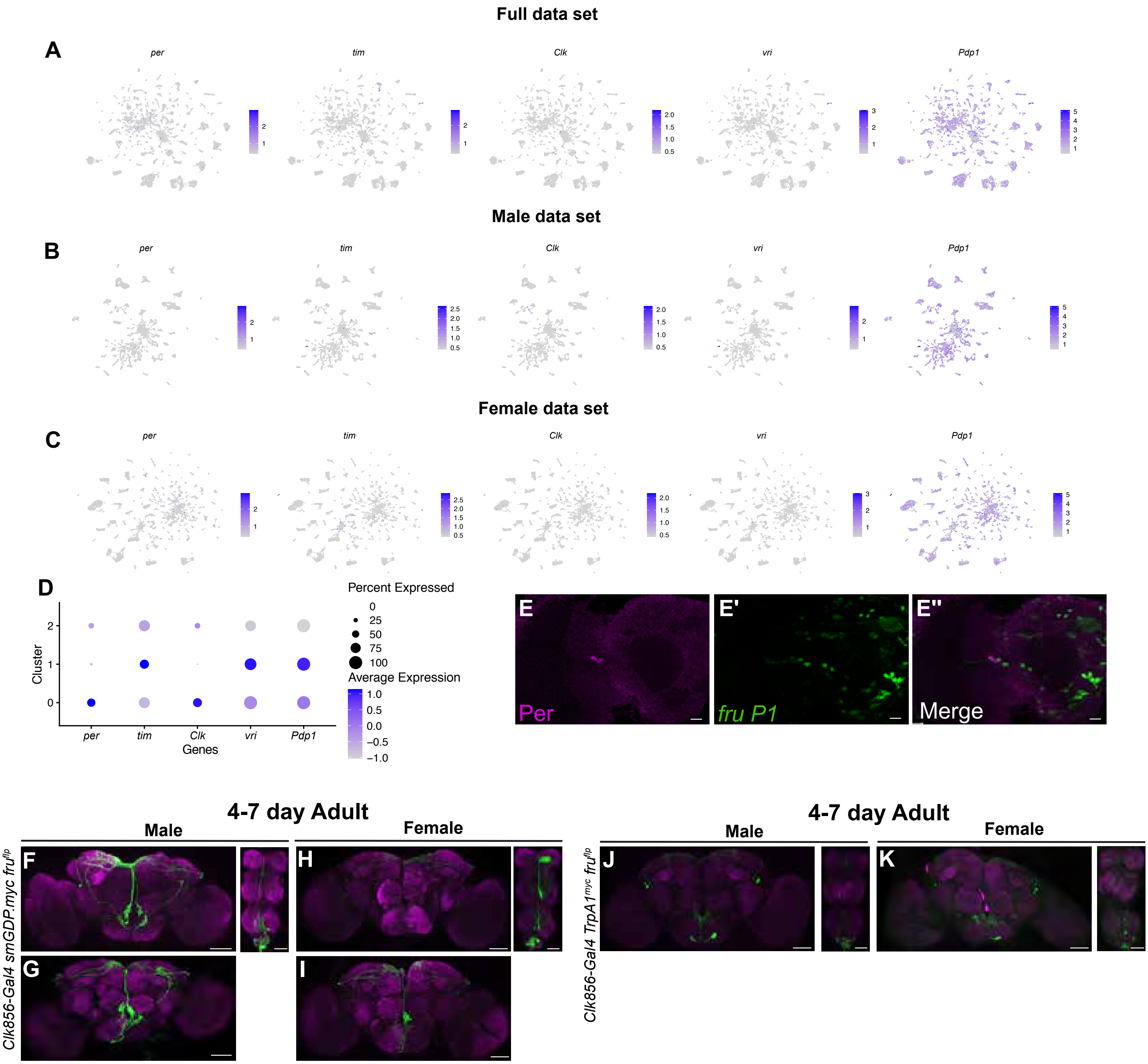

Full data set
A
per
tim
Clk
vri
Pdp1
Male data set
B
Female data set
C
D
Percent Expressed
0
25
50
75
100
Average Expression
1.0
0.5
0.0
−0.5
−1.0
Cluster
Genes
E
E'
E''
Per
fru P1
Merge
4-7 day Adult
Male
Female
F
G
H
I
Clk856-Gal4 smGDP.myc fruflp
J
K
Clk856-Gal4 TrpA1myc fruflp

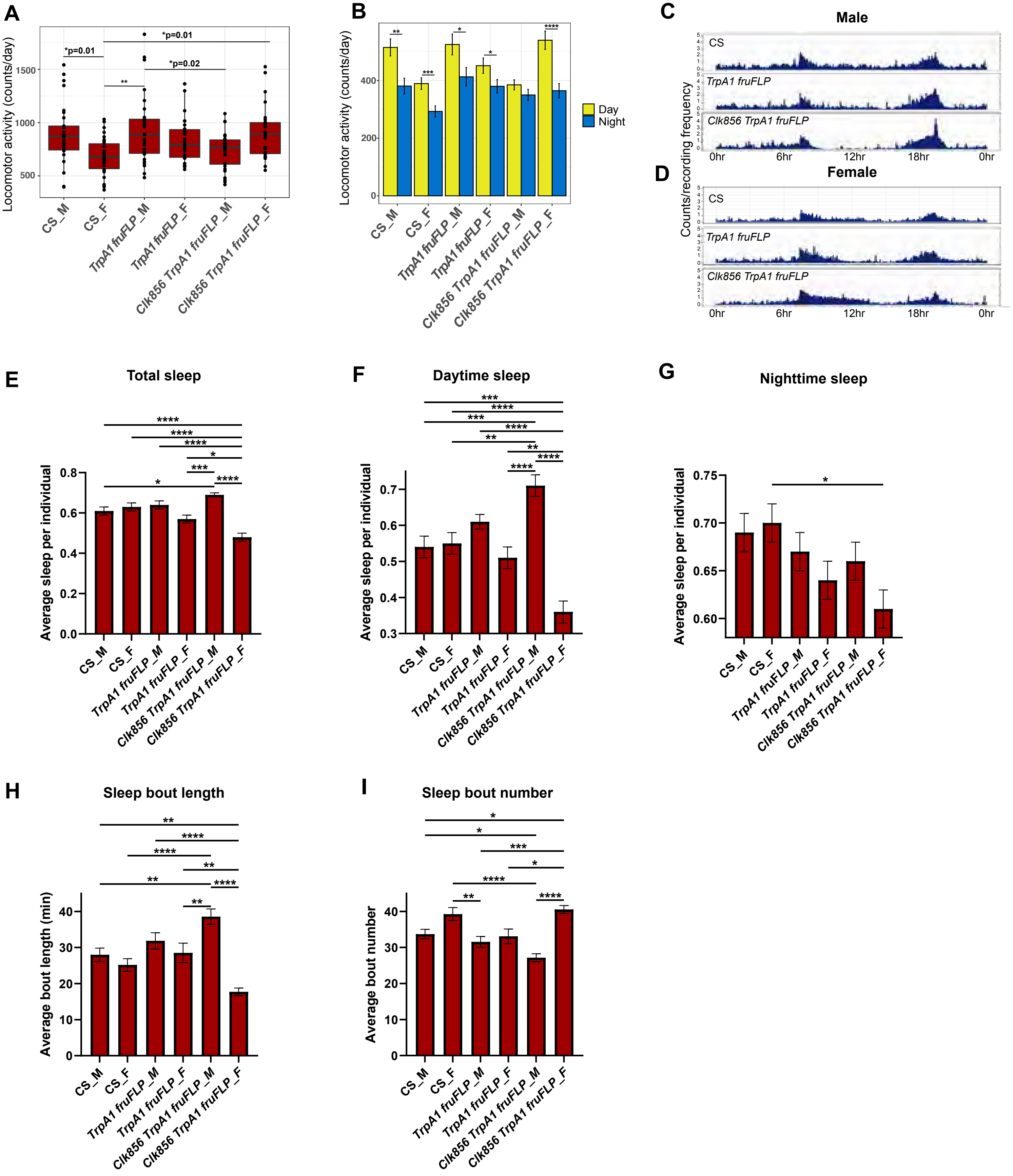
A
Locomotor activity (counts/day)
*p=0.01
*p=0.01
**
*p=0.02
CS_M
CS_F
TrpA1 fruFLP_M
TrpA1 fruFLP_F
Clk856 TrpA1 fruFLP_M
Clk856 TrpA1 fruFLP_F
B
Locomotor activity (counts/day)
Day
Night
C
Male
CS
TrpA1 fruFLP
Clk856 TrpA1 fruFLP
0hr
6hr
12hr
18hr
0hr
D
Female
Counts/recording frequency
E
Total sleep
Average sleep per individual
F
Daytime sleep
G
Nighttime sleep
H
Sleep bout length
Average bout length (min)
I
Sleep bout number
Average bout number

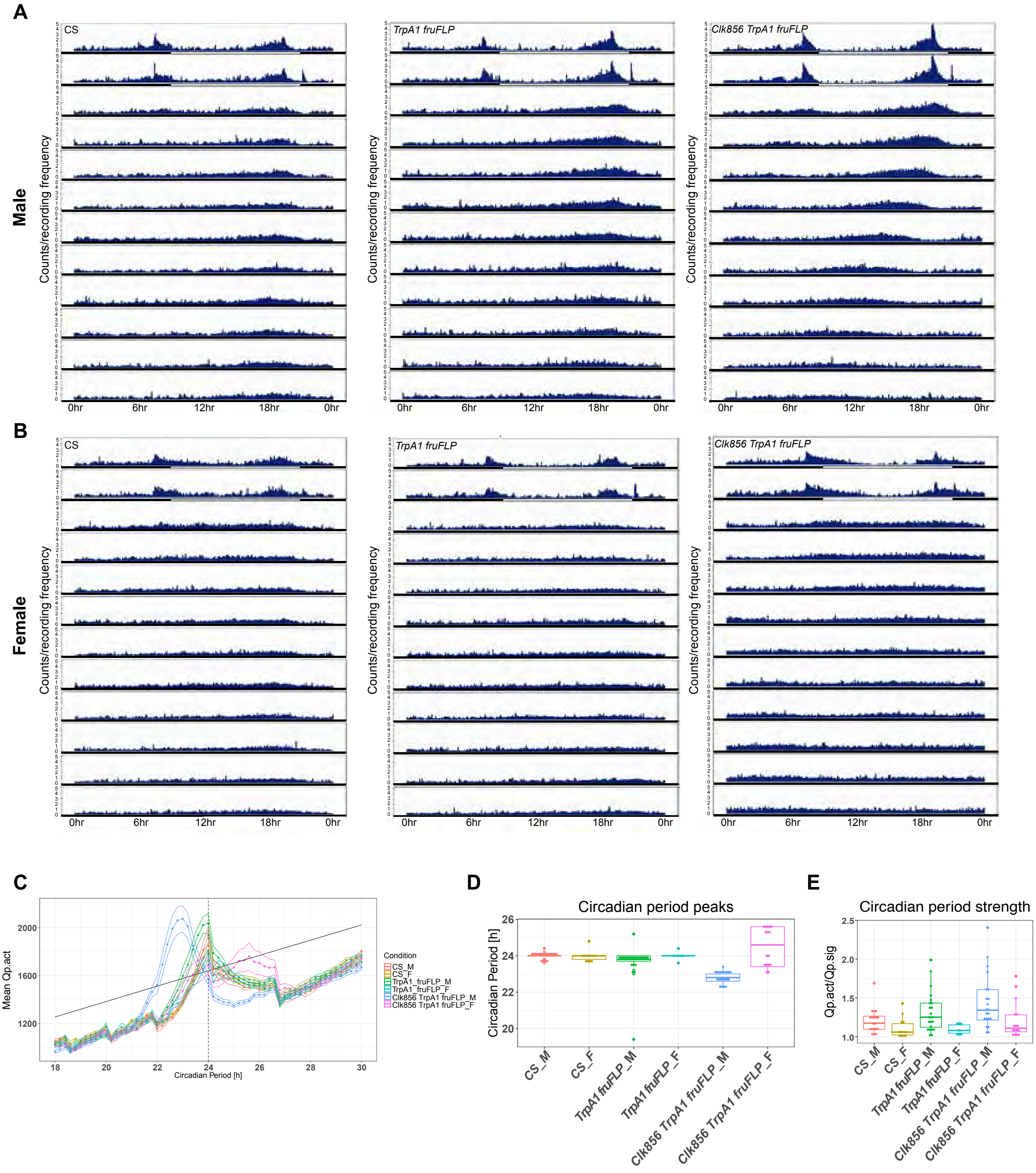
A
Male
Counts/recording frequency
CS
TrpA1 fruFLP
Clk856 TrpA1 fruFLP
0hr
6hr
12hr
18hr
0hr
B
Female
Counts/recording frequency
CS
TrpA1 fruFLP
Clk856 TrpA1 fruFLP
0hr
6hr
12hr
18hr
0hr
C
Mean Qp.act
Circadian Period [h]
Condition
CS_M
CS_F
TrpA1_fruFLP_M
TrpA1_fruFLP_F
Clk856 TrpA1 fruFLP_M
Clk856 TrpA1 fruFLP_F
D
Circadian period peaks
Circadian Period [h]
CS_M
CS_F
TrpA1 fruFLP_M
TrpA1 fruFLP_F
Clk856 TrpA1 fruFLP_M
Clk856 TrpA1 fruFLP_F
E
Circadian period strength
Qp.act/Qp.sig
CS_M
CS_F
TrpA1 fruFLP_M
TrpA1 fruFLP_F
Clk856 TrpA1 fruFLP_M
Clk856 TrpA1 fruFLP_F

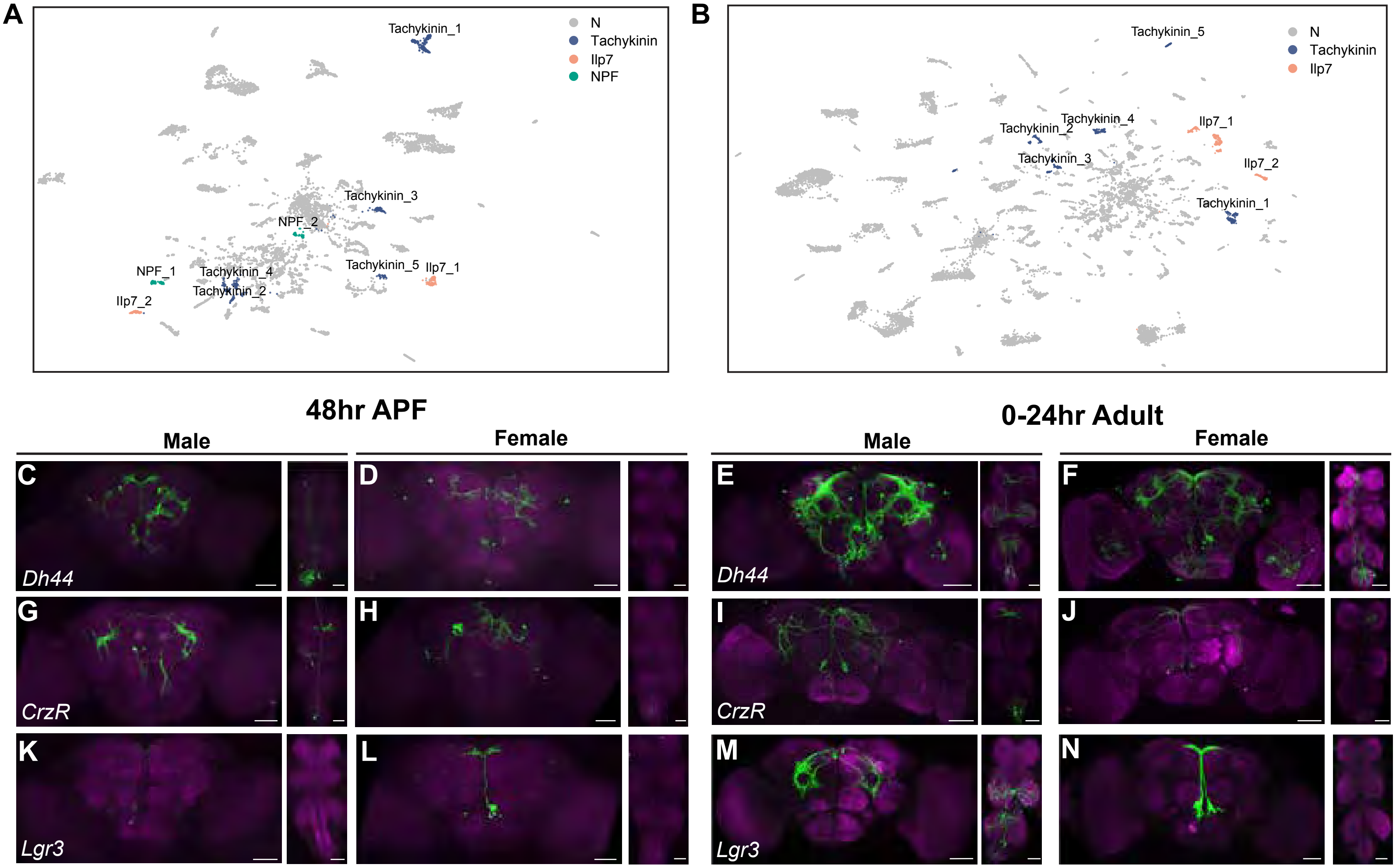
A
B
N
Tachykinin
Ilp7
NPF
Tachykinin_1
Tachykinin_2
Tachykinin_3
Tachykinin_4
Tachykinin_5
NPF_1
NPF_2
Ilp7_1
Ilp7_2
48hr APF
0-24hr Adult
Male
Female
C
D
E
F
G
H
I
J
K
L
M
N
Dh44
CrzR
Lgr3

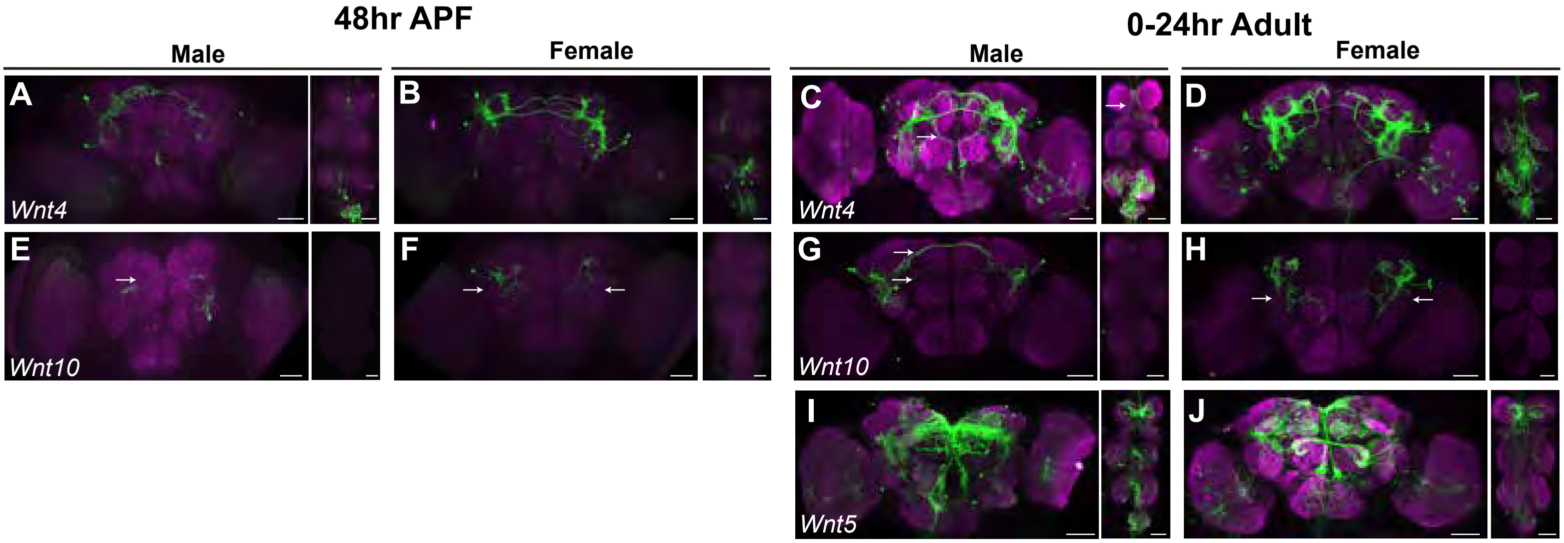
48hr APF
0-24hr Adult
Male
Female
Male
Female
A
B
C
D
Wnt4
Wnt4
E
F
G
H
Wnt10
Wnt10
I
J
Wnt5

A

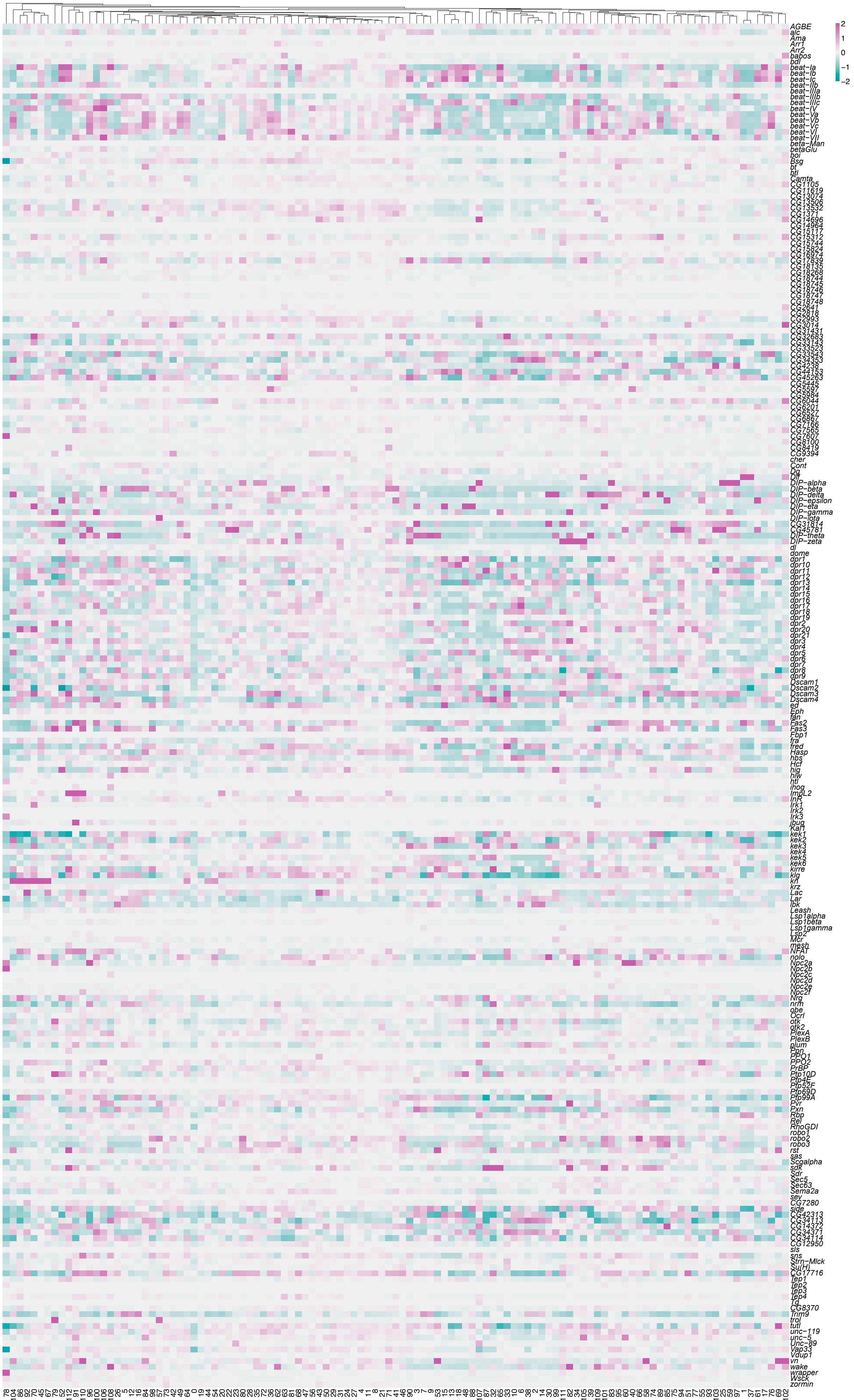
AGBE
alc
Ama
Arr1
Arr2
babos
bdl
beat−Ia
beat−Ib
beat−Ic
beat−IIb
beat−IIIa
beat−IIIb
beat−IIIc
beat−IV
beat−Va
beat−Vb
beat−Vc
beat−VI
beat−VII
beta−Man
betaGlu
boi
Bsg
bt
btl
Camta
CG1105
CG11619
CG13074
CG13506
CG13532
CG1371
CG14696
CG14964
CG15117
CG15312
CG15744
CG15824
CG16974
CG17839
CG18135
CG18268
CG18744
CG18745
CG18746
CG18747
CG18748
CG2641
CG2818
CG2993
CG3014
CG31431
CG32683
CG33143
CG33523
CG33543
CG34353
CG4238
CG44153
CG45263
CG5445
CG5597
CG5984
CG6044
CG6201
CG6527
CG6867
CG7166
CG7565
CG7607
CG8100
CG8419
CG9394
cher
Cont
Dg
Dif
DIP−alpha
DIP−beta
DIP−delta
DIP−epsilon
DIP−eta
DIP−gamma
DIP−iota
CG31814
CG45781
DIP−theta
DIP−zeta
dl
dome
dpr1
dpr10
dpr11
dpr12
dpr13
dpr14
dpr15
dpr16
dpr17
dpr18
dpr19
dpr2
dpr20
dpr21
dpr3
dpr4
dpr5
dpr6
dpr7
dpr8
dpr9
Dscam1
Dscam2
Dscam3
Dscam4
ed
Eph
fan
Fas2
Fas3
Fbp1
fra
fred
Hasp
hbs
Hcf
hig
hiw
htl
ihog
ImpL2
InR
Irk1
Irk2
Irk3
jbug
Kal1
kek1
kek2
kek3
kek4
kek5
kek6
kirre
klg
kn
krz
Lac
Lar
lbk
Leash
Lsp1alpha
Lsp1beta
Lsp1gamma
Lsp2
Mcr
mesh
NFAT
nolo
Npc2a
Npc2b
Npc2c
Npc2d
Npc2e
Npc2f
Nrg
nrm
obe
Ocrl
otk
otk2
PlexA
PlexB
plum
Ppn
PPO1
PPO2
PrBP
Ptp10D
Ptp4E
Ptp52F
Ptp69D
Ptp99A
Pvr
Pxn
Rbp
Rel
RhoGDI
robo1
robo2
robo3
rst
sas
Scgalpha
sdk
Sdr
Sec5
Sec63
Sema2a
sev
CG7280
side
CG42313
CG34113
CG14372
CG34371
CG34114
CG12950
sls
sns
Strn−Mlck
Su(H)
CG17716
Tep1
Tep2
Tep3
Tep4
Tg
CG8370
Trim9
trol
tutl
unc−119
unc−5
Unc−89
Vap33
Vdup1
vn
wake
wrapper
Wsck
zormin
2
1
0
−1
−2

A

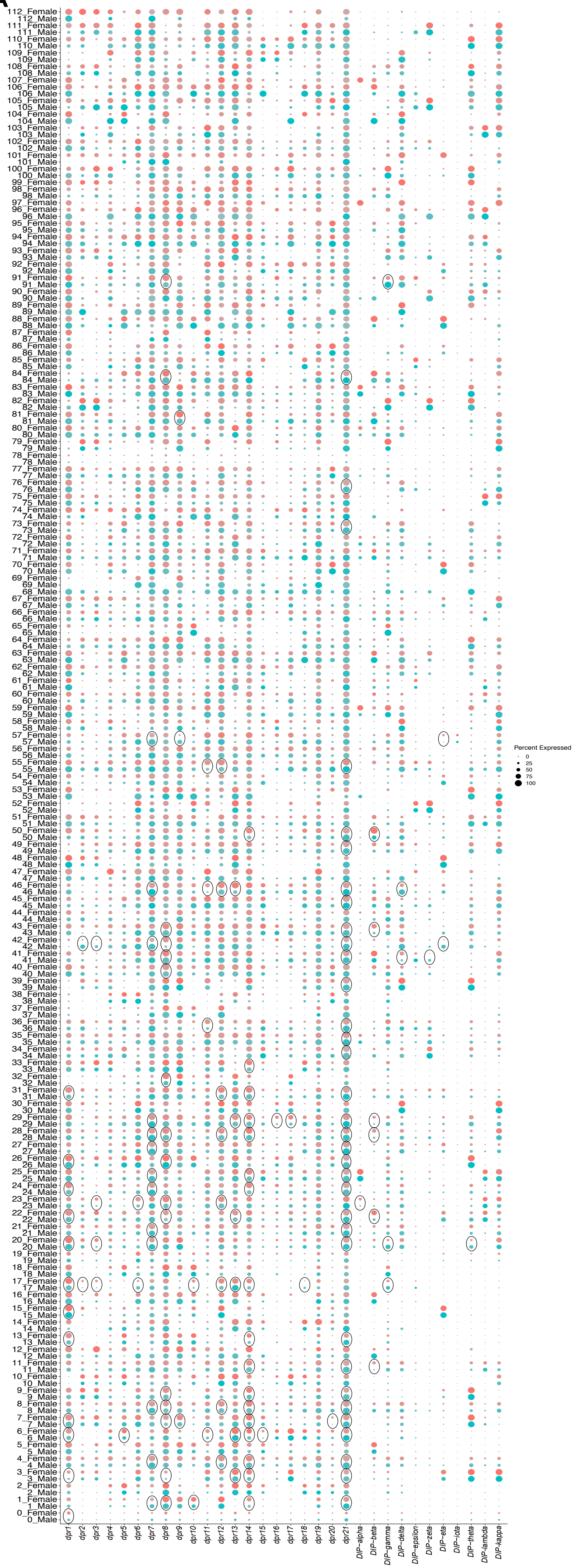
112_Female
112_Male
111_Female
111_Male
110_Female
110_Male
109_Female
109_Male
108_Female
108_Male
107_Female
107_Male
106_Female
106_Male
105_Female
105_Male
104_Female
104_Male
103_Female
103_Male
102_Female
102_Male
101_Female
101_Male
100_Female
100_Male
99_Female
99_Male
98_Female
98_Male
97_Female
97_Male
96_Female
96_Male
95_Female
95_Male
94_Female
94_Male
93_Female
93_Male
92_Female
92_Male
91_Female
91_Male
90_Female
90_Male
89_Female
89_Male
88_Female
88_Male
87_Female
87_Male
86_Female
86_Male
85_Female
85_Male
84_Female
84_Male
83_Female
83_Male
82_Female
82_Male
81_Female
81_Male
80_Female
80_Male
79_Female
79_Male
78_Female
78_Male
77_Female
77_Male
76_Female
76_Male
75_Female
75_Male
74_Female
74_Male
73_Female
73_Male
72_Female
72_Male
71_Female
71_Male
70_Female
70_Male
69_Female
69_Male
68_Female
68_Male
67_Female
67_Male
66_Female
66_Male
65_Female
65_Male
64_Female
64_Male
63_Female
63_Male
62_Female
62_Male
61_Female
61_Male
60_Female
60_Male
59_Female
59_Male
58_Female
58_Male
57_Female
57_Male
56_Female
56_Male
55_Female
55_Male
54_Female
54_Male
53_Female
53_Male
52_Female
52_Male
51_Female
51_Male
50_Female
50_Male
49_Female
49_Male
48_Female
48_Male
47_Female
47_Male
46_Female
46_Male
45_Female
45_Male
44_Female
44_Male
43_Female
43_Male
42_Female
42_Male
41_Female
41_Male
40_Female
40_Male
39_Female
39_Male
38_Female
38_Male
37_Female
37_Male
36_Female
36_Male
35_Female
35_Male
34_Female
34_Male
33_Female
33_Male
32_Female
32_Male
31_Female
31_Male
30_Female
30_Male
29_Female
29_Male
28_Female
28_Male
27_Female
27_Male
26_Female
26_Male
25_Female
25_Male
24_Female
24_Male
23_Female
23_Male
22_Female
22_Male
21_Female
21_Male
20_Female
20_Male
19_Female
19_Male
18_Female
18_Male
17_Female
17_Male
16_Female
16_Male
15_Female
15_Male
14_Female
14_Male
13_Female
13_Male
12_Female
12_Male
11_Female
11_Male
10_Female
10_Male
9_Female
9_Male
8_Female
8_Male
7_Female
7_Male
6_Female
6_Male
5_Female
5_Male
4_Female
4_Male
3_Female
3_Male
2_Female
2_Male
1_Female
1_Male
0_Female
0_Male
dpr1
dpr2
dpr3
dpr4
dpr5
dpr6
dpr7
dpr8
dpr9
dpr10
dpr11
dpr12
dpr13
dpr14
dpr15
dpr16
dpr17
dpr18
dpr19
dpr20
dpr21
DIP−alpha
DIP−beta
DIP−gamma
DIP−delta
DIP−epsilon
DIP−zeta
DIP−eta
DIP−iota
DIP−theta
DIP−lambda
DIP−kappa
Percent Expressed
0
25
50
75
100

A

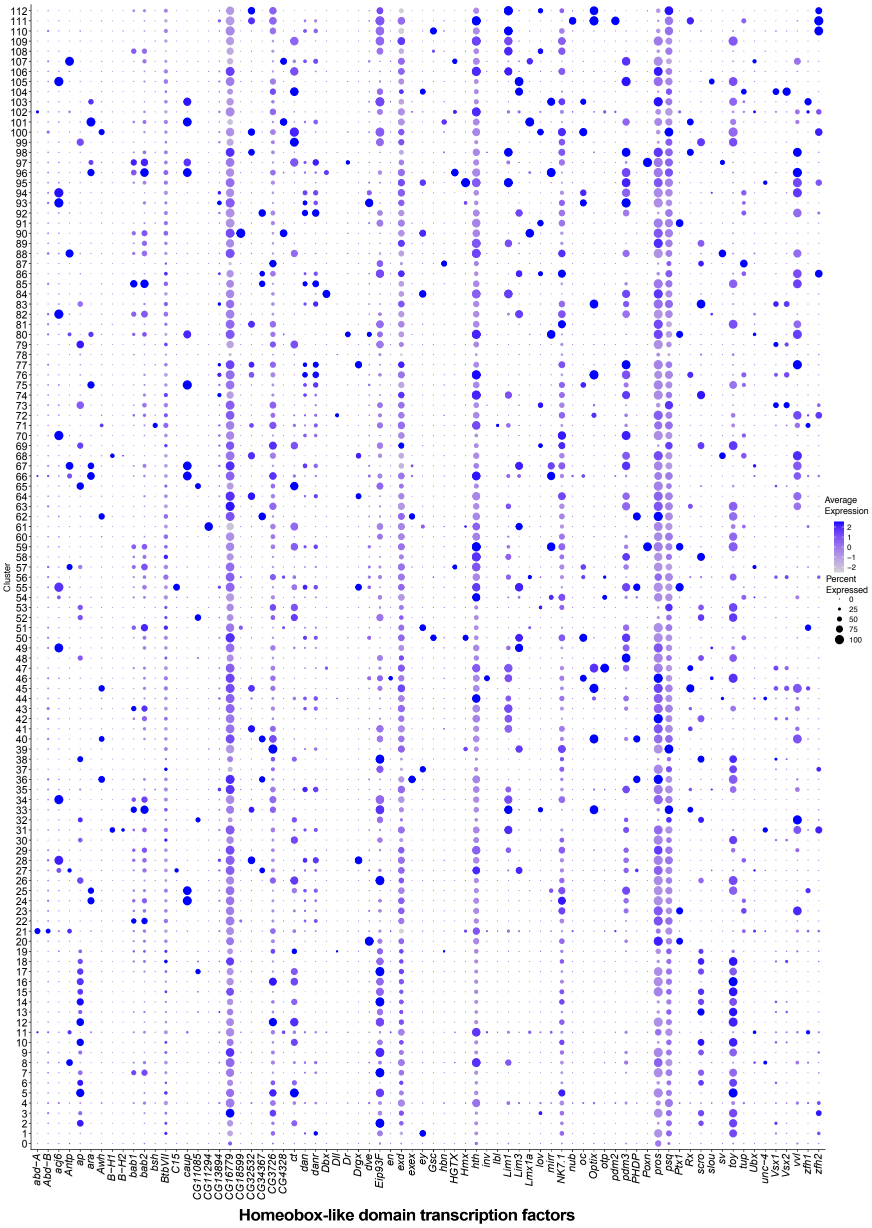

Cluster
0 1 2 3 4 5 6 7 8 9 10 11 12 13 14 15 16 17 18 19 20 21 22 23 24 25 26 27 28 29 30 31 32 33 34 35 36 37 38 39 40 41 42 43 44 45 46 47 48 49 50 51 52 53 54 55 56 57 58 59 60 61 62 63 64 65 66 67 68 69 70 71 72 73 74 75 76 77 78 79 80 81 82 83 84 85 86 87 88 89 90 91 92 93 94 95 96 97 98 99 100 101 102 103 104 105 106 107 108 109 110 111 112
abd−A Abd−B acj6 Antp ap ara Awh B−H1 B−H2 bab1 bab2 bsh BtbVII C15 caup CG11085 CG11294 CG13894 CG16779 CG18599 CG32532 CG34367 CG3726 CG4328 ct dan danr Dbx Dll Dr Drgx dve Eip93F en exd exex ey Gsc hbn HGTX Hmx hth inv lbl Lim1 Lim3 Lmx1a lov mirr NK7.1 nub oc Optix otp pdm2 pdm3 PHDP Poxn pros psq Ptx1 Rx scro slou sv toy tup Ubx unc−4 Vsx1 Vsx2 vvl zfh1 zfh2
Homeobox-like domain transcription factors
Average Expression
2
1
0
−1
−2
Percent Expressed
0
25
50
75
100

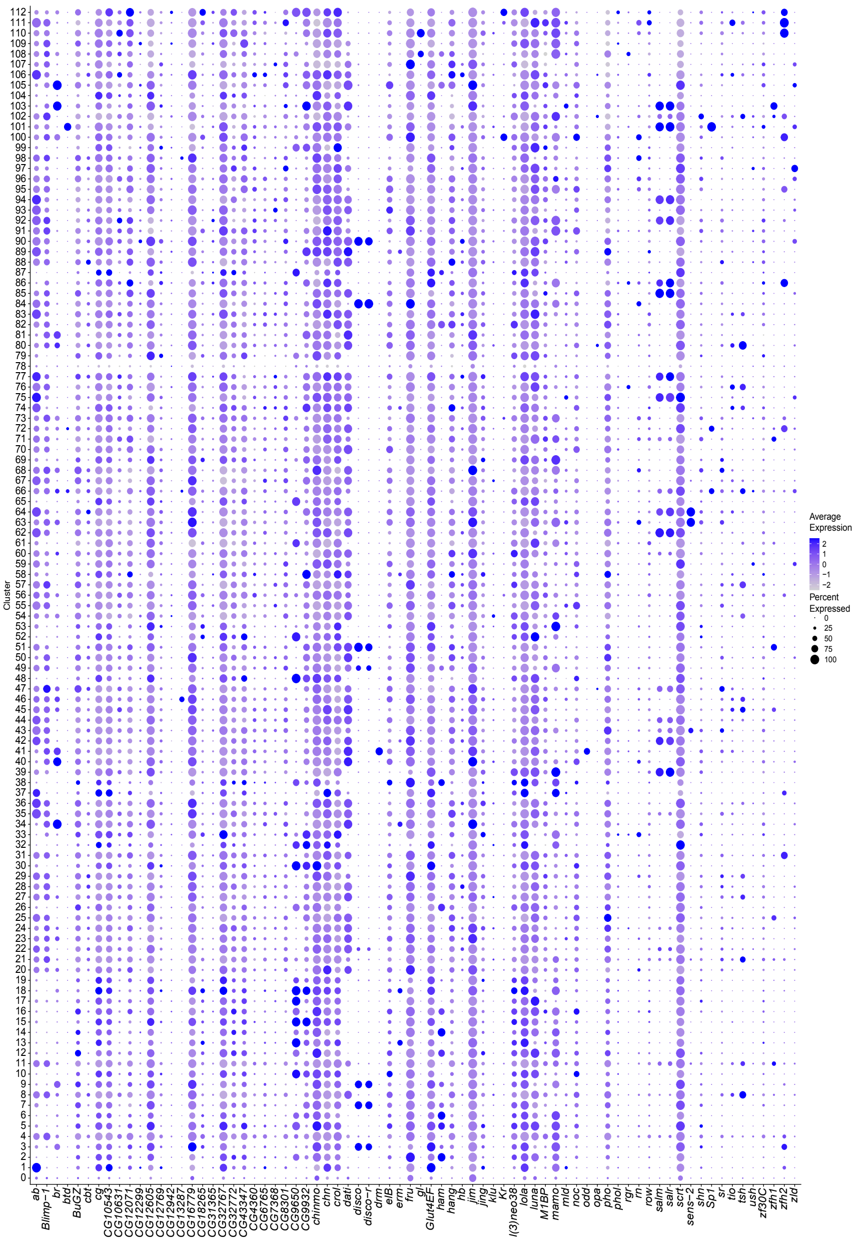


**Zinc finger C2H2 superfamily TFs**

ab Blimp-1 br btd BuGZ cbt cg CG10543 CG10631 CG12071 CG12299 CG12605 CG12769 CG12942 CG13287 CG16779 CG18265 CG31365 CG32767 CG32772 CG43347 CG4360 CG6765 CG7368 CG8301 CG9650 CG9932 chinmo chn crol dati disco disco-r drm elB erm fru gl Glut4EF ham hang hb jim jing klu Kr l(3)neo38 lola luna M1BP mamo mld noc odd opa pho phol rgr rn row salm salr scrt sens-2 shn Sp1 sr tio tsh ush zf30C zfh1 zfh2 zld

**A**

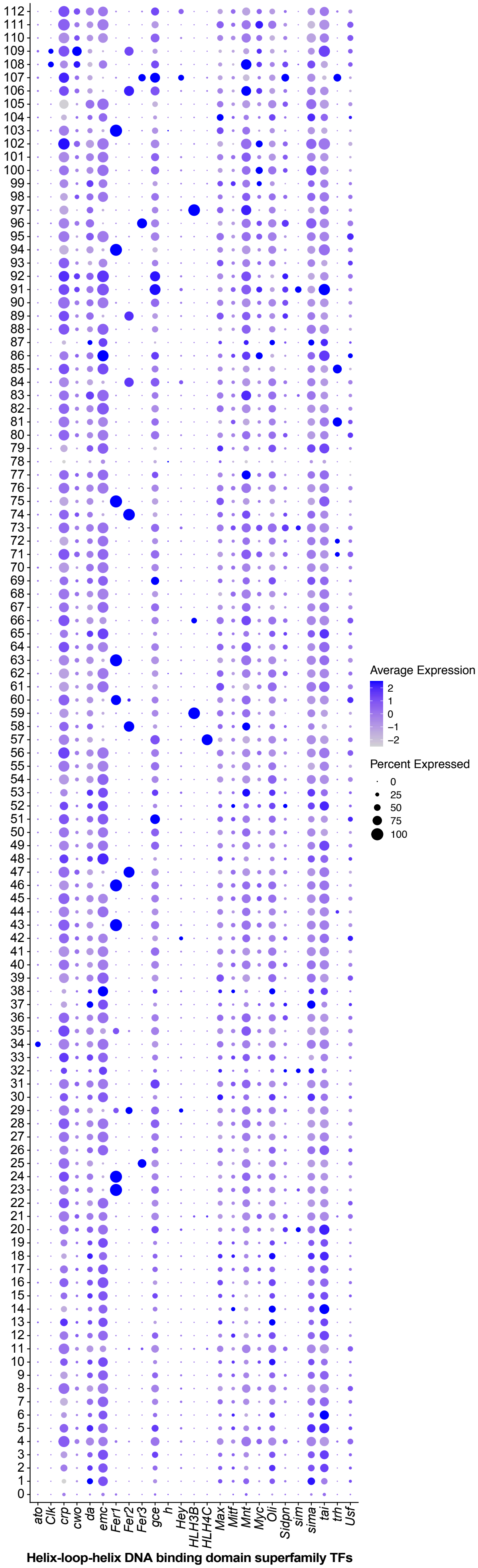

Average Expression
2
1
0
−1
−2
Percent Expressed
0
25
50
75
100
ato
Clk
crp
cwo
da
emc
Fer1
Fer2
Fer3
gce
h
Hey
HLH3B
HLH4C
Max
Mitf
Mnt
Myc
Oli
Sidpn
sim
sima
tai
trh
Usf
Helix-loop-helix DNA binding domain superfamily TFs
